# The biosynthesis of phospholipids is linked to the cell cycle in a model eukaryote

**DOI:** 10.1101/2021.02.19.431944

**Authors:** Milada Vítová, Vojtěch Lanta, Mária Čížková, Martin Jakubec, Frode Rise, Øyvind Halskau, Kateřina Bišová, Samuel Furse

## Abstract

The structural challenges faced by eukaryotic cells through the cell cycle are key for understanding cell viability and proliferation. In this study, we tested the hypothesis that the biosynthesis of structural lipids is linked to the cell cycle. If true, this would suggest that the cell’s structure would form part the control of the cell cycle. Lipidomics (^31^P NMR and MS), proteomics (Western immunoblotting) and transcriptomics (RT-qPCR) techniques were used to profile the lipid fraction and characterise aspects of its metabolism at seven stages of the cell cycle of the model eukaryote, *Desmodesmus quadricauda*. We found considerable, transient increases in the abundance of phosphatidylethanolamine during the G_1_ phase (+35%, ethanolamine phosphate cytidylyltransferase increased 2·5×) and phosphatidylglycerol over the G_1_/pre-replication phase boundary (+100%, phosphatidylglycerol synthase increased 22×). The relative abundance of phosphatidylcholine fell by ~35% during the G_1_. *N*-Methyl transferases for the conversion of phosphatidylethanolamine into phosphatidylcholine were not found in the *de novo* transcriptome profile, though a choline phosphate transferase was found, suggesting that the Kennedy pathway is the principal route for the synthesis of PC. The fatty acid profiles of the four most abundant lipids suggested that these lipids were not generally converted between one another. The relative abundance of both phosphatidylinositol and its synthase remained constant despite an eightfold increase in cell volume. We conclude that the biosynthesis of the three most abundant structural phospholipids is linked to the cell cycle in *D. quadricauda*.

## Introduction

The processes governing control of the cell cycle in eukaryotic organisms have been researched and characterised in considerable depth over the last half-century. This work has shown that checkpoints and the expression and degradation of cyclins and cyclin-dependent kinases are important in controlling progress through the cell cycle in fungi and metazoans (1–3). Similar mechanisms and homologous proteins were subsequently found in plants (4–6). However, successful completion of the cell cycle also presents a number of structural challenges. The plasma and compartment membranes must expand and undergo topological remodelling during the cell cycle, whilst maintain biochemical and barrier functions. The correct components of the daughter cells and their spatial arrangement must be organised. Finally, lipid membranes must be divided so that two or more viable daughter cells are produced. Interruption in the functions of the membrane, mis-timed membrane lysis or incorrect spatial distribution of cell components all represent perilous threats to cell survival that must be avoided for a cell to be viable.

The success of the cell cycle across countless species indicates that the factors that affect membrane behaviour, shape and size are under careful control through repeated cycles of cell division. This is also supported by recent reports about the structural integrity of cells through growth and division. Evidence for checkpoints that couple the structural integrity of the plasma membrane to DNA synthesis in eukaryotic cells is beginning to emerge (7). Control of division through cell size has also received attention in single-cell organisms such as yeast (8), and prokaryotes (9,10) but also in green algae (11,12). This work is particularly interesting in the light of evidence for the modulation of the composition of structural lipids in prokaryotes through their cell cycle (13,14). The changes in the topology of the cell envelope of prokaryotes and their change in lipid composition (14) are consistent with the wealth of evidence that lipid composition has an important influence on the geometry of the structures formed (15–20).

This evidence therefore indicates that lipids have a considerable role in determining membrane behaviour and thus cell structure. Taken with the importance of structural integrity for cell viability, this raises questions about how membrane systems are managed through the cell cycle. One suggestion is that the composition of lipid membranes is remodelled to minimise the energetic costs of changing their shape through the cell cycle (21). This led us to the hypothesis that the biosynthesis of structural lipids is linked to the cell cycle.

To test this hypothesis, we elected to use a model organism that has a well-characterised cell cycle, shows the structural challenges through the eukaryotic cell cycle as clearly as possible and comprises typical eukaryotic lipids. We also wanted to be able to collect samples of cells at different stages of the cell cycle without introducing artefacts associated with the drugs needed to synchronise cell cultures. All of these conditions were met by *Desmodesmus quadricauda*. This organism undergoes multiple fission producing eight daughter cells in one cell cycle (*Fig. 1*). Multiple fission is common in green algae, with some species dividing into up to 32 daughter cells in one mitosis (22,23). Cell division in *D. quad.* is also timed carefully as this photosynthetic species preferentially undergoes cytokinesis when there is insufficient light for the light-dependent part of photosynthesis. Thus, cultures can be synchronised through light and dark periods without the need for drug-based inhibition of DNA synthesis (24,25). *D. quad.* is also a good model for other eukaryotic cell types as it comprises similar lipids to most eukaryotes (21,26,27).

**Fig. 1.**
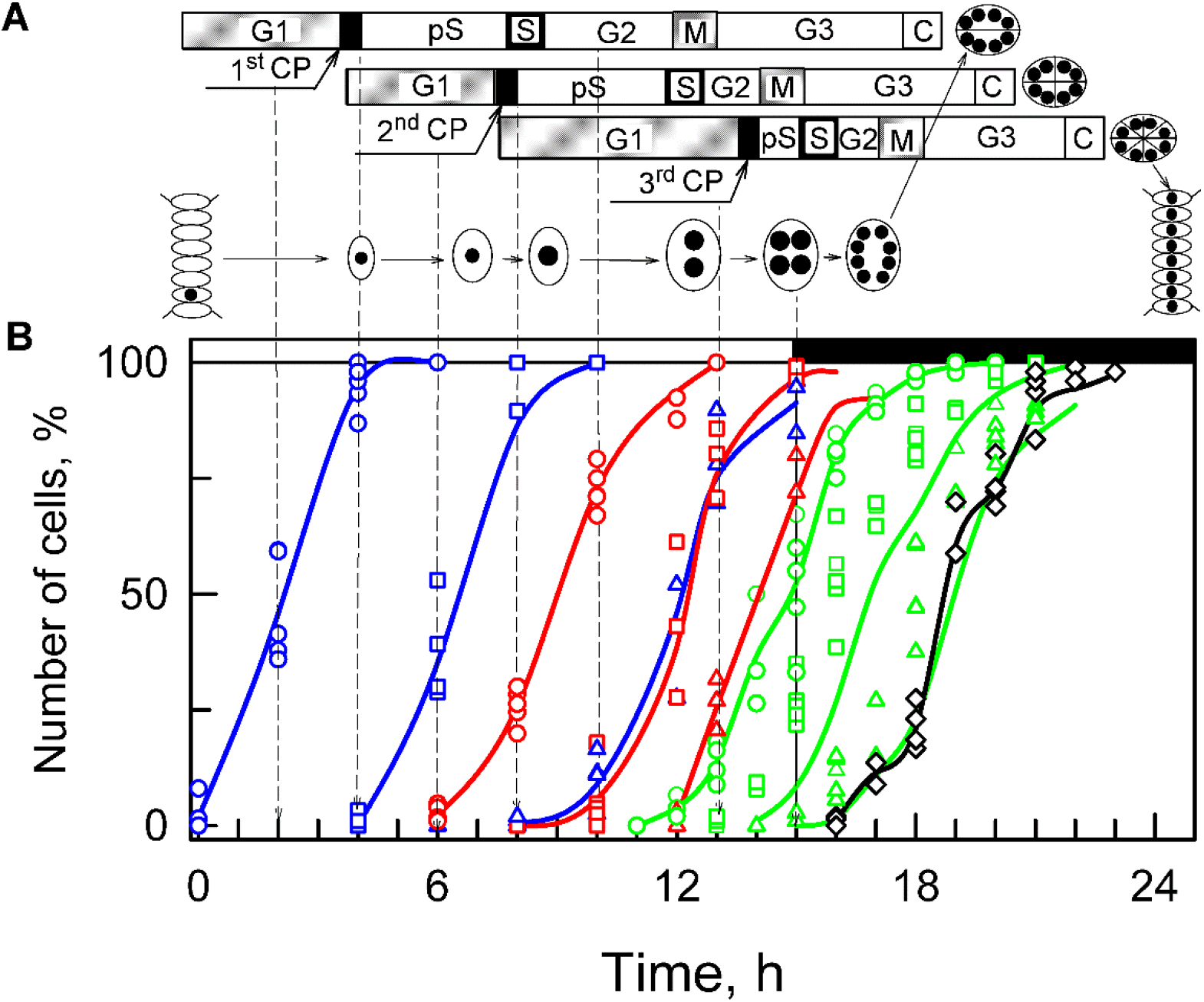
Diagram illustrating cell cycle pattern in synchronised chlorococcal alga Desmodesmus quadricauda grown under present experimental conditions (A). The three horizontal strips illustrate the simultaneous course of different phases from three consecutive sequences of growth and reproductive events. The individual sequences during which growth and reproductive processes lead to duplication of cell structures occur within one cell cycle. The cells divide into eight daughter cells connected in one coenobium. Designation of phases and events: **G_1_**, a **pre-commitment phase**, during which the threshold size of the cell is attained and completed by attainment of the commitment point. **CP**: **commitment point**, the stage at which the cell becomes committed to triggering and terminating of the sequence of processes leading to the duplication of reproductive structures (**post-commitment period**), which consists of: **pS**: the pre-replication phase between the commitment point and the beginning of DNA replication. The processes required for the initiation of DNA replication are assumed to happen during this phase. This phase is designated as late G_1_ phase in mammalian cells. **S**: the phase during which DNA replication takes place. **G_2_**: the phase between the termination of DNA replication and the start of mitosis. Processes leading to the initiation of mitosis are assumed to take place during this phase. **M**: the phase during which nuclear division occurs. **G_3_**: the phase between nuclear division and cell division. The processes leading to cellular division are assumed to take place during this phase. **C**: cytokinesis, the phase during which protoplast fission and forming of daughter cells is performed. Schematic pictures of cells indicate their size changes during the cell cycle and the black spots inside illustrate the size and number of nuclei. The size of the black spots indicates DNA level per nucleus. Time courses of individual commitment points, nuclear division, protoplast fission and daughter cell release (B) in the experimental cultures used in this work (n=6). Blue lines: cumulative percentage of the cells, which attained the commitment point for the first (circles), second (squares) and third (triangles) reproductive sequences, respectively; red lines: cumulative percentage of the cells, in which the first (circles), second (squares), and third (triangles) nuclear divisions were terminated; green lines: cumulative percentage of cells, in which the first (circles), second (squares) and third (triangles) cell divisions were terminated, respectively; black line, empty diamonds: percentage of the cells that released daughter coenobia. Light (15 hours) and dark periods (9 hours) are marked by stripes above panels and separated by vertical lines. The lines represent the means of at least six independent experiments. The raw values are plotted as dots and the line connects the mean values of the experiments. All values were calculated per parental cell, even after their division (17:00 to 22:00 h). Vertical dashed lines with arrows indicate time of sampling for lipidomics.

We cultivated populations of *D. quadricauda*, synchronised them and extracted the lipid fraction at defined points in the cell cycle. We developed a novel method for preparing cells that is compatible with established procedures for extracting lipids from biological samples (14,28,29). Lipid class abundance was measured using ^31^P Nuclear Magnetic Resonance Spectroscopy (NMR) and the fatty acid composition of lipids determined using high resolution mass spectrometry (HRMS). We assembled a *de novo* transcriptome of *D. quad.* to identify homologues of genes involved in cell cycle regulation and lipid metabolism. Finally, we used a combination of the reverse-transcription quantitative polymerase chain reaction (RT-qPCR) and Western Blotting to determine the mRNA and protein abundance of putative enzymes involved in lipid biosynthesis.

It was important to test this hypothesis because the physical integrity of cells through the cell cycle is a fundamental part of cell viability, but the control of the physical process is relatively poorly understood. Understanding how the structure of cells fail is of interest in controlling cell growth, either to hinder it (antibiotics, anti-tumour compounds) or to promote it (tissue regeneration). The processes that exist in evolved systems also have implications for preparations of artificial cells and for nanotechnology.

## Results

In order to test the hypothesis that lipid biosynthesis is linked to the eukaryotic cell cycle, we profiled the abundance of structural lipid classes through the cell cycle alongside a formal characterisation of growth and the cell cycle (*Fig. 2*). The growth characterisation was done through the cell volume and mass of RNA per cell, while that of cell cycle was done through monitoring DNA replication (DNA mass) and nuclear divisions (number of nuclei) per cell. As expected under given growth conditions, the cells increased their volume about eight-fold, which was accompanied by increase in mass of major macromolecules such as RNA (*Fig. 2*). Concomitant with cell growth the cells entered three sequences of DNA replication sequentially, nuclear and cellular division leading to division into eight-celled daughter coenobia (*Fig. 2*). We developed novel procedures for handling cultures as established procedures for handling this cell type for proteomics were not suitable for handling the lipid fraction. The lipids were extracted (*n* = 6 biological replicates) in a similar manner to previous work (14,28–30). Phosphorus (^31^P) NMR was then used to measure the relative abundance of lipid classes (28,29).

**Fig. 2.**
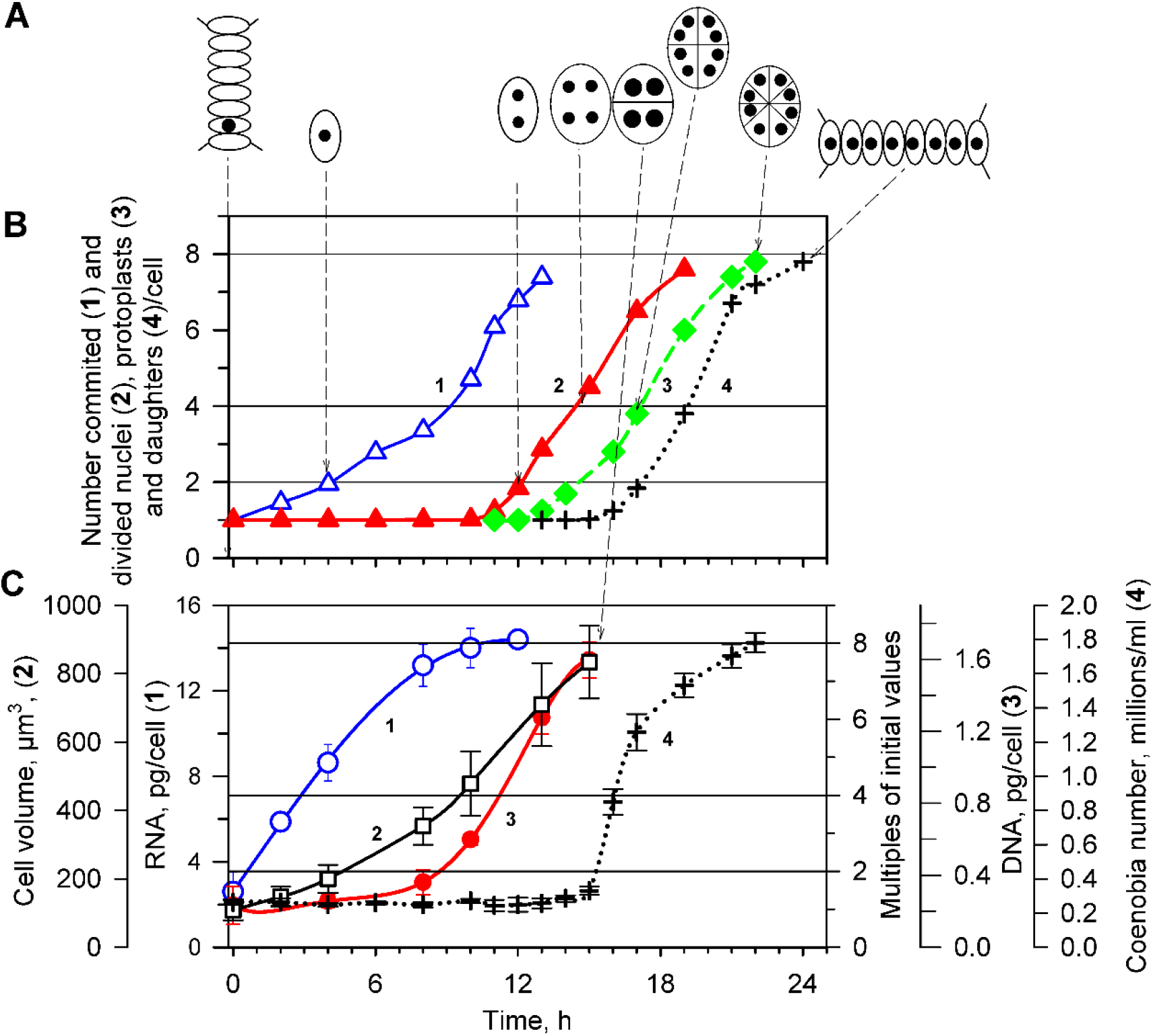
Characteristics of cell cycle progression and growth in the cultures used in this study: (A) Schematics of cell in the population at particular time. (B) Time course of **cell cycle events**: attainment of commitment points (curve 1, blue triangles △), nuclear divisions (curve 2,red triangles ▲) protoplast fissions (curve 3, green diamonds ♦) and daughter coenobia release (curve 4, black crosses **+**). The values plotted here are a cumulative version of the same data plotted on Fig. 1 to allow direct comparison with growth events. (C) Time course of **growth events**: RNA (curve 1, blue circles ○), mean cell volume (curve 2, black squares □) DNA replications (curve 3, red circles ⚫), and changes in cell number (curve 4, black crosses). Horizontal solid lines indicate doublings of initial value, vertical dashed lines with arrows indicate the schematics of a cell at given timepoint. All data of analytical analyses are presented as means ±SD of six experiments. Growth conditions: incident light intensity 750 μmol m^−2^ s^−1^, mean light intensity 530 μmol m^−2^ s^−1^, continuous illumination, 2 % of CO_2_ in aerating air, temperature 30 °C.

This showed that the relative abundance of major phospholipids was modulated through the cell cycle (*Fig. 3*). The lipid fraction was dominated by four lipid classes through the cell cycle: phosphatidylcholine (PC), phosphatidylethanolamine (PE), phosphatidylinositol (PI) and phosphatidylglycerol (PG). They make up over 95% of the phospholipid fraction between them (*Fig. 4*). During the G_1_ phase, in which the greater part of cell expansion occurs, the relative abundance of PC, PE and PG changed significantly (*Fig. 4*), including a significant change after the first commitment point. We therefore analysed the relationship between lipid biosynthesis through particular phases, as well as throughout the whole cell cycle. These are described in order.

**Fig. 3.**
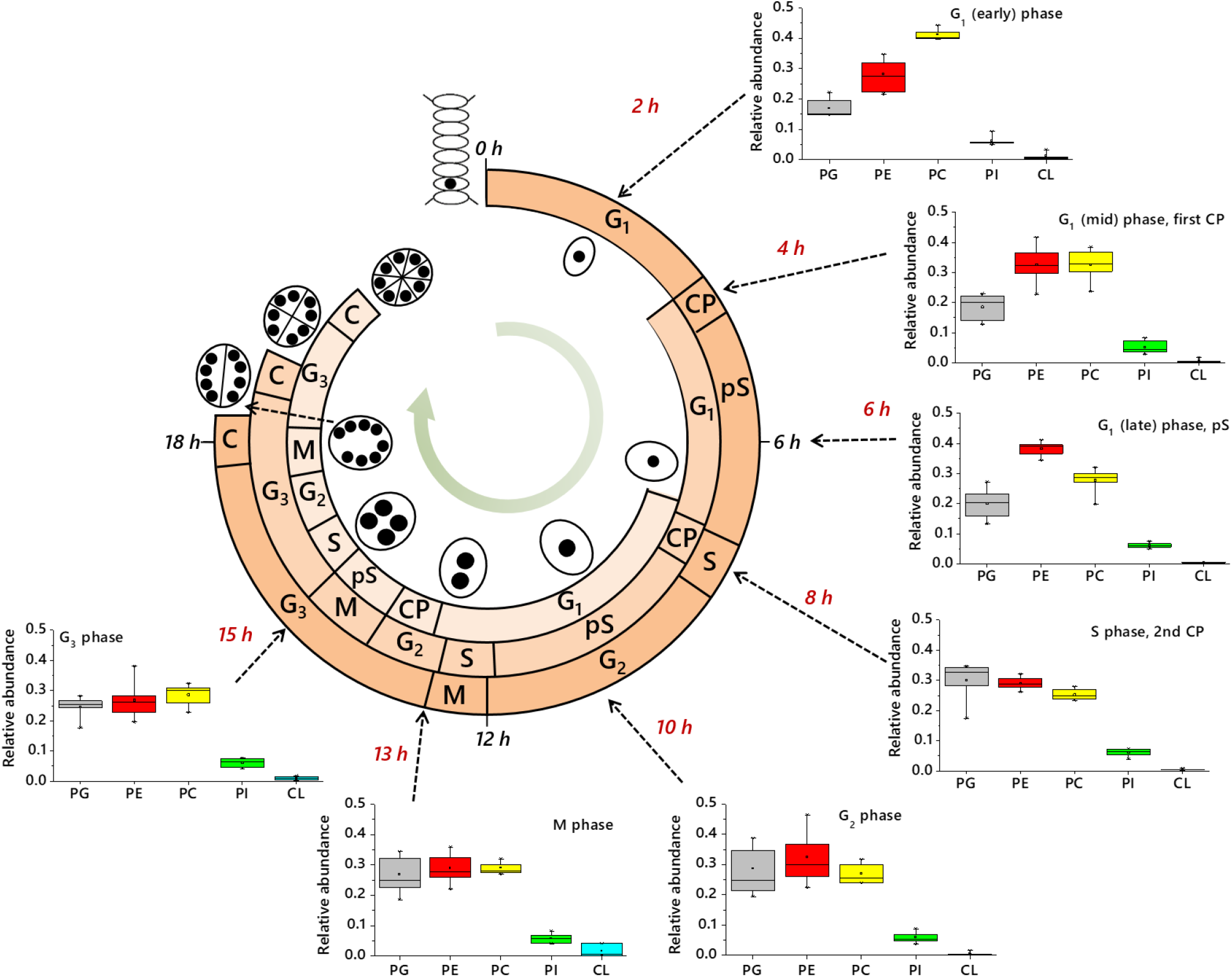
The relationship between the multiple fission cell cycle of D. Quadricauda and the abundance of its principal structural lipids. Inset box and whisker plots show the distribution of n *=* 6 values collected from ^31^P NMR measurements for the five most abundant phospholipids. Ordinate axes show the relative abundance as a fraction of total lipid as 1·0. Abscissa axes show lipid head groups. Dashed arrows indicate sampling times (times shown in red). Schematic representations of the DNA synthesis/nuclei in the cell shown towards the middle. Lipid abbreviations: PC, phosphatidylcholine; PE, phosphatidylethanolamine; PG, phosphatidylglycerol; PI, phosphatidylinositol. Phases of the cell cycle: CP, commitment point; G_1_, first gap phase; G_2_, second gap phase; G_3_, third gap phase; M, Mitosis; pS, pre-synthesis phase.

**Fig. 4.**
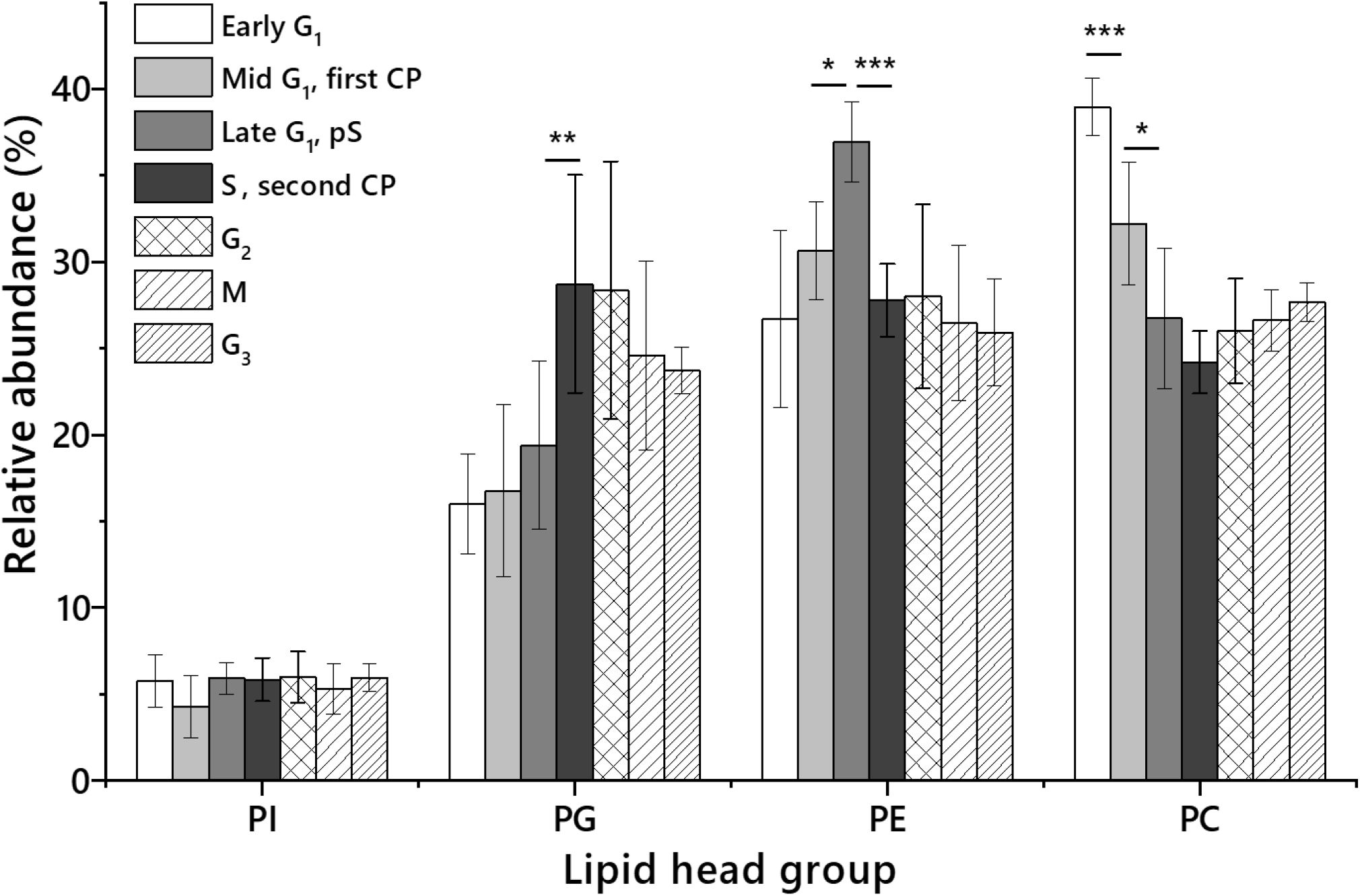
The lipid head group profile of the four most abundant lipids in D. quadricauda, determined using ^31^P NMR. The integral of resonance(s) assigned to each head group was calculated as a fraction of the total for that spectrum and the mean and standard deviation of all values taken to generate the values used. n = 6 values collected. Collection points of cell cultures are assigned as: +2 h, early part of G_1_; +4 h, mid-G_1_ (first CP); +6 h, end of G_1_ and pS; +8 h, S and second CP; +10 h, G_2_; +13 h, M; +15 h, G_3_. The presence of all lipid species was verified by HRMS/MS. Error bars indicate standard deviation. Asterisks indicate p-*values from Student’s* t*-tests that fall below given thresholds; *,* p*<0·05; **,* p*<0·01; ***,* p*<0·001.* PC, phosphatidylcholine; PE, phosphatidylethanolamine; PG, phosphatidylglycerol; PI, phosphatidylinositol.

*Lipid remodelling in G_1_*—The relative abundance of PC falls by a third in the G_1_ phase (*Fig. 4, Table S1*). However, during this phase, the cell volume increases by a factor of ~2·5 (*Fig. 2*), indicating that the overall mass of phospholipid in a cell increased by around the same factor. This suggests that the biosynthesis of PC does not occur at the same rate as cell growth. We therefore investigated the control of the biosynthesis of PC. There are two well-characterised routes for the biosynthesis of PC in eukaryotes. One is the methylation of PE, the other is the transfer of choline phosphate onto a diglyceride. We assembled a *de novo* transcriptome of *D. quadricauda* and then we investigated with combination of RT-qPCR, Western Blotting and lipidomics techniques to investigate which pathway was more important.

In the assembled transcriptome, we searched for putative homologues of individual enzymes involved in methylation of PE. However, we were unable to find phosphatidyl-*N*-methylethanolamine *N*-methyltransferase (PEMT). Mass spectrometric profiling of lipids shows that there are a number of isoforms of PC that are without equivalent in PE (*Fig. S3*), suggesting that not all PC can come directly from PE. Furthermore, a large increase in the abundance of PE is not met with an increase of PC, suggesting that PE is not a substrate for biosynthesis of PC. Taken together, this suggests that PE is not generally used to make PC in this organism and that the two lipids are not directly linked metabolically. This strongly suggested that PC was produced endogenously through transfer of choline rather than methylation of PE.

The increase in the relative abundance of PE during the G_1_ phase (40%) is accompanied by a factor of eight increase in the abundance of mRNA and factor of 2 increase in the abundance of the protein Ethanolamine Phosphate cytidylyl Transferase 1 (EPT1), the enzyme that synthesises PE from ethanolamine phosphate and diglycerides (*Fig. 5, S4*). This showed that the synthesis of PE is based upon transfer of ethanolamine onto a diglyceride rather than modification of PC or decarboxylation of phosphatidylserine (PS). This is therefore consistent with the results of the biosynthesis of PC that show that PC and PE are produced separately and not interconverted.

**Fig. 5.**
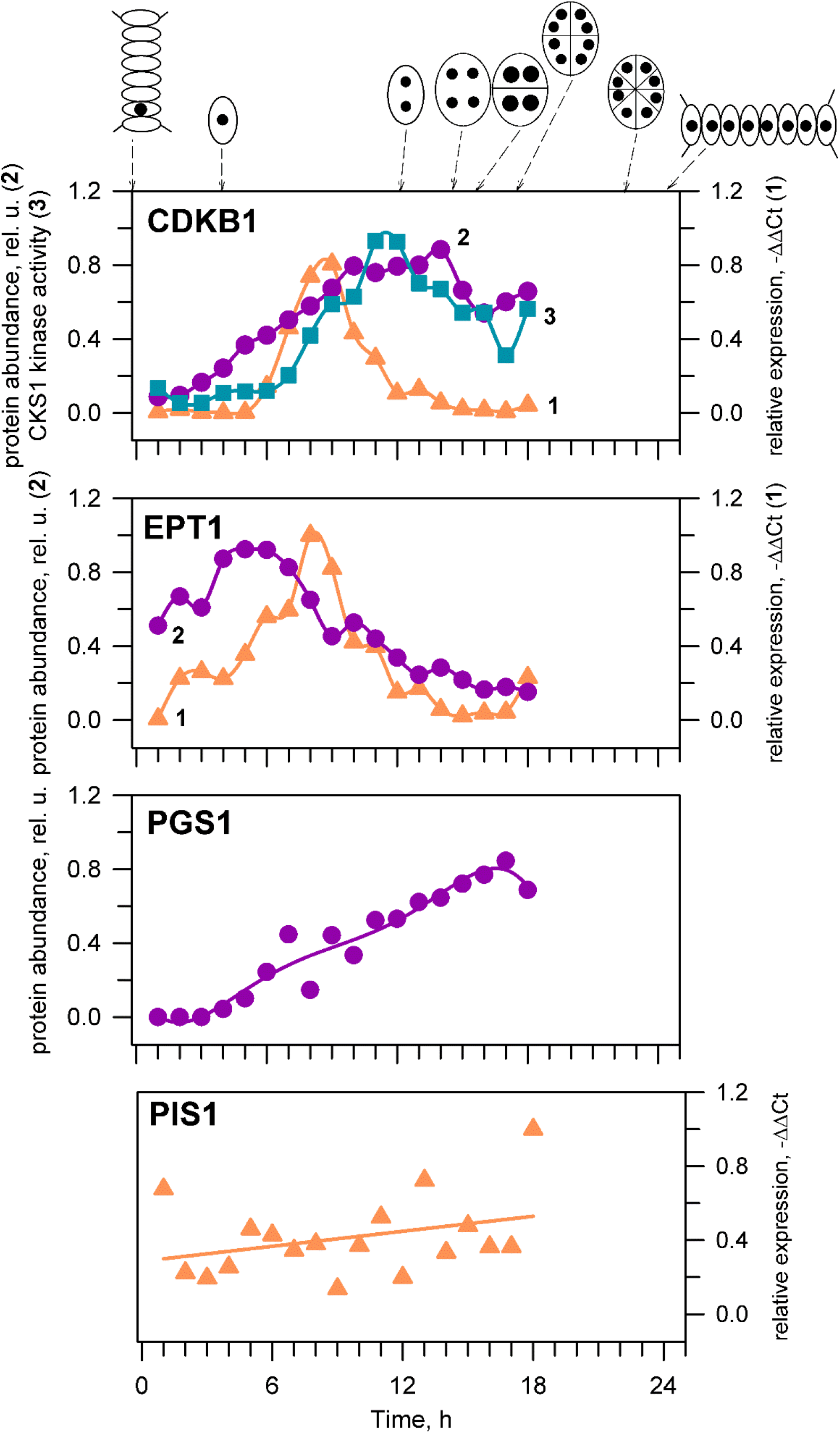
The mRNA expression (RT-qPCR), protein abundance (Western blotting) and activity (kinase activity) of a single cell cycle (CDKB) and three lipid metabolism genes during cell cycle. The cell cycle progression is documented by schematics above the panels and by the mRNA abundance (orange tringles), protein abundance (purple circles) and activity (light blue squares) of a cell cycle regulator, CDKB. Abundance of mRNA (orange triangles) and protein abundance (purple circles) of ethanolamine phosphate transferase (EPT1), phosphatidylglycerol synthase (PGS1) and phosphatidylinositol synthase (PIS1). The data from RT-qPCR were normalized against 18S RNA. The data from Western blotting were normalized to the signal of RuBISCo in the same samples. All the data were normalized to the maximum value in the dataset to allow for a simple comparison.

**Fig. 6.**
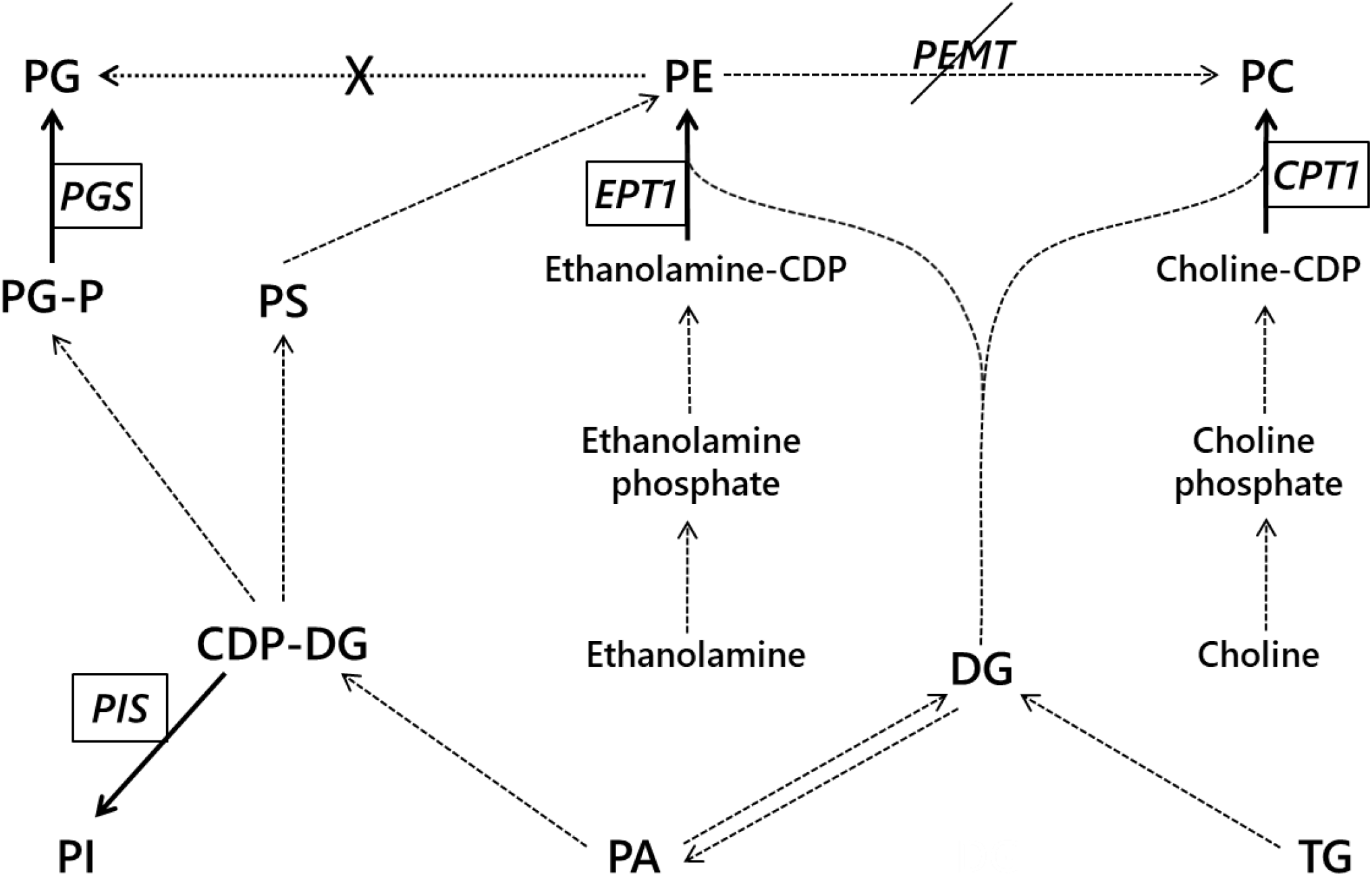
General phospholipid metabolism in the cytosol and Endoplasmic Reticulum of Chlorella spp. and other microalgae (71–74). Dashed arrows represent previously reported conversions, enzyme names omitted for clarity. Solid lines represent positive identification of the appropriate enzyme and resulting lipid in the present study. A dotted line with a cross indicates a connection that could not be made in the present study. Phosphatidylethanolamine N-methyltransferase (PEMT) could not be found in this organism. CPT1, choline phosphate transferase; EPT1, Ethanolamine phosphate transferase; PGS, phosphatidylglycerol synthase; PIS, phosphatidylinositol synthase. CDP-DAG, cytidine diphosphate diglyceride; CL, cardiolipin; DG, diglyceride; PA, phosphatidic acid; PC, phosphatidylcholine; PE, phosphatidylethanolamine; PG, phosphatidylglycerol; PI, phosphatidylinositol; PS, phosphatidylserine; TG, triglyceride.

### Lipid remodelling over the pS/S boundary

There is a rapid increase in the relative abundance of PG and a reduction in that of PE towards the end of the pS phase and into the replication phase (S). This suggested that PE is biosynthesised rapidly in G_1_ before the relative abundance of PG overtakes it. We therefore tested the hypothesis that biosynthesis of PG relied upon PE.

A comparison of the fatty acid profile of PE and PG suggests that they are not closely related (*Fig. S3*). The fatty acid residue (FAR) profile of PG is more consistent with prokaryotic lipid metabolism (more saturated, shorter FARs), where PE is more eukaryotic (dominated by FA(16:4) and unsaturated C_18_ FARs), *Fig. S4*. This is consistent with our understanding that the bulk of the PG fraction resides in chloroplasts (a prokaryotic compartment) and the bulk of PE is typically in the plasma membrane and eukaryotic compartments. Lastly, enzymes involved in converting PE to PG have not yet been reported. This led us to determine how the synthesis of PG is controlled with respect to the cell cycle.

Western blotting revealed that the abundance of PG synthase 1 (PGS1) increases by a factor of 22 from the beginning of the pS phase, and starts to fall back zero by the start of the cell division, preceded by chloroplast division (*Fig. 5 and S4*), indicating that PG is produced by this enzyme from cytidyl diphosphate-diglyceride (CDP-DAG) and glycerol-3-phosphate. These data are consistent with *de novo* synthesis of PG from phosphatidic acid in chloroplasts and not from PE, indicating a control of the biosynthesis at pS/S boundary that is not tied to other lipids.

### Lipid remodelling in the G_2_, M and G_3_

The abundance of scarcer lipids was modulated through Mitosis and G_3_ (*Table S1*). Small amounts of phosphatidic acid (PA, FAR profile in *Table S2*), PS (*Table S3*) and evidence of plasmalogens of PC and PE were found. The abundance PA is 0·5-1·0% from G_1_ to the end of G_2_, but increases to around 3% during mitosis. PC-plasmalogen may also increase in abundance during the same period. As the cell increases in volume only from ~710 μm^3^ to ~830 μm^3^ (*Fig. 2*) during G_3_ and Mitosis, these shifts are not expected to be consistent with major structural changes in either eukaryotic or prokaryotic compartments. PE-plasmalogen and PS remain roughly constant at ~1% throughout the cell cycle (*Table S1*). Although such species have been detected before (31,32), it is not clear what their role is in the cell cycle, if any.

### Lipid remodelling across the whole cell cycle

The relative abundance of PI is maintained at ~5% throughout the cell cycle *(Fig. 4*), indicating that rate of its synthesis correlates closely with the increase in the volume of the cell. However, the population of PI molecules must increase by a factor of eight through the cell cycle. RT-qPCR indicated that the expression of PI synthase (PIS) is constant through the cell cycle (*Fig. 5*). This implied a constant supply of PI relative to other phospholipids and suggests its production is separate from that of PG, PE and PC. Fatty acid profiling of isoforms of PI and PC showed that PI’s profile is not consistent with PC (*Fig. S1*), nor with the much less abundant PIPs (*Fig. S2*).

This suggested first that PI and PC are not generally interconverted and second that there are many isoforms of PI that have non-signalling roles, something also observed in HeLa cells (33). PI appears only to be present in the membranes of eukaryotic compartments (34), suggesting to be exclusively eukaryotic in origin. The FA profile of PI in *D. quadricauda* (*Table S1*) is characterised by polyunsaturated and longer chain configurations. These results are consistent with PI as a lipid that is required in small amounts throughout the cell cycle. Through HRMS alone, we also observed the presence of a number of known phytolipids and phytosterols throughout the cell cycle that have not been observed in this species before (*Tables S4–6*). These species were not visible by ^31^P NMR so were not quantified and thus could not be linked to the cell cycle.

## Discussion

This study was motivated by the hypothesis that the biosynthesis of structural lipids is linked to the cell cycle in eukaryotes. Testing this hypothesis provided evidence that the biosynthesis of the three most abundant phospholipids (PC, PE and PG) in the model used are modulated through the cell cycle and linked to it. PC, PE and PG all dominate the overall lipid fraction at different points. This is in contrast to PI that does not change in relative abundance through the cell cycle. Evidence from MS, Western blots and RT-qPCT suggests that PC, PE and PG, and the next most abundant, PI, are not interconverted between one another and are thus metabolically independent after the CDP-DAG is assembled.

The evidence for independent synthesis of the four most abundant lipids and that three of them dominate the organism’s lipid fraction at different stages of the cell cycle is consistent with cell-cycle-based control of lipid biosynthesis through different pathways that are switched on and off appropriately, and locally. It also characterises those stages of the cell cycle as having a particular focus in their lipid metabolism. Furthermore, the change in abundance of EPT1 and PSG1, and the increase in abundance of PE and PG that follow, suggests that these are the principal synthetases of these lipids in *D. quad*. The abundance of the enzymes increased rapidly just before the abundance of the appropriate lipid does, and then most of the enzyme is lost in a way that limited synthesis of those lipids, up to 90% in the case of EPT1. This indicated that the final step in the biosynthesis of those two crucial lipids occurs through only one enzyme each, and that the expression of both enzymes is linked to the cell cycle.

Significant changes in PC, PE and PG—lipids that are both abundant and well known to have structural roles—invites questions about the physical role of these molecular species. Studies of lyotropic phase behaviour have established that PG and PC are bilayer-forming lipids and that the anionic PG may have an important charge-based effect (35). Unlike PC and PG however, PE is a non-bilayer forming lipid that typically favours the assembly of curved lipid mesophases (15). Recent work in biological systems indicates that PE is part of the control of fluidity of membranes *in vivo* (36) and has an essential role in cytokinesis in at least one eukaryotic cell type (37). This suggests that an increase in the abundance of PE in the G_1_ phase represents a shift in membrane properties through this phase. PE’s propensity for forming inverse mesophases (15,38) suggests that more curved membranes are required as the cells enter from the pS to S phase. During this period, preparations for DNA synthesis are made but the cells do not grow as rapidly (22), suggesting re-organisation is more important than expansion at this point. This is consistent with the theory that cells remodel their lipid composition in order to lower the energy of the succeeding phase (21), as the internal structural needs of the cell change at this point. The precise role and distribution of PE is required to understand this more deeply and could be used to answer the importance of the site of biosynthesis in driving membrane reorganisation.

These results are also of interest in the light of evidence that the rate as well as the timing of lipid biosynthesis differs between lipid classes. For example, it is not clear from this work that the biosynthesis of PI has a peak, either through the abundance of the lipid or the expression of its synthase (PIS). It is possible that it is produced continuously, correlating with cell size. However, it is not clear from our data whether there is a mechanism that links PI synthesis to cell size or whether inhibiting PI synthesis would disrupt progress through the cell cycle through structural means. This is important because PI is also beginning to be recognised as a non-bilayer lipid. Studies of the lyotropic phase behaviour of PI have shown that it has a concentration- and time-dependent effect on the geometry of lipid systems, induces considerable inverse curvature at lower hydrations (17) and introduces defects into lipid bilayers (39). This is similar to PE that is also characterised by inverse curvature (15). This evidence hints that a local abundance of PI may be able to reduce the energetic cost of membrane fission. Experiments of the distribution of lipids in Chinese Hamster Ovary cells have shown that PE and PI-derived PIP2 must be present the cleavage furrow of CHO cells in order for cytokinesis to take place (37,40,41). This suggests that lipids associated with inverse curvature such as PI and PE have a role in membrane scission.

The increase in abundance of PG and evidence that it arises from prokaryotic rather than eukaryotic lipid biosynthesis is consistent with an expansion in chloroplast size at that point in the cell cycle. This is consistent with a link between lipid biosynthesis and chloroplast as the structure making up the vast majority of cell volume. The role of PG in photosynthesis is well-established (42). PG’s lyotropic behaviour under physiological conditions is dominated by bilayer (membrane-like) systems (19). It is also anionic, suggesting that this lipid’s synthesis is associated with an increase in membrane area and a change in electrostatic interactions. Despite being a bulk lipid, evidence that PG has a role in regulating protein orientation in chloroplasts is emerging (43). PGS1 has also been found in chloroplasts in the alga *Chlamydomonas reinhardtii* (44) and in higher plants (45,46). This suggests that PG has a role in chloroplast membranes in all plants, adding to the view of this lipid as a ubiquitous one (47).

Recent work on fission yeast has begun to show that a gene involved managing lipid metabolism is involved the controlling the cell cycle (8). The gene m*ga2* regulates lipid homeostasis (48) and lipid synthesis (49) and when deleted, leads to cells that are unable to correct size deviations within individual cell cycles (8). This is consistent with the ‘sizer’ hypothesis about the control of cell division in eukaryotes (9,10) and may also apply to *D. quad.* as there is a clear size component to the preparation of a cell for multiple fission. This study shows that individual enzymes involved in lipid metabolism are linked to the cell cycle, however this work suggests that more general genes governing cell size are also involved. This is interesting because it can be used to inform the interpretation of cell-cycle-based lipid biosynthesis in other studies.

This includes general shifts in lipid abundance during cell elongation (50) but also specific ones such as the increase in the abundance of PA in the leaves of *Arabidopsis thaliana* during dark periods (51). The latter may be consistent with our observations of an increase in the abundance of PA during mitosis. Jueppener *et al.* reported a qualitative study of lipids through the cell cycle of *Chlamydomonas reinhardtii*, finding evidence for shifts in the lipid profile through that organism’s cell cycle and highlighting the cyclical nature of lipid metabolism in that species (52). Interestingly, *Chlamydomonas reinhardtii* appears to make considerable use of uncharged glyceride lipids such as *mono-*galactosyl diglycerides (MGDGs) and *di*-galactosyl diglycerides (DGDG) (53). In the present study, we found galactosyl-glycerides and others in *D. quad.* that have not been reported in this species before (*Tables S4–6*).

The evidence for the biosynthesis of lipids at particular points of the cell cycle raises questions about the extent of the link between the two. One question is whether there is a particular link between the availability of fatty acids and the synthesis of phospholipids. Evidence that particular FAs can favour the synthesis of phospholipids is beginning to emerge (54), suggesting a link between *de novo* fatty acid synthesis and progression through the cell cycle. Indeed, studies in yeast have provided evidence for a lipase that releases FAs from TGs that is linked to progress through the cell cycle (55).

An understanding of polyunsaturated fatty acid synthesis in algae may also be useful for harnessing them for industrial triglyceride synthesis (56,57). This is attractive as a sustainable source of essential fatty acids such as docosahexaenoic acid (DHA) and eicosapentaenoic acid (EPA). Evidence for lipid metabolism being linked to progress through the cell cycle implies that fatty acid biosynthesis is not directed solely towards triglyceride metabolism. Thus, a characterisation of lipid metabolism through the cell cycle can be used to inform the preparation of industrial cultures in which the balance between progress through the cell cycle and accumulation of triglycerides is struck.

## Conclusions

This study was motivated by the hypothesis that the biosynthesis of structural lipids is linked to the cell cycle. Lipid profiling showed that the lipid fraction is remodelled several times through the cell cycle in this organism. A combination of mass spectrometry, proteomics and transcriptomics indicate that the most abundant phospholipids are not directly connected to one another metabolically, but are connected to progress through the cell cycle. This has implications not only for our general understanding of the cell cycle, but also our understanding of the physical aspects of cell division. It may also be useful for informing the design of artificial cells and the use of algae for industrial production of triglycerides and characterisation of the migration of biomass in food chains through nutrients such as DHA. Continuous re-modelling of the lipid fraction through the course of the cell cycle implies that lipid metabolism is as important as that of proteins and nucleic acids for the success of this process.

## Experimental Procedures

### Reagents & Chemicals

Solvents, and fine chemicals were purchased from *SigmaAldrich* (Gillingham, Dorset, UK) except phosSTOP tablets were purchased from Roche (Welwyn, Hertfordshire, UK; stored at 4°C). Chemicals for the growth medium were purchased from Penta (Chrudim, CZ).

### Cultivation of D. quadricauda

The experimental organism *Desmodesmus quadricauda* (Turpin) Brébisson (previously known as *Scenedesmus quadricauda*), strain Greifswald/15, was obtained from the Culture Collection of Autotrophic Organisms, Institute of Botany (CCALA, Czech Acad. Sci., Třeboň, CZR). The synchronous cultures of *D. quadricauda* were cultivated in flat glass photobioreactors (3 L) in liquid mineral medium (58) at 30 °C and continuous light. The photobioreactors were illuminated from one side by fluorescent lamps (Osram DULUX L, 55W/840, Italy) at a surface incident irradiance of 750 μmoL m^−2^ s^−1^. Cultures were aerated with air containing 2% carbon dioxide (*v/v*).

Synchronisation was done according to reported procedures (24). Briefly, before the start of the experiment, the cells were synchronized and then grown for one more whole cell cycle. At the beginning of the following light period they were diluted to the initial density (1×10^6^ cells mL^−1^). The synchronization itself was carried out by alternating light/dark periods (15 h/10 h), the lengths of which were chosen according to the growth parameters of the cells. The optimum time for turning off the illumination was when the cells started their first protoplast fission. The length of the dark period was chosen to allow all cells of the population to release their daughter cells. Under the conditions described above, cell division started at about the 15th hour of the cell cycle and the cells typically divided into eight daughter cells (*Fig. 1, 2*).

### Preparation of dried cell lysates

The active culture (650 mL) was filtered and the filtrate collected and centrifuged (4000 × *g*, 5 min). The resulting pellet was resuspended in a mixture of chaeotropes (5 mL; thiourea 1·5 M and guanidinium chloride 6 M), phosSTOP (1 tab/sample, dissolved in 1 mL PBS) and 2-butoxyphenylboronic acid (BPBA, 2 mg/mL final concentration, ethanolic stock solution 100 mg/mL). The suspension (~10 mL) was agitated vigorously with glass beads (1 min, air-tight Falcon tube, 50 mL) before being frozen in liquid nitrogen and then freeze-dried. The freeze-dried material was stored under a nitrogen atmosphere and transported at room temperature.

### Isolation of lipid fraction

Freeze-dried cell lysate was powdered (pestle) and rehydrated (PBS, 5 mL) with agitation (1 min) but without sonication. The mixture was frozen (193K) and freeze-dried. The resulting dry, free-flowing powder was resuspended in a mixture of dichloromethane (20 mL) and water (20 mL) and diluted with sufficient methanol to make a stable uniphasic solution (40-45 mL, 500 mL separating funnel). The mixture was then made biphasic by addition of dichloromethane (20 mL). The dichloromethane solution was separated and the aqueous solution washed (dichloromethane, 20 mL). Triethylammonium chloride (TEAC) was added to the remaining aqueous solution (final concentration of 2 mM, 2M stock) and the aqueous solution washed with dichloromethane (2 × 20 mL). The combined organic solutions (~90 mL) were filtered through filter paper and concentrated *in vacuo* before storage of the resulting lipid film under nitrogen at −20°C.

### Solution phase ^31^P NMR

Lipid films were dissolved in the CUBO solvent system (28,29) (450 μL, 23-26 mg isolate/sample). Data acquisition was similar to published work (13,14), but using a Bruker 400 MHz Avance III HD spectrometer equipped with a 5 mm BBO S1 (smart) probe operating at 298K. ^31^P NMR spectra were acquired at 161·98 MHz using inverse gated proton decoupling, with 2048 scans per sample and a spectral width of 19·99 ppm. An overall recovery delay of 6·5 s was used which gave full relaxation. Data were processed using line broadening of 2·00 Hz prior to zero filling to 19428 points, Fourier transform and automatic baseline correction. Spectra were processed and analysed using TopSpin 3.2. The dcon function was used to deconvolute spectra in order to determine the integration of each resonance, in a similar manner to previous studies (28,29). The integration of each resonance was divided by the total integration for that spectrum, and assigned according to known shifts (14). The integrations (as fractions of the total for that spectrum) were used for statistical calculations (*n* = 6 spectra).

### Mass Spectrometry of Lipids

Samples were prepared and measurements taken in a similar manner to published methods (28,59–61). Raw data were processed using software by Kochen *et al.* (62) with some additional code (14). Dried lipid fractions (~5 mg) were dissolved (isopropanol/dichloromethane 1:1, 300 μL) for accurate mass LC-MS (ThermoFisher Q exactive with Dionex Ultimate 3000 sample handler and Waters acquity UPLC BEH C18 with 1·7 μm particle size LC column) in ES+ mode. Gradient separation was performed at 40°C, analysis time 20 min, flow rate 0·4 mL/min. Mobile phase A consisted of 0·1% formic acid in water at *p*H 6·0, and mobile phase B was 55% acetonitrile, 40% isopropanol and 5% water with 0·1% formic acid. Ions were monitored the range *m/z* 300 to 2000. The calibrated mass accuracy was 1 (ES+) milli mass units and the resolution was 140,000 for MS1 and 17,500 for MS2 spectra. Analyses were performed using Thermo Xcalibur 3.0.63. Original code for non-standard head groups was written in Matlab R2015b.

The lipid signals obtained were relative to the total, *i.e.* ‘semi-quantitative’ or relative abundance, with the signal intensity of each lipid expressed relative to the total lipid signal intensity, for each individual, per mille (‰). Raw high-resolution mass-spectrometry data were processed using XCMS (http://www.bioconductor.org) and Peakpicker v 2.0 (an in-house R script). Lists of known species (by *m/z*) were used, *n =* 1740 incl. standards (28,60,61,63). Signals that deviated by more than 5 ppm were ignored, and thus assignments were made on the basis of HRMS only.

### Protein extraction

Whole cell protein extracts were prepared as described previously (6,64). Briefly, samples consisting of 2×10^7^ cells were harvested and centrifuged, and the pellets washed (SCE buffer [100 mM sodium citrate, 2.7 mM EDTA-Na_2_, pH 7 (citric acid)], 1 mL). The pellets were frozen in liquid nitrogen (193K) and stored at −70°C. Extracts were subject to both Western immunoblotting and a kinase assay.

### Western immunoblotting

Protein extracts were mixed with 5×SDS-PAGE sample buffer (250 mM Tris-HCl (pH 6.8), 50% (*w/v*) glycerol, 10% SDS, 100 mM dithiothreitol, 0.5% (*w/v*) bromophenol blue), incubated 5 min at 65 °C and separated by 12% SDS-PAGE (65) in the Mini Protean 3 Apparatus (BioRad Laboratories, Hercules, CA, USA). The final concentration of proteins was 10 μg per lane. After the separation, proteins were transferred onto a PVDF membrane (pore size 0.45 mm, Immobilon-P, Millipore, www.millipore.com) (66) at 1 mA/cm^2^ for 1·5 h. The membrane was blocked in 5% (*w/v*) skimmed dry milk solution in TBS-T buffer (20 mM Tris *p*H 7.5, 0.5 M NaCl, 0.05% (*v/v*) Tween 20), over night at 4°C. The proteins of interest were probed with following primary antibodies: anti-CDKB1 rabbit antiserum (diluted 1:1000) raised against a QDLHRIFPSLDDSGC peptide of *C. reinhardtii* CDKB1 protein (Genscript, www.genescript.com)(64), anti-CYCB rabbit antiserum (diluted 1:1000) raised against CKYSSTKYNEAAKKP peptide of *C. reinhardtii* cycB protein (67), anti-EPT1 rabbit antiserum (diluted 1:2000) raised against a LPAKERAHKQLGQCG peptide of *D. quadricauda* putative EPT protein, anti-PGS1 rabbit antiserum (diluted 1:2000) raised against a LGVNREEEDFVSPYC peptide of *D. quadricauda* predicted PGS protein (Genscript, www.genescript.com), and anti-Rubisco goat antiserum (diluted 1:3000) (Santa Cruz Biotechnology, Santa Cruz, CA, USA). Secondary antibodies were peroxidase-conjugated goat anti-rabbit IgG (A9169 Sigma, www.sigmaaldrich.com) (diluted 1:40 000), and peroxidase-conjugated rabbit anti-goat IgG (A5420 Sigma, www.sigmaaldrich.com) (diluted 1:40 000). Immunoreactive bands were detected by chemiluminescence (SuperSignal™ West Dura Extended Duration Substrate, Thermo Fisher Scientific, USA, www.thermofisher.com) according to the provided protocol and were visualized using a luminiscent image reader (ImageQuant LAS4000, GE Healthcare Bio-Sciences AB, Uppsala, Sweden). The extent of chemiluminiscence was quantified using Image Studio Lite software (LI-COR Biotechnology, www.licor.com). To compare between samples and experiments, the sum of pixel intensity within the same area was normalized to the background pixel intensity to yield pixel intensity values of the signal. Each experiment was repeated at least three times and representative experimental results are shown.

### Kinase assay

The same number of cells from the same volume of culture was used; the cultures were not diluted during experiments. The cleared protein lysates (see above) were used immediately for the assay or were affinity purified by CrCKS1 beads as described (6) and incubated at 4°C for 2 h (64). Histone H1 kinase activity was assayed as previously (68) in a final volume of 10 μL with either 7 μL of clear whole cell lysate or CrCKS1 beads fraction corresponding to 20 μL of whole cell lysate. The reactions were started by adding the master mix to a final composition of 20 mM HEPES, *p*H 7.5, 15 mM MgCl_2_, 5 mM EGTA, 1 mM DTT, 0.1 mM ATP, 0.2% (*w/v*) histone (Sigma H5505) and 0.370 MBq [γ ^32^P] ATP. All reactions were incubated for 30 minutes at room temperature and stopped by addition of 5 μl of 5 × SDS sample buffer [250 mM Tris-HCl, pH 6.8, 50% (*v/v*) glycerol, 10% (*w/v*) SDS, 100 mM dithiothreitol, 0.5% (*w/v*) bromphenol blue], incubated 2 minutes at 98°C and immediately cooled. Proteins were loaded on 15% gels and separated by SDS-PAGE (65). Phosphorylated histone bands were visualized by autoradiography and analysed using a phosphoimager (Storm 860, GE Healthcare Bio-Sciences AB, Uppsala, Sweden). The extent of phosphorylation was quantified as described above.

### Quantitative real-time polymerase chain reaction

Cell pellets containing 2 × 10^7^ cells were harvested during the cell cycle, re-suspended in a DNA/RNA Shield (cat. no. R1100, Zymoresearch, Irvine, CA, USA) and stored at −20 °C. RNA was isolated using a Quick RNA plant kit (cat. no. R2024, Zymoresearch, Irvine, CA, USA) according to the manufacturer’s instructions including DNase treatment. The quality of RNA was verified by gel electrophoresis and only intact RNA was used for cDNA synthesis. cDNA synthesis was carried out according to previous work (69), except that random hexamers were used for amplification. For a list of primers, see Supplementary table S7. Quantitative RT-PCR was performed in a Rotor-Gene RG-3000 (Corbett Science) under the following conditions: initial denaturation, 10 min at 95°C followed by 45 cycles of amplification (20 sec at 95°C, 20 sec at 60°C, 30 sec at 72°C). Each PCR reaction was performed in technical duplicate, differing by less than 5% between each other; the experiments were repeated at least three times with RNA isolated from independent cultures. To ensure that no primer-dimers were present, a melting curve was followed for each PCR. The results were normalized against 18S rRNA [30].

### Transcriptomics and transcriptome de novo assembly

RNA from cells prior to CP (pre-CP), at the time when approximately 50 % of them reached the first CP (CP) and when approximately 50 % of the cells divided nuclei into two (M) was isolated the same way as for RT-qPCR. For each sample, three biological replicates were analysed. RNA quality was checked using Agilent Bioanalyzer and only high quality RNA was further processed. Libraries were prepared from 1 μg of total RNA using NEBNext Ultra™ Directional RNA Library Prep Kit for Illumina (New England Biolabs, Ipswich, MA, USA) by poly(A) enrichment and pair-end sequenced using Illumina HiSeq 3000/4000 with read length 2 × 150 nt. Raw reads were adapter trimmed using CLC Genomics Workbench and reads from all conditions were used for *de novo* RNA assembly using the same software.

### Lipid gene metabolism identification

The lipid metabolism genes were identified by BLASTN homology based search (70) with protein sequences of *Chlamydomonas reinhardtii* homologues for *EPT1*, CDP-Ethanolamine:DAG Ethanolamine phosphotransferase (Cre12.g538450), *PGP3*, CDP-diacylglycerol-glycerol-3-phosphate 3-phosphatidyltransferase / Phosphatidylglycerophosphate synthase (Cre03.g162600) and *PIS1*, Phosphatidylinositol synthase (Cre10.g419800) used as baits. The protein sequences identified were verified by reciprocal BLAST against *Chlamydomonas reinhardtii* genome ver. 5.5 at Phytozome 12 (https://phytozome.jgi.doe.gov). A single homologue for *PIS1* and *EPT1* was identified, and two homologues of *PGP3*, designated *PGS1* and *PGS2*.

### Statistical tests

Univariate statistical tests were done using Excel 2013 or 2016. Graphs were produced in OriginLab2018. The integrations of each resonance (as a fraction of the total for that spectrum) were used to calculate the means, standard deviations and student’s *t*-tests reported for the abundance of lipids. The calculations were based on *n* = 6 biological replicates.

## Acknowledgements

We would like to thank Dr F. Kryuchkov for making the mass spectrometry instruments available and for training MJ. SF would like to thank Dr S. Liddell for helpful discussions. KB would like to thank Dr. M. Tichý for help with transcriptome assembly. Funding from the RCN (grant application ES528542), BBSRC (BB/M027252/1 and BB/T014210/1) and the National Programme of Sustainability I, ID: LO1416 is gratefully acknowledged.

## Author Contributions

**MV** co-designed the study, collected and analysed cell cycle data, advised on species type, co-developed handling procedures, designed experiments, supervised VL and co-wrote the manuscript. **VL** grew cultures, collected cells, acquired cell cycle data, prepared dried cell pellets and co-developed handling procedures. **MČ** ran all western blots and collected all qRT-PCR data. **MJ** acquired and converted all MS data and assisted in forging the collaboration. **FR** made the NMR instrument available, configured parameters of NMR experiments, collected all NMR data and ensured its quality. **KB** wrote the grant proposal that funded MV, VL and MČ, prepared and analysed the transcriptome and identified homologues of lipid metabolism genes, designed RT-qPCR and Western blot experiments, and co-wrote the manuscript**. ØH** wrote the original grant proposal that funded FR, SF and MJ, made equipment available, ensured data quality and. **SF** conceived the hypothesis, designed the study, extracted lipids and prepared all samples for lipid profiling, developed methods, analysed NMR and MS data and co-wrote the manuscript. All authors commented on the manuscript and approved the final version.

## Tables

**Table 1.** The collection points of cultures through the cell cycle of Desmodesmus quadricauda *used in this study.*

| Phase | Collection time after Starting culture |
| --- | --- |
| Early G<sub>1</sub> | +2 h |
| Mid G<sub>1</sub>, the first CP | +4 h |
| Late G<sub>1</sub>, pS | +6 h |
| S, the second CP | +8 h |
| G<sub>2</sub> | +10 h |
| M | +13 h |
| G<sub>3</sub> | +15 h |

## Supplementary Information

For Vítová *et al.*, The biosynthesis of phospholipids is linked to the cell cycle in a model eukaryote.

**Fig. S1.**
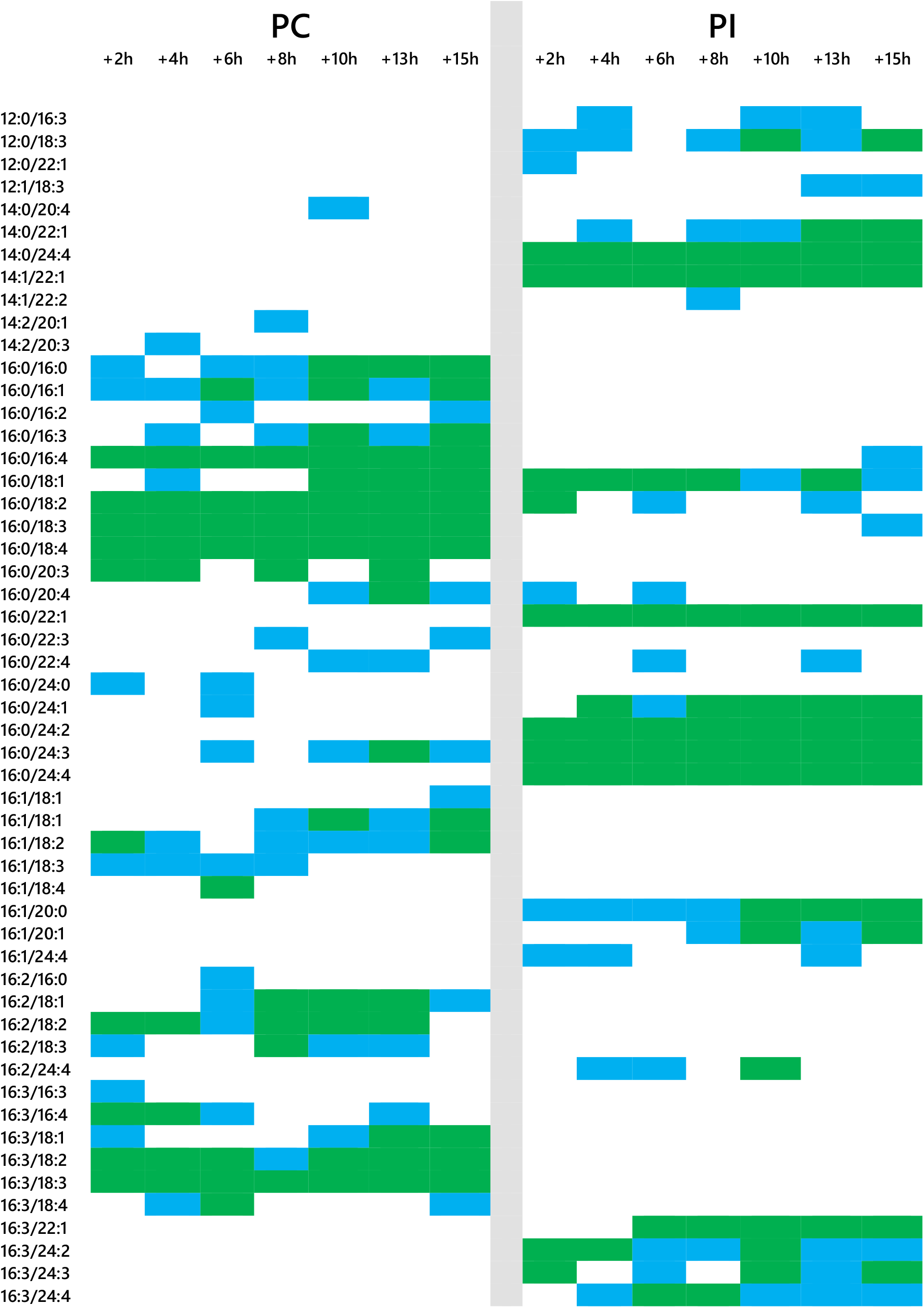
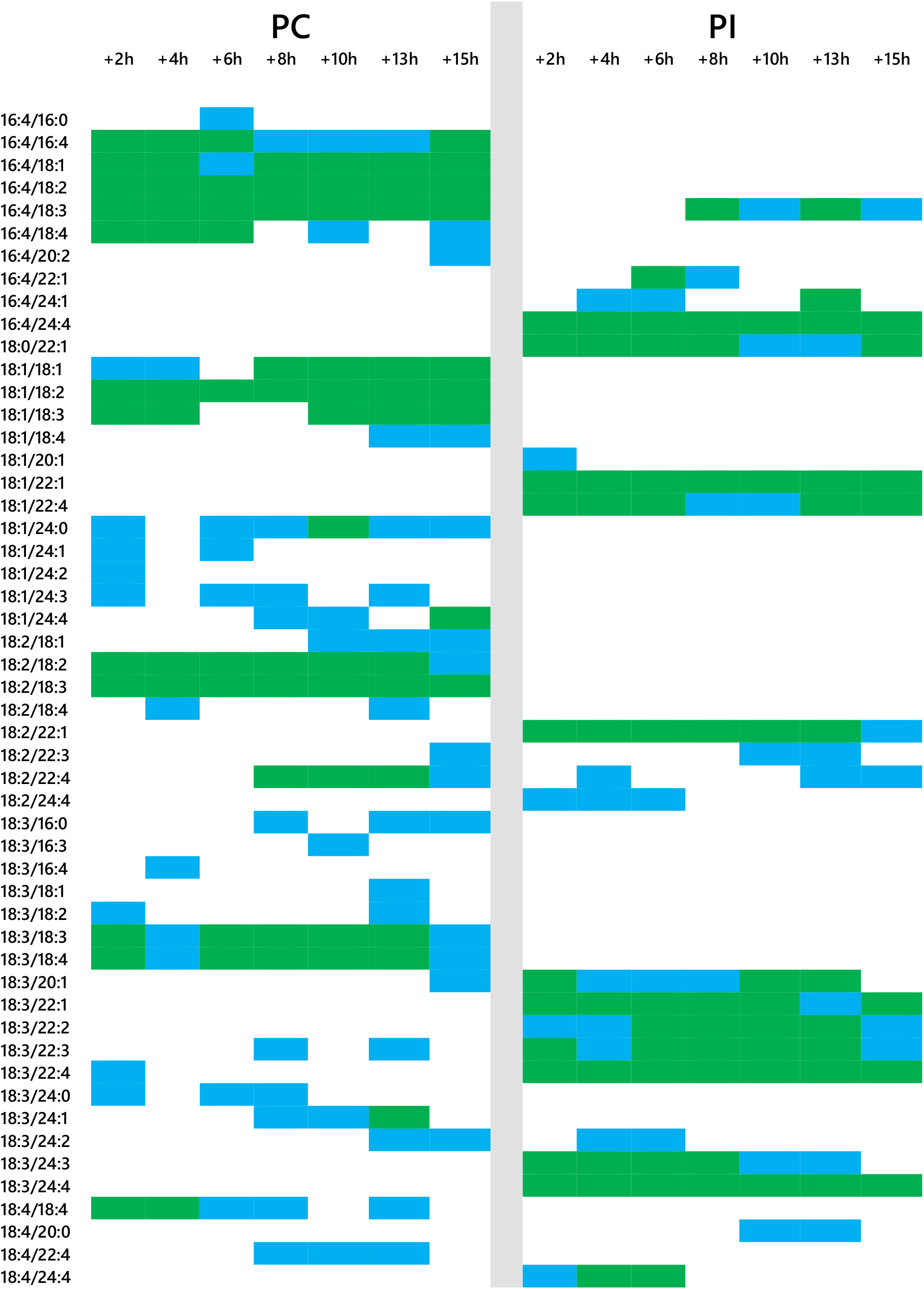
Matrix showing the frequency of the detection of isoforms of PC (left) and PI (right) in the samples used in the present study, through the cell cycle of Desmodesmus quadricauda, using mass spectrometry. White indicates that the isoform was not found in any of the samples, green indicates that that isoform was found in all samples tested and blue that it was found in one sample. Collection points: +2 h, early part of G_1_; +4 h, mid-G_1_ (first CP); +6 h, end of G_1_ and pS; +8 h, S and second CP; +10 h, G_2_; +13 h, M; +15 h, G_3_, for others see Figure 1.

**Fig. S2.**
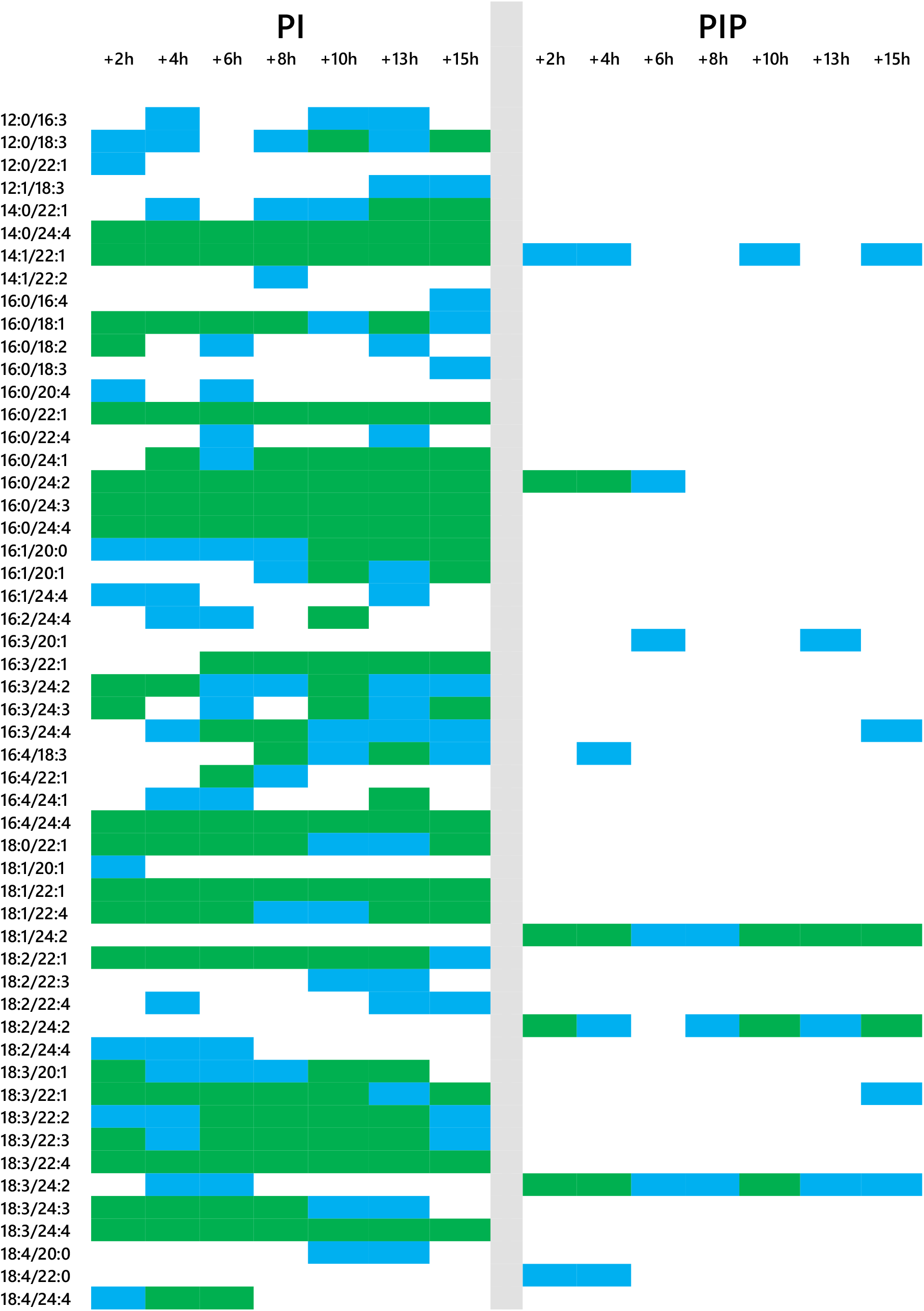
Matrices showing the isoform profiles of PI and PIPs from Desmodesmus quadricauda in the samples used in the present study, through the cell cycle of Desmodesmus quadricauda, using mass spectrometry. White indicates that the isoform was not found in any of the samples, green indicates that that isoform was found in all samples tested and blue that it was found in one sample. Collection points: +2 h, early part of G_1_; +4 h, mid-G_1_ (first CP); +6 h, end of G_1_ and pS; +8 h, S and second CP; +10 h, G_2_; +13 h, M; +15 h, G_3_, for others see Figure 1.

**Fig. S3.**
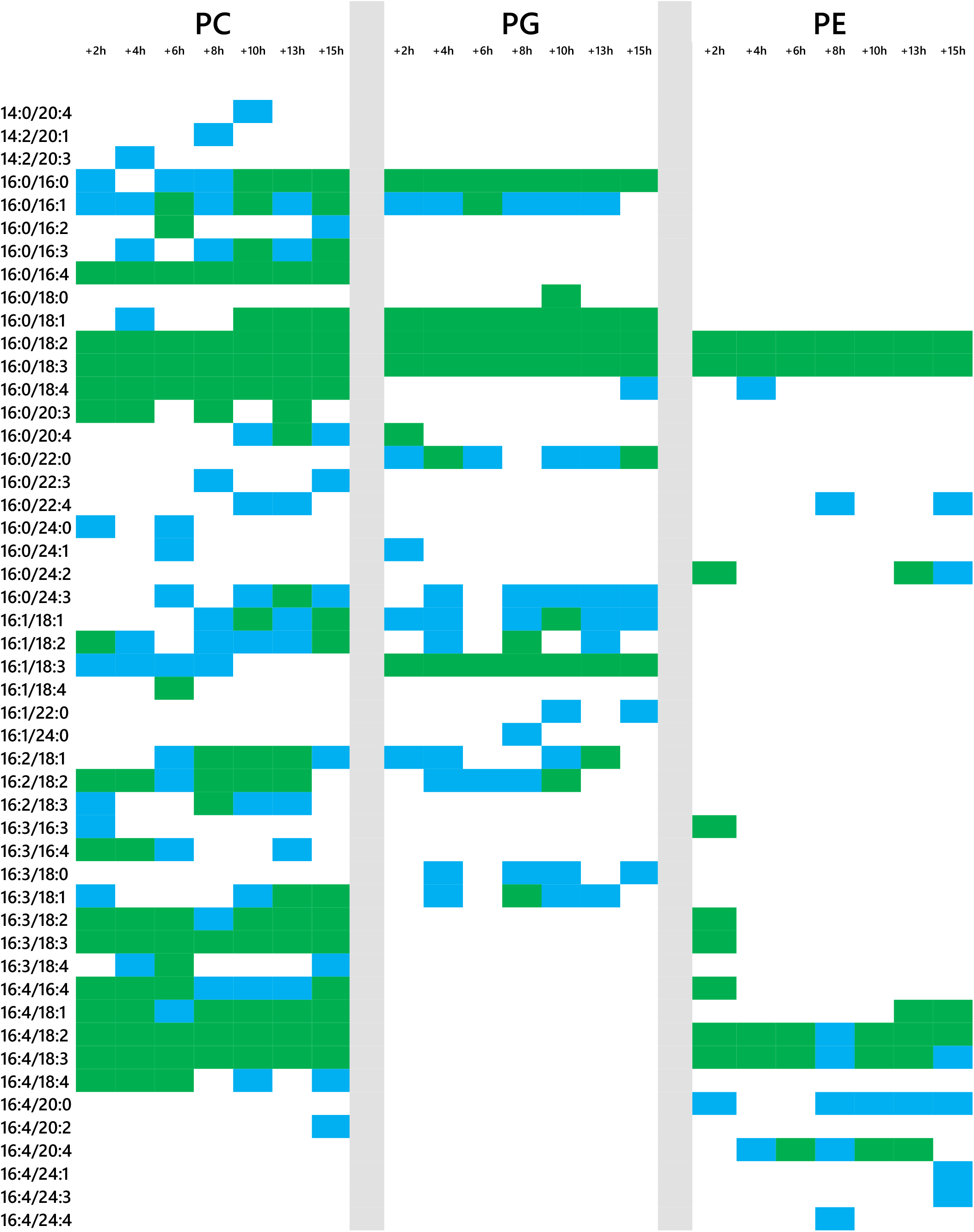
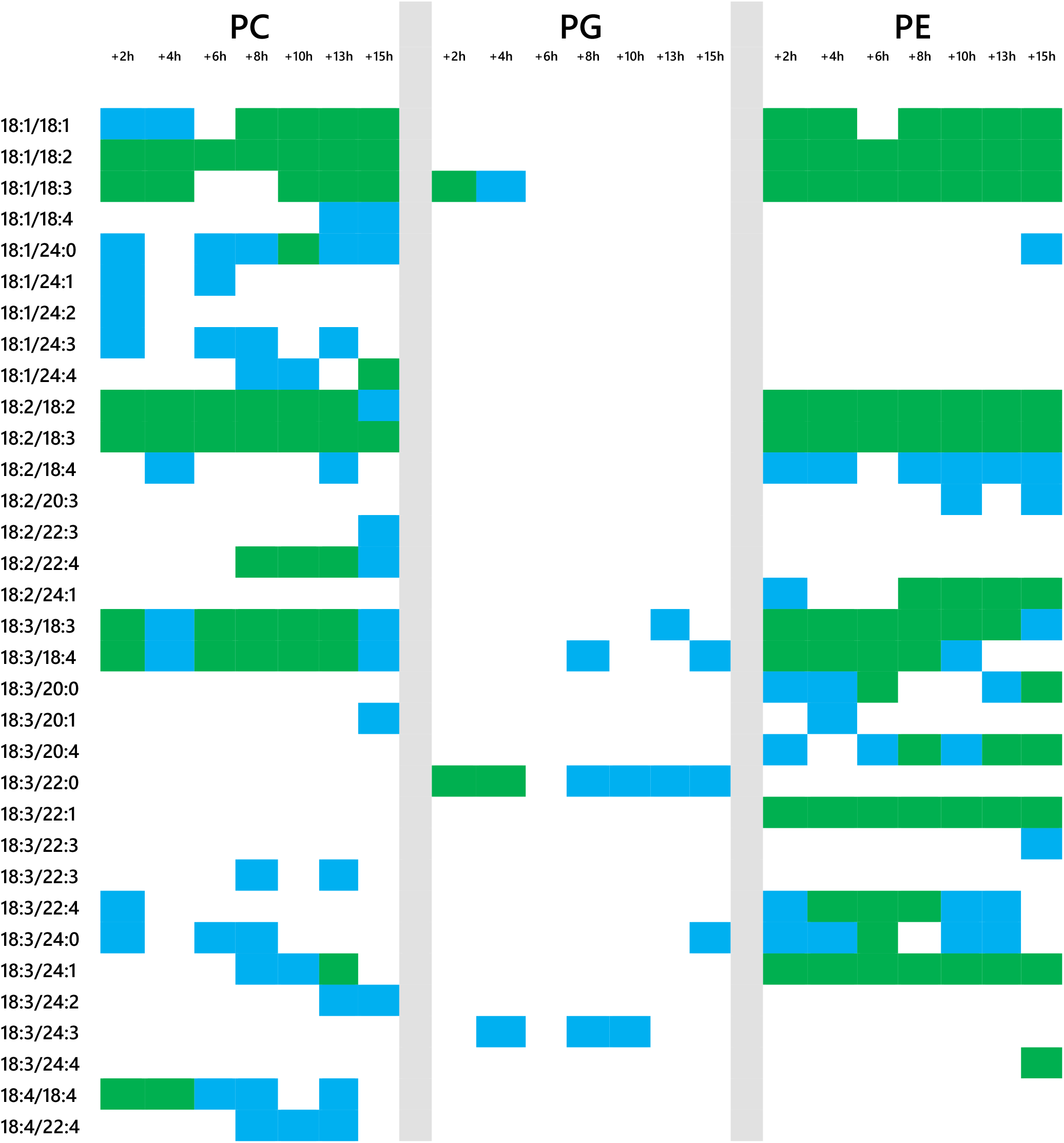
Matrix showing the frequency of the detection of isoforms of PC, PG and PE in the samples used in the present study, through the cell cycle of Desmodesmus quadricauda, using mass spectrometry. White indicates that the isoform was not found in any of the samples, green indicates that that isoform was found in all samples tested and blue that it was found in one sample. Collection points: +2 h, early part of G_1_; +4 h, mid-G_1_ (first CP); +6 h, end of G_1_ and pS; +8 h, S and second CP; +10 h, G_2_; +13 h, M; +15 h, G_3_, for others see Figure 1.

**Fig. S4.**
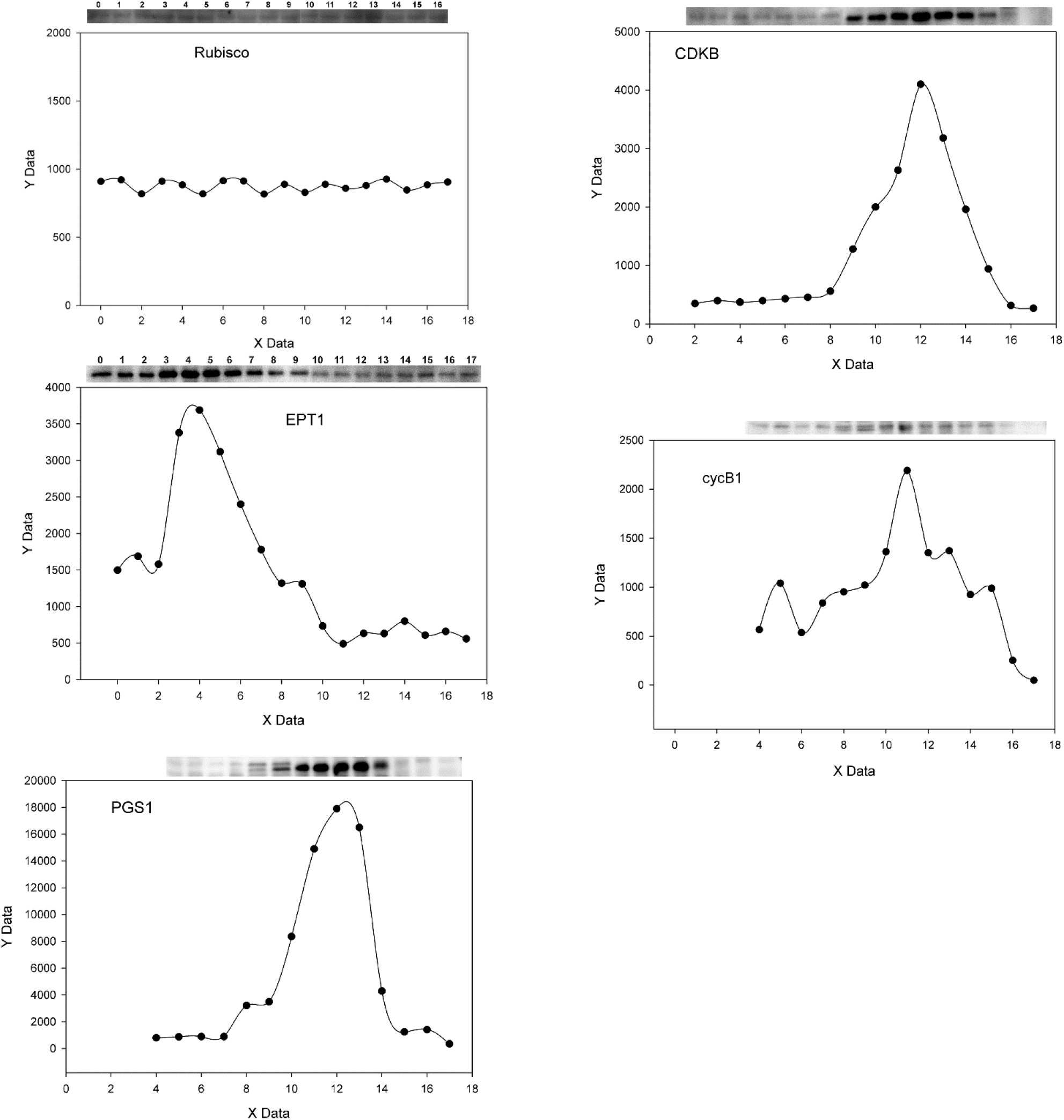
Western blots and accompanying plots of relative abundance of ethanolamine phosphate transferase (EPT1) and phosphatidylglycerol synthase (PGS1) through a typical cell cycle (as demonstrated by the abundance of CDKB and cycB1) and against a steady rate of protein synthesis (as demonstrated by the abundance of constitutive protein RuBISCo). X Data = time (h); Y Data = relative abundance (a.u.).

## Supplementary Tables

**Table S1.**
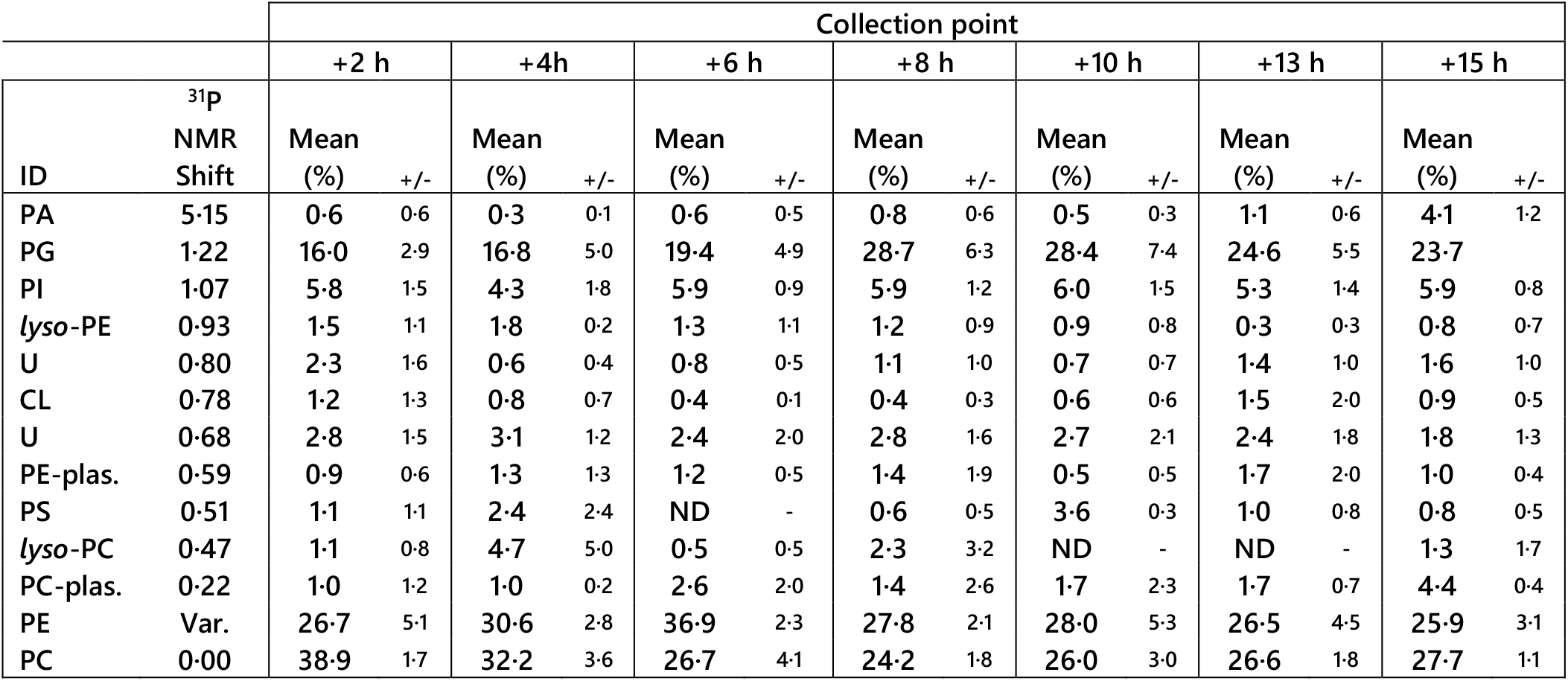
The lipid head group profile of D. quadricauda determined using ^31^P NMR. ^31^P assignments made using literature references (28–30). CL, cardiolipin; PA, phosphatidic acid; PC, phosphatidylcholine; PE, phosphatidylethanolamine; PG, phosphatidylglycerol; PI, phosphatidylinositol; PS, phosphatidylserine; U, unidentified. Collection points: +2 h, early part of G_1_; +4 h, mid-G_1_ (first CP); +6 h, end of G_1_ and pS; +8 h, S and second CP; +10 h, G_2_; +13 h, M; +15 h, G_3_, for others see Figure 1.

**Table S2.**
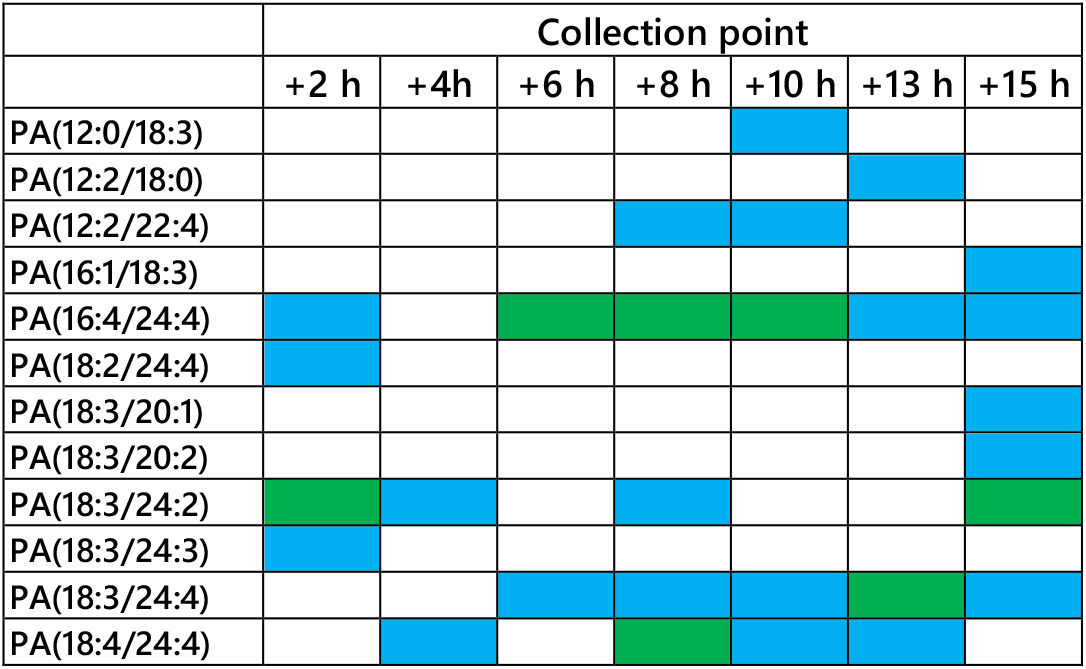
The Isoform profile of phosphatidic acids from D. quadricauda determined using MS/MS. White indicates not found, blue indicates found in one or more samples, green indicates found in all samples tested. Collection points: +2 h, early part of G_1_; +4 h, mid-G_1_ (first CP); +6 h, end of G_1_ and pS; +8 h, S and second CP; +10 h, G_2_; +13 h, M; +15 h, G_3_, for others see Figure 1.

**Table S3.**
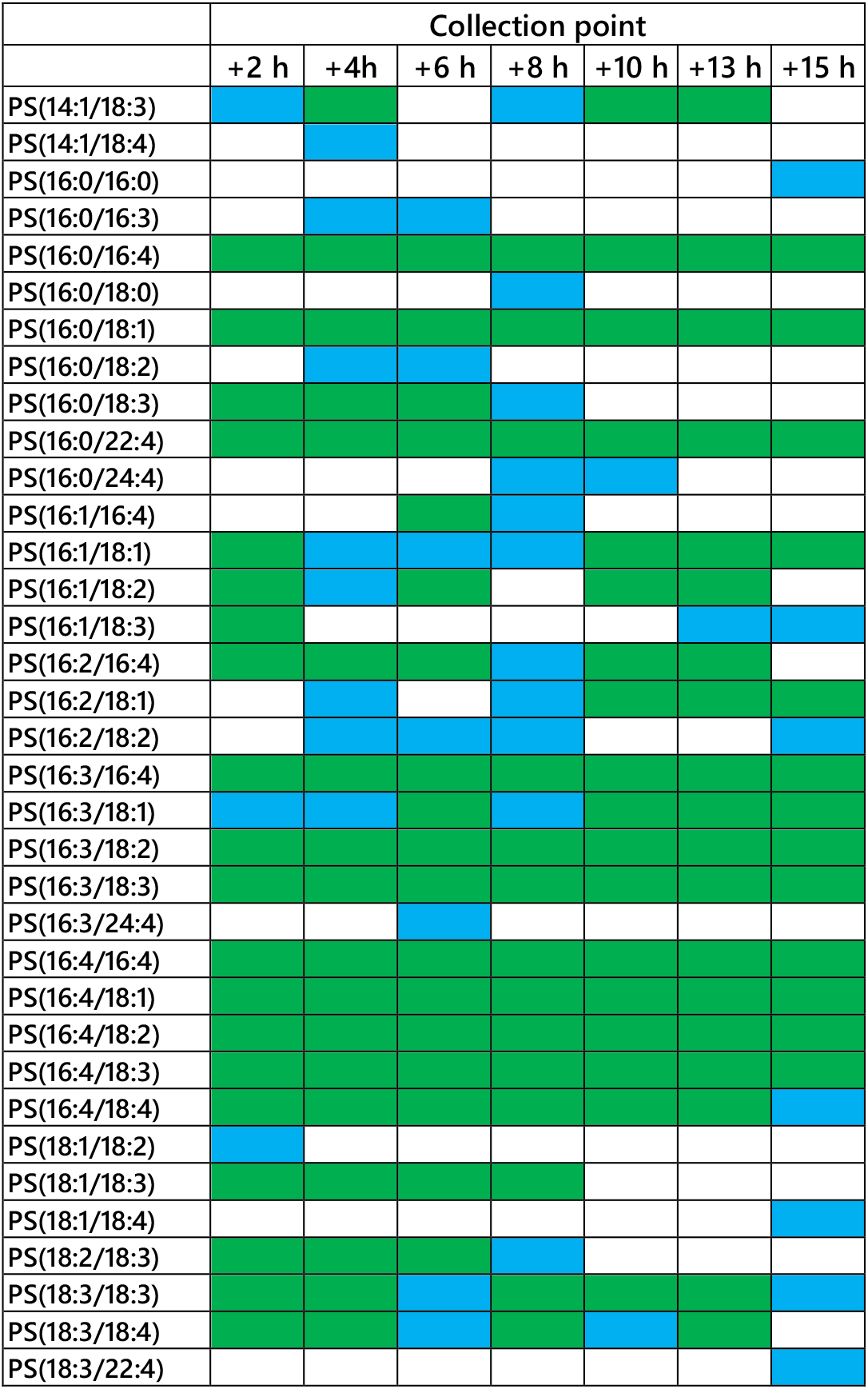
The Isoform profile of phosphatidylserines from D. quadricauda determined using MS/MS. White indicates not found, blue indicates found in one sample, green indicates found in both samples tested. Collection points: +2 h, early part of G_1_; +4 h, mid-G_1_ (first CP); +6 h, end of G_1_ and pS; +8 h, S and second CP; +10 h, G_2_; +13 h, M; +15 h, G_3_, for others see Figure 1.

**Table S4.**
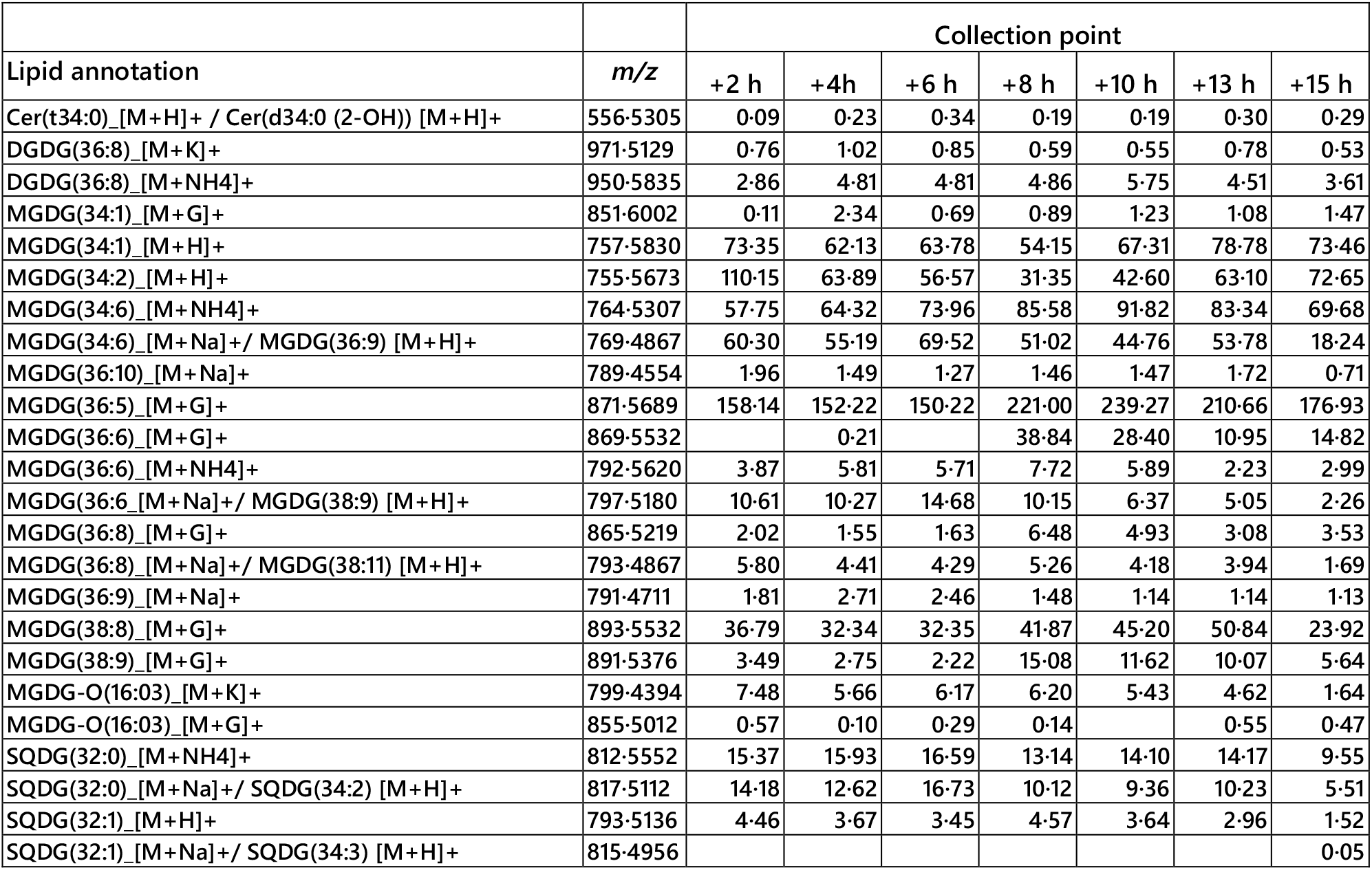
Signal intensities for high resolution m/z values consistent with isoforms of galactosyl-diglycerides and one phosphatidylinositol-ceramide (n = 2). Signals were assigned according to high resolution masses with a deviation of <12 ppm from the calculated monoisotopic mass and checked with a visual inspection of a subset of the original chromatograms. CE, Campestryl ester; G, guanidine; MGDG, mono-galactosyl diglyceride; MGDG-O, mono-galactosyl diglyceride oxide; PI-Cer, phosphatidylinositol ceramide; SQDG, sulfoquinovosyl diglyceride; TG, triglyceride. Collection points: +2 h, early part of G_1_; +4 h, mid-G_1_ (first CP); +6 h, end of G_1_ and pS; +8 h, S and second CP; +10 h, G_2_; +13 h, M; +15 h, G_3_, for others see Figure 1.

**Table S5.**
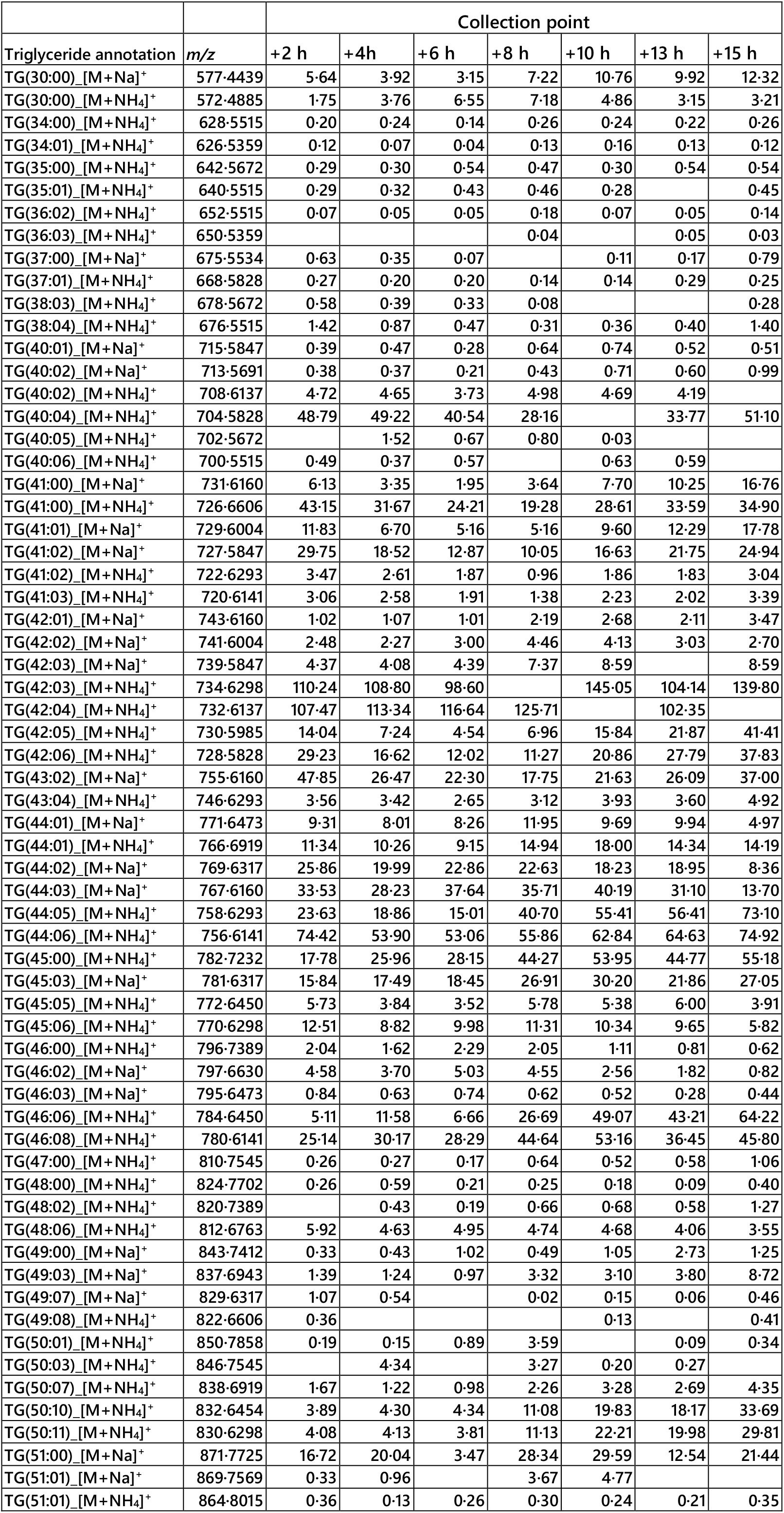
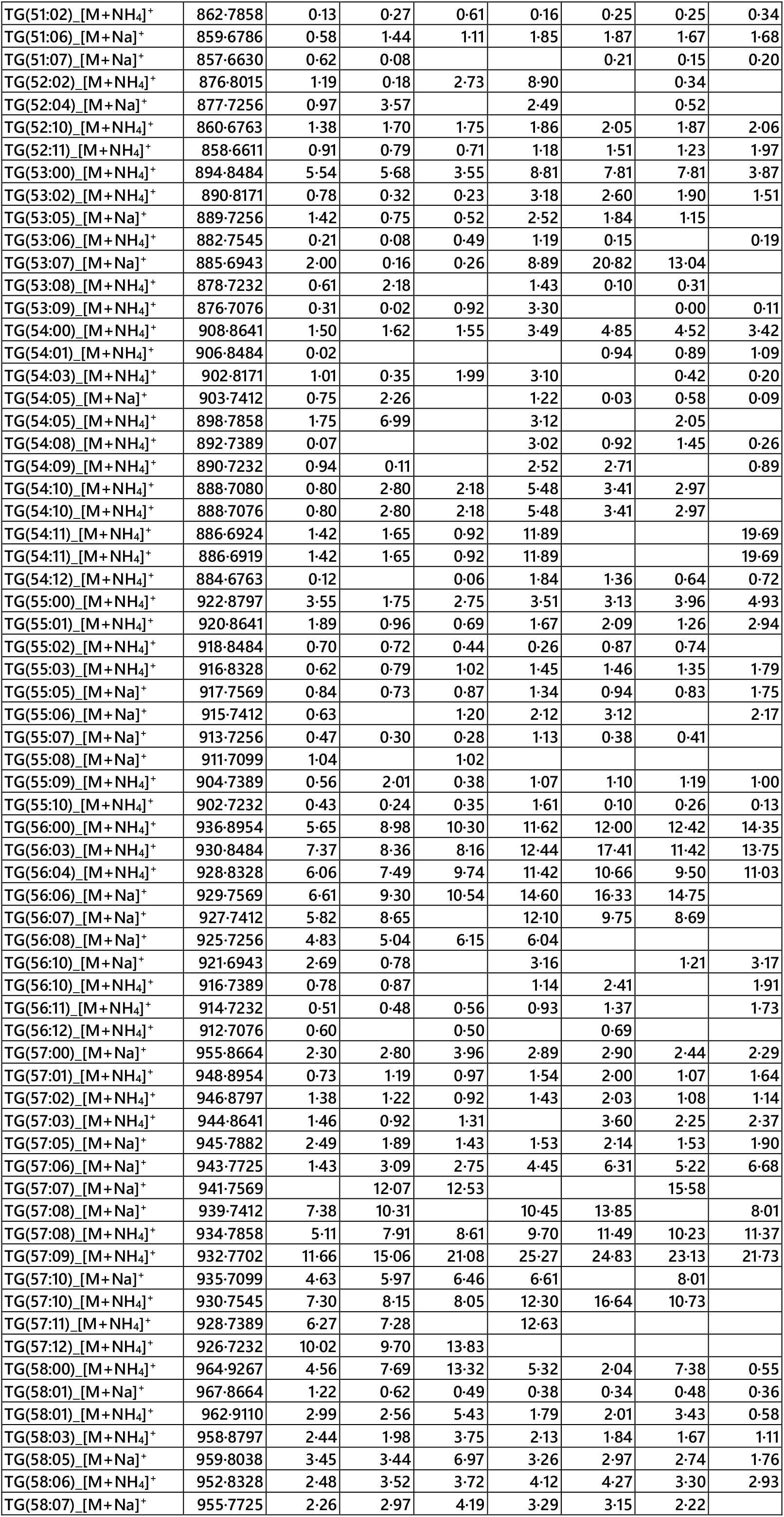
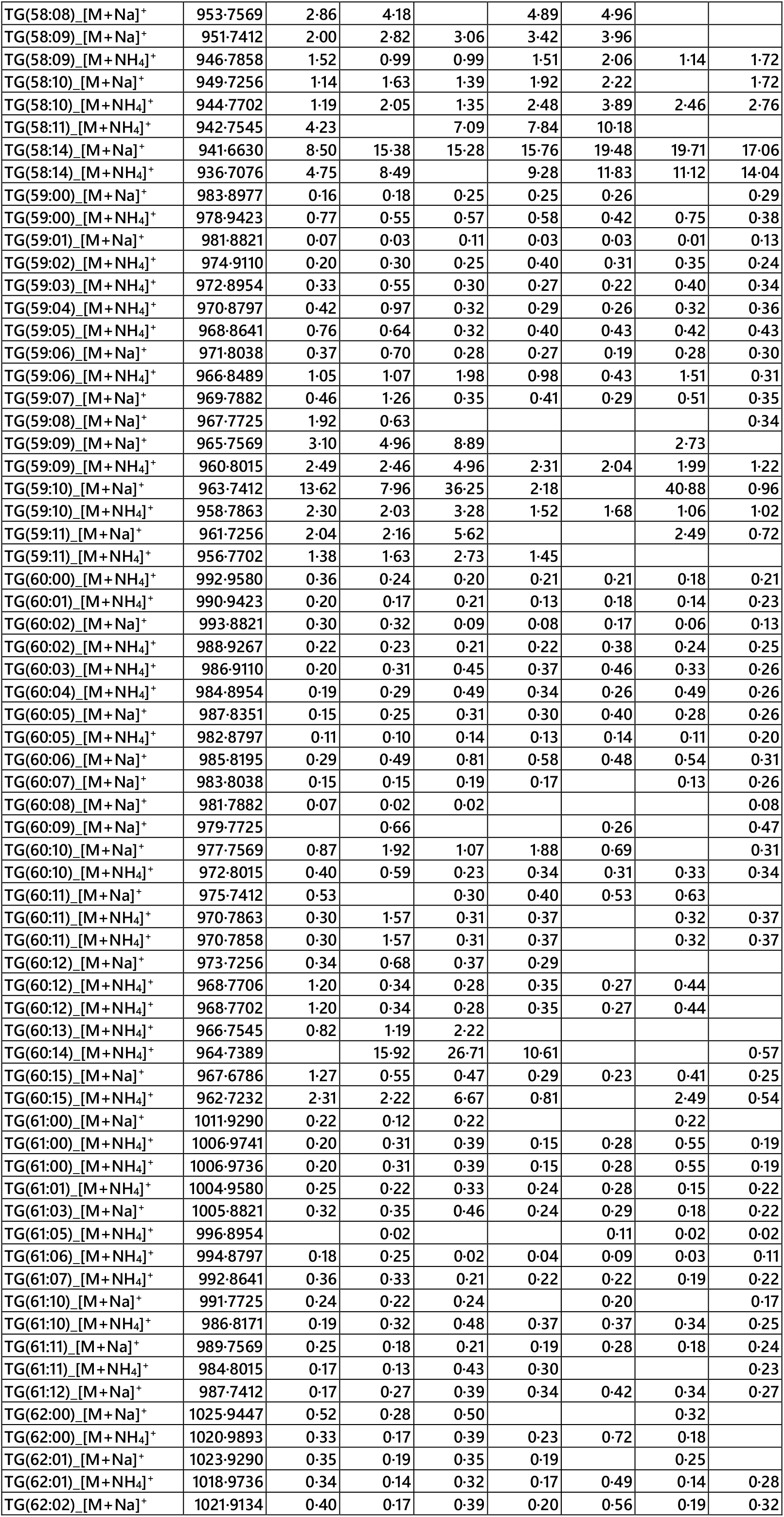
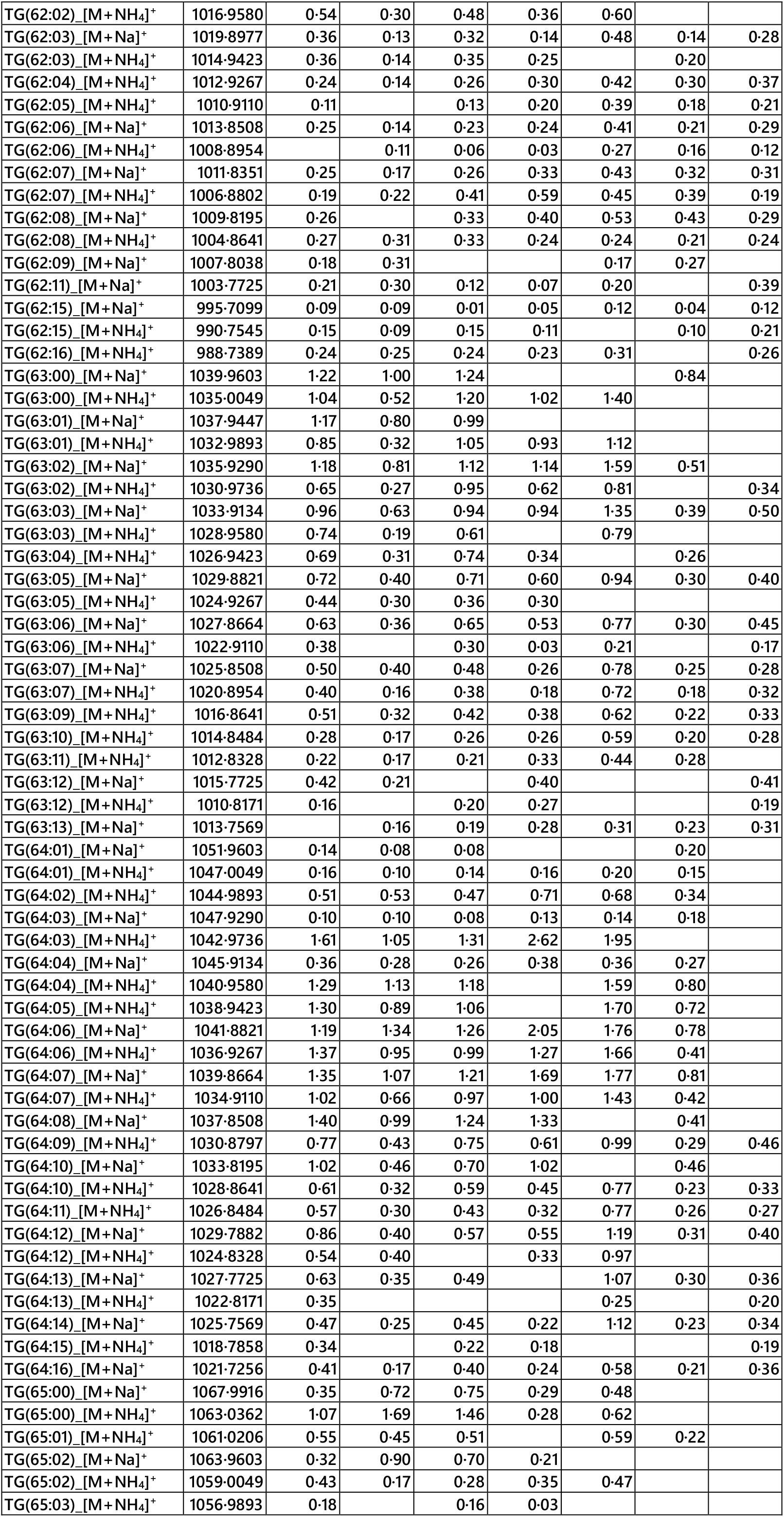
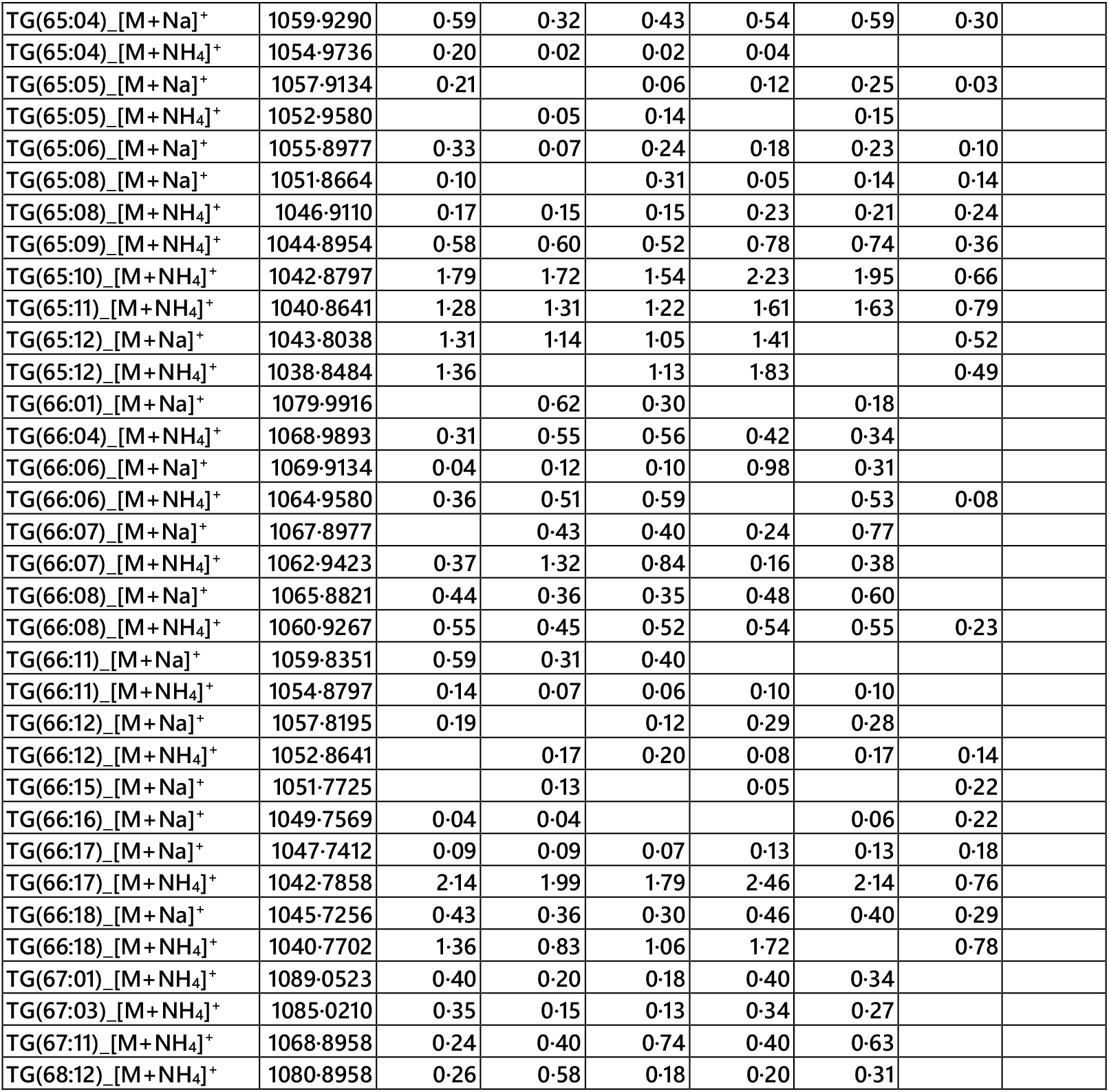
Signal intensities for high resolution m/z values consistent with triglyceride isoforms (n = 2). Signals were assigned according to high resolution masses with a deviation of <12 ppm from the calculated monoisotopic mass. Quantitative standards were not used. G, guanidinium. Collection points: +2 h, early part of G_1_; +4 h, mid-G_1_ (first CP); +6 h, end of G_1_ and pS; +8 h, S and second CP; +10 h, G_2_; +13 h, M; +15 h, G_3_, for others see Figure 1.

| Triglyceride annotation | m/z | Collection point |  |  |  |  |  |  |
| --- | --- | --- | --- | --- | --- | --- | --- | --- |
|  |  | +2 h | +4h | +6 h | +8 h | +10 h | +13 h | +15 h |
| TG(30:00) [M+Na]<sup>+</sup> | 577-4439 | 5-64 | 3-92 | 3-15 | 7-22 | 10-76 | 9-92 | 12-32 |
| TG(30:00) [M+NH<sub>4</sub>]<sup>+</sup> | 572-4885 | 1-75 | 3-76 | 6-55 | 7-18 | 4-86 | 3-15 | 3-21 |
| TG(34:00) [M+NH<sub>4</sub>]<sup>+</sup> | 628-5515 | 0-20 | 0-24 | 0-14 | 0-26 | 0-24 | 0-22 | 0-26 |
| TG(34:01) [M+NH<sub>4</sub>]<sup>+</sup> | 626-5359 | 0-12 | 0-07 | 0-04 | 0-13 | 0-16 | 0-13 | 0-12 |
| TG(35:00) [M+NH<sub>4</sub>]<sup>+</sup> | 642-5672 | 0-29 | 0-30 | 0-54 | 0-47 | 0-30 | 0-54 | 0-54 |
| TG(35:01) [M+NH<sub>4</sub>]<sup>+</sup> | 640-5515 | 0-29 | 0-32 | 0-43 | 0-46 | 0-28 |  | 0-45 |
| TG(36:02) [M+NH<sub>4</sub>]<sup>+</sup> | 652-5515 | 0-07 | 0-05 | 0-05 | 0-18 | 0-07 | 0-05 | 0-14 |
| TG(36:03) [M+NH<sub>4</sub>]<sup>+</sup> | 650-5359 |  |  |  | 0-04 |  | 0-05 | 0-03 |
| TG(37:00) [M+Na]<sup>+</sup> | 675-5534 | 0-63 | 0-35 | 0-07 |  | 0-11 | 0-17 | 0-79 |
| TG(37:01) [M+NH<sub>4</sub>]<sup>+</sup> | 668-5828 | 0-27 | 0-20 | 0-20 | 0-14 | 0-14 | 0-29 | 0-25 |
| TG(38:03) [M+NH<sub>4</sub>]<sup>+</sup> | 678-5672 | 0-58 | 0-39 | 0-33 | 0-08 |  |  | 0-28 |
| TG(38:04) [M+NH<sub>4</sub>]<sup>+</sup> | 676-5515 | 1-42 | 0-87 | 0-47 | 0-31 | 0-36 | 0-40 | 1-40 |
| TG(40:01) [M+Na]<sup>+</sup> | 715-5847 | 0-39 | 0-47 | 0-28 | 0-64 | 0-74 | 0-52 | 0-51 |
| TG(40:02) [M+Na]<sup>+</sup> | 713-5691 | 0-38 | 0-37 | 0-21 | 0-43 | 0-71 | 0-60 | 0-99 |
| TG(40:02) [M+NH<sub>4</sub>]<sup>+</sup> | 708-6137 | 4-72 | 4-65 | 3-73 | 4-98 | 4-69 | 4-19 |  |
| TG(40:04) [M+NH<sub>4</sub>]<sup>+</sup> | 704-5828 | 48-79 | 49-22 | 40-54 | 28-16 |  | 33-77 | 51-10 |
| TG(40:05) [M+NH<sub>4</sub>]<sup>+</sup> | 702-5672 |  | 1-52 | 0-67 | 0-80 | 0-03 |  |  |
| TG(40:06) [M+NH<sub>4</sub>]<sup>+</sup> | 700-5515 | 0-49 | 0-37 | 0-57 |  | 0-63 | 0-59 |  |
| TG(41:00) [M+Na]<sup>+</sup> | 731-6160 | 6-13 | 3-35 | 1-95 | 3-64 | 7-70 | 10-25 | 16-76 |
| TG(41:00) [M+NH<sub>4</sub>]<sup>+</sup> | 726-6606 | 43-15 | 31-67 | 24-21 | 19-28 | 28-61 | 33-59 | 34-90 |
| TG(41:01) [M+Na]<sup>+</sup> | 729-6004 | 11-83 | 6-70 | 5-16 | 5-16 | 9-60 | 12-29 | 17-78 |
| TG(41:02) [M+Na]<sup>+</sup> | 727-5847 | 29-75 | 18-52 | 12-87 | 10-05 | 16-63 | 21-75 | 24-94 |
| TG(41:02) [M+NH<sub>4</sub>]<sup>+</sup> | 722-6293 | 3-47 | 2-61 | 1-87 | 0-96 | 1-86 | 1-83 | 3-04 |
| TG(41:03) [M+NH<sub>4</sub>]<sup>+</sup> | 720-6141 | 3-06 | 2-58 | 1-91 | 1-38 | 2-23 | 2-02 | 3-39 |
| TG(42:01) [M+Na]<sup>+</sup> | 743-6160 | 1-02 | 1-07 | 1-01 | 2-19 | 2-68 | 2-11 | 3-47 |
| TG(42:02) [M+Na]<sup>+</sup> | 741-6004 | 2-48 | 2-27 | 3-00 | 4-46 | 4-13 | 3-03 | 2-70 |
| TG(42:03) [M+Na]<sup>+</sup> | 739-5847 | 4-37 | 4-08 | 4-39 | 7-37 | 8-59 |  | 8-59 |
| TG(42:03) [M+NH<sub>4</sub>]<sup>+</sup> | 734-6298 | 110-24 | 108-80 | 98-60 |  | 145-05 | 104-14 | 139-80 |
| TG(42:04) [M+NH<sub>4</sub>]<sup>+</sup> | 732-6137 | 107-47 | 113-34 | 116-64 | 125-71 |  | 102-35 |  |
| TG(42:05) [M+NH<sub>4</sub>]<sup>+</sup> | 730-5985 | 14-04 | 7-24 | 4-54 | 6-96 | 15-84 | 21-87 | 41-41 |
| TG(42:06) [M+NH<sub>4</sub>]<sup>+</sup> | 728-5828 | 29-23 | 16-62 | 12-02 | 11-27 | 20-86 | 27-79 | 37-83 |
| TG(43:02) [M+Na]<sup>+</sup> | 755-6160 | 47-85 | 26-47 | 22-30 | 17-75 | 21-63 | 26-09 | 37-00 |
| TG(43:04) [M+NH<sub>4</sub>]<sup>+</sup> | 746-6293 | 3-56 | 3-42 | 2-65 | 3-12 | 3-93 | 3-60 | 4-92 |
| TG(44:01) [M+Na]<sup>+</sup> | 771-6473 | 9-31 | 8-01 | 8-26 | 11-95 | 9-69 | 9-94 | 4-97 |
| TG(44:01) [M+NH<sub>4</sub>]<sup>+</sup> | 766-6919 | 11-34 | 10-26 | 9-15 | 14-94 | 18-00 | 14-34 | 14-19 |
| TG(44:02) [M+Na]<sup>+</sup> | 769-6317 | 25-86 | 19-99 | 22-86 | 22-63 | 18-23 | 18-95 | 8-36 |
| TG(44:03) [M+Na]<sup>+</sup> | 767-6160 | 33-53 | 28-23 | 37-64 | 35-71 | 40-19 | 31-10 | 13-70 |
| TG(44:05) [M+NH<sub>4</sub>]<sup>+</sup> | 758-6293 | 23-63 | 18-86 | 15-01 | 40-70 | 55-41 | 56-41 | 73-10 |
| TG(44:06) [M+NH<sub>4</sub>]<sup>+</sup> | 756-6141 | 74-42 | 53-90 | 53-06 | 55-86 | 62-84 | 64-63 | 74-92 |
| TG(45:00) [M+NH<sub>4</sub>]<sup>+</sup> | 782-7232 | 17-78 | 25-96 | 28-15 | 44-27 | 53-95 | 44-77 | 55-18 |
| TG(45:03) [M+Na]<sup>+</sup> | 781-6317 | 15-84 | 17-49 | 18-45 | 26-91 | 30-20 | 21-86 | 27-05 |
| TG(45:05) [M+NH<sub>4</sub>]<sup>+</sup> | 772-6450 | 5-73 | 3-84 | 3-52 | 5-78 | 5-38 | 6-00 | 3-91 |
| TG(45:06) [M+NH<sub>4</sub>]<sup>+</sup> | 770-6298 | 12-51 | 8-82 | 9-98 | 11-31 | 10-34 | 9-65 | 5-82 |
| TG(46:00) [M+NH<sub>4</sub>]<sup>+</sup> | 796-7389 | 2-04 | 1-62 | 2-29 | 2-05 | 1-11 | 0-81 | 0-62 |
| TG(46:02) [M+Na]<sup>+</sup> | 797-6630 | 4-58 | 3-70 | 5-03 | 4-55 | 2-56 | 1-82 | 0-82 |
| TG(46:03) [M+Na]<sup>+</sup> | 795-6473 | 0-84 | 0-63 | 0-74 | 0-62 | 0-52 | 0-28 | 0-44 |
| TG(46:06) [M+NH<sub>4</sub>]<sup>+</sup> | 784-6450 | 5-11 | 11-58 | 6-66 | 26-69 | 49-07 | 43-21 | 64-22 |
| TG(46:08) [M+NH<sub>4</sub>]<sup>+</sup> | 780-6141 | 25-14 | 30-17 | 28-29 | 44-64 | 53-16 | 36-45 | 45-80 |
| TG(47:00) [M+NH<sub>4</sub>]<sup>+</sup> | 810-7545 | 0-26 | 0-27 | 0-17 | 0-64 | 0-52 | 0-58 | 1-06 |
| TG(48:00) [M+NH<sub>4</sub>]<sup>+</sup> | 824-7702 | 0-26 | 0-59 | 0-21 | 0-25 | 0-18 | 0-09 | 0-40 |
| TG(48:02) [M+NH<sub>4</sub>]<sup>+</sup> | 820-7389 |  | 0-43 | 0-19 | 0-66 | 0-68 | 0-58 | 1-27 |
| TG(48:06) [M+NH<sub>4</sub>]<sup>+</sup> | 812-6763 | 5-92 | 4-63 | 4-95 | 4-74 | 4-68 | 4-06 | 3-55 |
| TG(49:00) [M+Na]<sup>+</sup> | 843-7412 | 0-33 | 0-43 | 1-02 | 0-49 | 1-05 | 2-73 | 1-25 |
| TG(49:03) [M+Na]<sup>+</sup> | 837-6943 | 1-39 | 1-24 | 0-97 | 3-32 | 3-10 | 3-80 | 8-72 |
| TG(49:07) [M+Na]<sup>+</sup> | 829-6317 | 1-07 | 0-54 |  | 0-02 | 0-15 | 0-06 | 0-46 |
| TG(49:08) [M+NH<sub>4</sub>]<sup>+</sup> | 822-6606 | 0-36 |  |  |  | 0-13 |  | 0-41 |
| TG(50:01) [M+NH<sub>4</sub>]<sup>+</sup> | 850-7858 | 0-19 | 0-15 | 0-89 | 3-59 |  | 0-09 | 0-34 |
| TG(50:03) [M+NH<sub>4</sub>]<sup>+</sup> | 846-7545 |  | 4-34 |  | 3-27 | 0-20 | 0-27 |  |
| TG(50:07) [M+NH<sub>4</sub>]<sup>+</sup> | 838-6919 | 1-67 | 1-22 | 0-98 | 2-26 | 3-28 | 2-69 | 4-35 |
| TG(50:10) [M+NH<sub>4</sub>]<sup>+</sup> | 832-6454 | 3-89 | 4-30 | 4-34 | 11-08 | 19-83 | 18-17 | 33-69 |
| TG(50:11) [M+NH<sub>4</sub>]<sup>+</sup> | 830-6298 | 4-08 | 4-13 | 3-81 | 11-13 | 22-21 | 19-98 | 29-81 |
| TG(51:00) [M+Na]<sup>+</sup> | 871-7725 | 16-72 | 20-04 | 3-47 | 28-34 | 29-59 | 12-54 | 21-44 |
| TG(51:01) [M+Na]<sup>+</sup> | 869-7569 | 0-33 | 0-96 |  | 3-67 | 4-77 |  |  |
| TG(51:01) [M+NH<sub>4</sub>]<sup>+</sup> | 864-8015 | 0-36 | 0-13 | 0-26 | 0-30 | 0-24 | 0-21 | 0-35 |
| TG(51:02)_[M+NH<sub>4</sub>]<sup>+</sup> | 862-7858 | 0-13 | 0-27 | 0-61 | 0-16 | 0-25 | 0-25 | 0-34 |
| TG(51:06)_[M+Na]<sup>+</sup> | 859-6786 | 0-58 | 1-44 | 1-11 | 1-85 | 1-87 | 1-67 | 1-68 |
| TG(51:07)_[M+Na]<sup>+</sup> | 857-6630 | 0-62 | 0-08 |  |  | 0-21 | 0-15 | 0-20 |
| TG(52:02)_[M+NH<sub>4</sub>]<sup>+</sup> | 876-8015 | 1-19 | 0-18 | 2-73 | 8-90 |  | 0-34 |  |
| TG(52:04)_[M+Na]<sup>+</sup> | 877-7256 | 0-97 | 3-57 |  | 2-49 |  | 0-52 |  |
| TG(52:10)_[M+NH<sub>4</sub>]<sup>+</sup> | 860-6763 | 1-38 | 1-70 | 1-75 | 1-86 | 2-05 | 1-87 | 2-06 |
| TG(52:11)_[M+NH<sub>4</sub>]<sup>+</sup> | 858-6611 | 0-91 | 0-79 | 0-71 | 1-18 | 1-51 | 1-23 | 1-97 |
| TG(53:00)_[M+NH<sub>4</sub>]<sup>+</sup> | 894-8484 | 5-54 | 5-68 | 3-55 | 8-81 | 7-81 | 7-81 | 3-87 |
| TG(53:02)_[M+NH<sub>4</sub>]<sup>+</sup> | 890-8171 | 0-78 | 0-32 | 0-23 | 3-18 | 2-60 | 1-90 | 1-51 |
| TG(53:05)_[M+Na]<sup>+</sup> | 889-7256 | 1-42 | 0-75 | 0-52 | 2-52 | 1-84 | 1-15 |  |
| TG(53:06)_[M+NH<sub>4</sub>]<sup>+</sup> | 882-7545 | 0-21 | 0-08 | 0-49 | 1-19 | 0-15 |  | 0-19 |
| TG(53:07)_[M+Na]<sup>+</sup> | 885-6943 | 2-00 | 0-16 | 0-26 | 8-89 | 20-82 | 13-04 |  |
| TG(53:08)_[M+NH<sub>4</sub>]<sup>+</sup> | 878-7232 | 0-61 | 2-18 |  | 1-43 | 0-10 | 0-31 |  |
| TG(53:09)_[M+NH<sub>4</sub>]<sup>+</sup> | 876-7076 | 0-31 | 0-02 | 0-92 | 3-30 |  | 0-00 | 0-11 |
| TG(54:00)_[M+NH<sub>4</sub>]<sup>+</sup> | 908-8641 | 1-50 | 1-62 | 1-55 | 3-49 | 4-85 | 4-52 | 3-42 |
| TG(54:01)_[M+NH<sub>4</sub>]<sup>+</sup> | 906-8484 | 0-02 |  |  |  | 0-94 | 0-89 | 1-09 |
| TG(54:03)_[M+NH<sub>4</sub>]<sup>+</sup> | 902-8171 | 1-01 | 0-35 | 1-99 | 3-10 |  | 0-42 | 0-20 |
| TG(54:05)_[M+Na]<sup>+</sup> | 903-7412 | 0-75 | 2-26 |  | 1-22 | 0-03 | 0-58 | 0-09 |
| TG(54:05)_[M+NH<sub>4</sub>]<sup>+</sup> | 898-7858 | 1-75 | 6-99 |  | 3-12 |  | 2-05 |  |
| TG(54:08)_[M+NH<sub>4</sub>]<sup>+</sup> | 892-7389 | 0-07 |  |  | 3-02 | 0-92 | 1-45 | 0-26 |
| TG(54:09)_[M+NH<sub>4</sub>]<sup>+</sup> | 890-7232 | 0-94 | 0-11 |  | 2-52 | 2-71 |  | 0-89 |
| TG(54:10)_[M+NH<sub>4</sub>]<sup>+</sup> | 888-7080 | 0-80 | 2-80 | 2-18 | 5-48 | 3-41 | 2-97 |  |
| TG(54:10)_[M+NH<sub>4</sub>]<sup>+</sup> | 888-7076 | 0-80 | 2-80 | 2-18 | 5-48 | 3-41 | 2-97 |  |
| TG(54:11)_[M+NH<sub>4</sub>]<sup>+</sup> | 886-6924 | 1-42 | 1-65 | 0-92 | 11-89 |  |  | 19-69 |
| TG(54:11)_[M+NH<sub>4</sub>]<sup>+</sup> | 886-6919 | 1-42 | 1-65 | 0-92 | 11-89 |  |  | 19-69 |
| TG(54:12)_[M+NH<sub>4</sub>]<sup>+</sup> | 884-6763 | 0-12 |  | 0-06 | 1-84 | 1-36 | 0-64 | 0-72 |
| TG(55:00)_[M+NH<sub>4</sub>]<sup>+</sup> | 922-8797 | 3-55 | 1-75 | 2-75 | 3-51 | 3-13 | 3-96 | 4-93 |
| TG(55:01)_[M+NH<sub>4</sub>]<sup>+</sup> | 920-8641 | 1-89 | 0-96 | 0-69 | 1-67 | 2-09 | 1-26 | 2-94 |
| TG(55:02)_[M+NH<sub>4</sub>]<sup>+</sup> | 918-8484 | 0-70 | 0-72 | 0-44 | 0-26 | 0-87 | 0-74 |  |
| TG(55:03)_[M+NH<sub>4</sub>]<sup>+</sup> | 916-8328 | 0-62 | 0-79 | 1-02 | 1-45 | 1-46 | 1-35 | 1-79 |
| TG(55:05)_[M+Na]<sup>+</sup> | 917-7569 | 0-84 | 0-73 | 0-87 | 1-34 | 0-94 | 0-83 | 1-75 |
| TG(55:06)_[M+Na]<sup>+</sup> | 915-7412 | 0-63 |  | 1-20 | 2-12 | 3-12 |  | 2-17 |
| TG(55:07)_[M+Na]<sup>+</sup> | 913-7256 | 0-47 | 0-30 | 0-28 | 1-13 | 0-38 | 0-41 |  |
| TG(55:08)_[M+Na]<sup>+</sup> | 911-7099 | 1-04 |  | 1-02 |  |  |  |  |
| TG(55:09)_[M+NH<sub>4</sub>]<sup>+</sup> | 904-7389 | 0-56 | 2-01 | 0-38 | 1-07 | 1-10 | 1-19 | 1-00 |
| TG(55:10)_[M+NH<sub>4</sub>]<sup>+</sup> | 902-7232 | 0-43 | 0-24 | 0-35 | 1-61 | 0-10 | 0-26 | 0-13 |
| TG(56:00)_[M+NH<sub>4</sub>]<sup>+</sup> | 936-8954 | 5-65 | 8-98 | 10-30 | 11-62 | 12-00 | 12-42 | 14-35 |
| TG(56:03)_[M+NH<sub>4</sub>]<sup>+</sup> | 930-8484 | 7-37 | 8-36 | 8-16 | 12-44 | 17-41 | 11-42 | 13-75 |
| TG(56:04)_[M+NH<sub>4</sub>]<sup>+</sup> | 928-8328 | 6-06 | 7-49 | 9-74 | 11-42 | 10-66 | 9-50 | 11-03 |
| TG(56:06)_[M+Na]<sup>+</sup> | 929-7569 | 6-61 | 9-30 | 10-54 | 14-60 | 16-33 | 14-75 |  |
| TG(56:07)_[M+Na]<sup>+</sup> | 927-7412 | 5-82 | 8-65 |  | 12-10 | 9-75 | 8-69 |  |
| TG(56:08)_[M+Na]<sup>+</sup> | 925-7256 | 4-83 | 5-04 | 6-15 | 6-04 |  |  |  |
| TG(56:10)_[M+Na]<sup>+</sup> | 921-6943 | 2-69 | 0-78 |  | 3-16 |  | 1-21 | 3-17 |
| TG(56:10)_[M+NH<sub>4</sub>]<sup>+</sup> | 916-7389 | 0-78 | 0-87 |  | 1-14 | 2-41 |  | 1-91 |
| TG(56:11)_[M+NH<sub>4</sub>]<sup>+</sup> | 914-7232 | 0-51 | 0-48 | 0-56 | 0-93 | 1-37 |  | 1-73 |
| TG(56:12)_[M+NH<sub>4</sub>]<sup>+</sup> | 912-7076 | 0-60 |  | 0-50 |  | 0-69 |  |  |
| TG(57:00)_[M+Na]<sup>+</sup> | 955-8664 | 2-30 | 2-80 | 3-96 | 2-89 | 2-90 | 2-44 | 2-29 |
| TG(57:01)_[M+NH<sub>4</sub>]<sup>+</sup> | 948-8954 | 0-73 | 1-19 | 0-97 | 1-54 | 2-00 | 1-07 | 1-64 |
| TG(57:02)_[M+NH<sub>4</sub>]<sup>+</sup> | 946-8797 | 1-38 | 1-22 | 0-92 | 1-43 | 2-03 | 1-08 | 1-14 |
| TG(57:03)_[M+NH<sub>4</sub>]<sup>+</sup> | 944-8641 | 1-46 | 0-92 | 1-31 |  | 3-60 | 2-25 | 2-37 |
| TG(57:05)_[M+Na]<sup>+</sup> | 945-7882 | 2-49 | 1-89 | 1-43 | 1-53 | 2-14 | 1-53 | 1-90 |
| TG(57:06)_[M+Na]<sup>+</sup> | 943-7725 | 1-43 | 3-09 | 2-75 | 4-45 | 6-31 | 5-22 | 6-68 |
| TG(57:07)_[M+Na]<sup>+</sup> | 941-7569 |  | 12-07 | 12-53 |  |  | 15-58 |  |
| TG(57:08)_[M+Na]<sup>+</sup> | 939-7412 | 7-38 | 10-31 |  | 10-45 | 13-85 |  | 8-01 |
| TG(57:08)_[M+NH<sub>4</sub>]<sup>+</sup> | 934-7858 | 5-11 | 7-91 | 8-61 | 9-70 | 11-49 | 10-23 | 11-37 |
| TG(57:09)_[M+NH<sub>4</sub>]<sup>+</sup> | 932-7702 | 11-66 | 15-06 | 21-08 | 25-27 | 24-83 | 23-13 | 21-73 |
| TG(57:10)_[M+Na]<sup>+</sup> | 935-7099 | 4-63 | 5-97 | 6-46 | 6-61 |  | 8-01 |  |
| TG(57:10)_[M+NH<sub>4</sub>]<sup>+</sup> | 930-7545 | 7-30 | 8-15 | 8-05 | 12-30 | 16-64 | 10-73 |  |
| TG(57:11)_[M+NH<sub>4</sub>]<sup>+</sup> | 928-7389 | 6-27 | 7-28 |  | 12-63 |  |  |  |
| TG(57:12)_[M+NH<sub>4</sub>]<sup>+</sup> | 926-7232 | 10-02 | 9-70 | 13-83 |  |  |  |  |
| TG(58:00)_[M+NH<sub>4</sub>]<sup>+</sup> | 964-9267 | 4-56 | 7-69 | 13-32 | 5-32 | 2-04 | 7-38 | 0-55 |
| TG(58:01)_[M+Na]<sup>+</sup> | 967-8664 | 1-22 | 0-62 | 0-49 | 0-38 | 0-34 | 0-48 | 0-36 |
| TG(58:01)_[M+NH<sub>4</sub>]<sup>+</sup> | 962-9110 | 2-99 | 2-56 | 5-43 | 1-79 | 2-01 | 3-43 | 0-58 |
| TG(58:03)_[M+NH<sub>4</sub>]<sup>+</sup> | 958-8797 | 2-44 | 1-98 | 3-75 | 2-13 | 1-84 | 1-67 | 1-11 |
| TG(58:05)_[M+Na]<sup>+</sup> | 959-8038 | 3-45 | 3-44 | 6-97 | 3-26 | 2-97 | 2-74 | 1-76 |
| TG(58:06)_[M+NH<sub>4</sub>]<sup>+</sup> | 952-8328 | 2-48 | 3-52 | 3-72 | 4-12 | 4-27 | 3-30 | 2-93 |
| TG(58:07)_[M+Na]<sup>+</sup> | 955-7725 | 2-26 | 2-97 | 4-19 | 3-29 | 3-15 | 2-22 |  |
| TG(58:08)_[M+Na]<sup>+</sup> | 953-7569 | 2-86 | 4-18 |  | 4-89 | 4-96 |  |  |
| TG(58:09)_[M+Na]<sup>+</sup> | 951-7412 | 2-00 | 2-82 | 3-06 | 3-42 | 3-96 |  |  |
| TG(58:09)_[M+NH<sub>4</sub>]<sup>+</sup> | 946-7858 | 1-52 | 0-99 | 0-99 | 1-51 | 2-06 | 1-14 | 1-72 |
| TG(58:10)_[M+Na]<sup>+</sup> | 949-7256 | 1-14 | 1-63 | 1-39 | 1-92 | 2-22 |  | 1-72 |
| TG(58:10)_[M+NH<sub>4</sub>]<sup>+</sup> | 944-7702 | 1-19 | 2-05 | 1-35 | 2-48 | 3-89 | 2-46 | 2-76 |
| TG(58:11)_[M+NH<sub>4</sub>]<sup>+</sup> | 942-7545 | 4-23 |  | 7-09 | 7-84 | 10-18 |  |  |
| TG(58:14)_[M+Na]<sup>+</sup> | 941-6630 | 8-50 | 15-38 | 15-28 | 15-76 | 19-48 | 19-71 | 17-06 |
| TG(58:14)_[M+NH<sub>4</sub>]<sup>+</sup> | 936-7076 | 4-75 | 8-49 |  | 9-28 | 11-83 | 11-12 | 14-04 |
| TG(59:00)_[M+Na]<sup>+</sup> | 983-8977 | 0-16 | 0-18 | 0-25 | 0-25 | 0-26 |  | 0-29 |
| TG(59:00)_[M+NH<sub>4</sub>]<sup>+</sup> | 978-9423 | 0-77 | 0-55 | 0-57 | 0-58 | 0-42 | 0-75 | 0-38 |
| TG(59:01)_[M+Na]<sup>+</sup> | 981-8821 | 0-07 | 0-03 | 0-11 | 0-03 | 0-03 | 0-01 | 0-13 |
| TG(59:02)_[M+NH<sub>4</sub>]<sup>+</sup> | 974-9110 | 0-20 | 0-30 | 0-25 | 0-40 | 0-31 | 0-35 | 0-24 |
| TG(59:03)_[M+NH<sub>4</sub>]<sup>+</sup> | 972-8954 | 0-33 | 0-55 | 0-30 | 0-27 | 0-22 | 0-40 | 0-34 |
| TG(59:04)_[M+NH<sub>4</sub>]<sup>+</sup> | 970-8797 | 0-42 | 0-97 | 0-32 | 0-29 | 0-26 | 0-32 | 0-36 |
| TG(59:05)_[M+NH<sub>4</sub>]<sup>+</sup> | 968-8641 | 0-76 | 0-64 | 0-32 | 0-40 | 0-43 | 0-42 | 0-43 |
| TG(59:06)_[M+Na]<sup>+</sup> | 971-8038 | 0-37 | 0-70 | 0-28 | 0-27 | 0-19 | 0-28 | 0-30 |
| TG(59:06)_[M+NH<sub>4</sub>]<sup>+</sup> | 966-8489 | 1-05 | 1-07 | 1-98 | 0-98 | 0-43 | 1-51 | 0-31 |
| TG(59:07)_[M+Na]<sup>+</sup> | 969-7882 | 0-46 | 1-26 | 0-35 | 0-41 | 0-29 | 0-51 | 0-35 |
| TG(59:08)_[M+Na]<sup>+</sup> | 967-7725 | 1-92 | 0-63 |  |  |  |  | 0-34 |
| TG(59:09)_[M+Na]<sup>+</sup> | 965-7569 | 3-10 | 4-96 | 8-89 |  |  | 2-73 |  |
| TG(59:09)_[M+NH<sub>4</sub>]<sup>+</sup> | 960-8015 | 2-49 | 2-46 | 4-96 | 2-31 | 2-04 | 1-99 | 1-22 |
| TG(59:10)_[M+Na]<sup>+</sup> | 963-7412 | 13-62 | 7-96 | 36-25 | 2-18 |  | 40-88 | 0-96 |
| TG(59:10)_[M+NH<sub>4</sub>]<sup>+</sup> | 958-7863 | 2-30 | 2-03 | 3-28 | 1-52 | 1-68 | 1-06 | 1-02 |
| TG(59:11)_[M+Na]<sup>+</sup> | 961-7256 | 2-04 | 2-16 | 5-62 |  |  | 2-49 | 0-72 |
| TG(59:11)_[M+NH<sub>4</sub>]<sup>+</sup> | 956-7702 | 1-38 | 1-63 | 2-73 | 1-45 |  |  |  |
| TG(60:00)_[M+NH<sub>4</sub>]<sup>+</sup> | 992-9580 | 0-36 | 0-24 | 0-20 | 0-21 | 0-21 | 0-18 | 0-21 |
| TG(60:01)_[M+NH<sub>4</sub>]<sup>+</sup> | 990-9423 | 0-20 | 0-17 | 0-21 | 0-13 | 0-18 | 0-14 | 0-23 |
| TG(60:02)_[M+Na]<sup>+</sup> | 993-8821 | 0-30 | 0-32 | 0-09 | 0-08 | 0-17 | 0-06 | 0-13 |
| TG(60:02)_[M+NH<sub>4</sub>]<sup>+</sup> | 988-9267 | 0-22 | 0-23 | 0-21 | 0-22 | 0-38 | 0-24 | 0-25 |
| TG(60:03)_[M+NH<sub>4</sub>]<sup>+</sup> | 986-9110 | 0-20 | 0-31 | 0-45 | 0-37 | 0-46 | 0-33 | 0-26 |
| TG(60:04)_[M+NH<sub>4</sub>]<sup>+</sup> | 984-8954 | 0-19 | 0-29 | 0-49 | 0-34 | 0-26 | 0-49 | 0-26 |
| TG(60:05)_[M+Na]<sup>+</sup> | 987-8351 | 0-15 | 0-25 | 0-31 | 0-30 | 0-40 | 0-28 | 0-26 |
| TG(60:05)_[M+NH<sub>4</sub>]<sup>+</sup> | 982-8797 | 0-11 | 0-10 | 0-14 | 0-13 | 0-14 | 0-11 | 0-20 |
| TG(60:06)_[M+Na]<sup>+</sup> | 985-8195 | 0-29 | 0-49 | 0-81 | 0-58 | 0-48 | 0-54 | 0-31 |
| TG(60:07)_[M+Na]<sup>+</sup> | 983-8038 | 0-15 | 0-15 | 0-19 | 0-17 |  | 0-13 | 0-26 |
| TG(60:08)_[M+Na]<sup>+</sup> | 981-7882 | 0-07 | 0-02 | 0-02 |  |  |  | 0-08 |
| TG(60:09)_[M+Na]<sup>+</sup> | 979-7725 |  | 0-66 |  |  | 0-26 |  | 0-47 |
| TG(60:10)_[M+Na]<sup>+</sup> | 977-7569 | 0-87 | 1-92 | 1-07 | 1-88 | 0-69 |  | 0-31 |
| TG(60:10)_[M+NH<sub>4</sub>]<sup>+</sup> | 972-8015 | 0-40 | 0-59 | 0-23 | 0-34 | 0-31 | 0-33 | 0-34 |
| TG(60:11)_[M+Na]<sup>+</sup> | 975-7412 | 0-53 |  | 0-30 | 0-40 | 0-53 | 0-63 |  |
| TG(60:11)_[M+NH<sub>4</sub>]<sup>+</sup> | 970-7863 | 0-30 | 1-57 | 0-31 | 0-37 |  | 0-32 | 0-37 |
| TG(60:11)_[M+NH<sub>4</sub>]<sup>+</sup> | 970-7858 | 0-30 | 1-57 | 0-31 | 0-37 |  | 0-32 | 0-37 |
| TG(60:12)_[M+Na]<sup>+</sup> | 973-7256 | 0-34 | 0-68 | 0-37 | 0-29 |  |  |  |
| TG(60:12)_[M+NH<sub>4</sub>]<sup>+</sup> | 968-7706 | 1-20 | 0-34 | 0-28 | 0-35 | 0-27 | 0-44 |  |
| TG(60:12)_[M+NH<sub>4</sub>]<sup>+</sup> | 968-7702 | 1-20 | 0-34 | 0-28 | 0-35 | 0-27 | 0-44 |  |
| TG(60:13)_[M+NH<sub>4</sub>]<sup>+</sup> | 966-7545 | 0-82 | 1-19 | 2-22 |  |  |  |  |
| TG(60:14)_[M+NH<sub>4</sub>]<sup>+</sup> | 964-7389 |  | 15-92 | 26-71 | 10-61 |  |  | 0-57 |
| TG(60:15)_[M+Na]<sup>+</sup> | 967-6786 | 1-27 | 0-55 | 0-47 | 0-29 | 0-23 | 0-41 | 0-25 |
| TG(60:15)_[M+NH<sub>4</sub>]<sup>+</sup> | 962-7232 | 2-31 | 2-22 | 6-67 | 0-81 |  | 2-49 | 0-54 |
| TG(61:00)_[M+Na]<sup>+</sup> | 1011-9290 | 0-22 | 0-12 | 0-22 |  |  | 0-22 |  |
| TG(61:00)_[M+NH<sub>4</sub>]<sup>+</sup> | 1006-9741 | 0-20 | 0-31 | 0-39 | 0-15 | 0-28 | 0-55 | 0-19 |
| TG(61:00)_[M+NH<sub>4</sub>]<sup>+</sup> | 1006-9736 | 0-20 | 0-31 | 0-39 | 0-15 | 0-28 | 0-55 | 0-19 |
| TG(61:01)_[M+NH<sub>4</sub>]<sup>+</sup> | 1004-9580 | 0-25 | 0-22 | 0-33 | 0-24 | 0-28 | 0-15 | 0-22 |
| TG(61:03)_[M+Na]<sup>+</sup> | 1005-8821 | 0-32 | 0-35 | 0-46 | 0-24 | 0-29 | 0-18 | 0-22 |
| TG(61:05)_[M+NH<sub>4</sub>]<sup>+</sup> | 996-8954 |  | 0-02 |  |  | 0-11 | 0-02 | 0-02 |
| TG(61:06)_[M+NH<sub>4</sub>]<sup>+</sup> | 994-8797 | 0-18 | 0-25 | 0-02 | 0-04 | 0-09 | 0-03 | 0-11 |
| TG(61:07)_[M+NH<sub>4</sub>]<sup>+</sup> | 992-8641 | 0-36 | 0-33 | 0-21 | 0-22 | 0-22 | 0-19 | 0-22 |
| TG(61:10)_[M+Na]<sup>+</sup> | 991-7725 | 0-24 | 0-22 | 0-24 |  | 0-20 |  | 0-17 |
| TG(61:10)_[M+NH<sub>4</sub>]<sup>+</sup> | 986-8171 | 0-19 | 0-32 | 0-48 | 0-37 | 0-37 | 0-34 | 0-25 |
| TG(61:11)_[M+Na]<sup>+</sup> | 989-7569 | 0-25 | 0-18 | 0-21 | 0-19 | 0-28 | 0-18 | 0-24 |
| TG(61:11)_[M+NH<sub>4</sub>]<sup>+</sup> | 984-8015 | 0-17 | 0-13 | 0-43 | 0-30 |  |  | 0-23 |
| TG(61:12)_[M+Na]<sup>+</sup> | 987-7412 | 0-17 | 0-27 | 0-39 | 0-34 | 0-42 | 0-34 | 0-27 |
| TG(62:00)_[M+Na]<sup>+</sup> | 1025-9447 | 0-52 | 0-28 | 0-50 |  |  | 0-32 |  |
| TG(62:00)_[M+NH<sub>4</sub>]<sup>+</sup> | 1020-9893 | 0-33 | 0-17 | 0-39 | 0-23 | 0-72 | 0-18 |  |
| TG(62:01)_[M+Na]<sup>+</sup> | 1023-9290 | 0-35 | 0-19 | 0-35 | 0-19 |  | 0-25 |  |
| TG(62:01)_[M+NH<sub>4</sub>]<sup>+</sup> | 1018-9736 | 0-34 | 0-14 | 0-32 | 0-17 | 0-49 | 0-14 | 0-28 |
| TG(62:02)_[M+Na]<sup>+</sup> | 1021-9134 | 0-40 | 0-17 | 0-39 | 0-20 | 0-56 | 0-19 | 0-32 |
| TG(62:02)_[M+NH<sub>4</sub>]<sup>+</sup> | 1016-9580 | 0-54 | 0-30 | 0-48 | 0-36 | 0-60 |  |  |
| TG(62:03)_[M+Na]<sup>+</sup> | 1019-8977 | 0-36 | 0-13 | 0-32 | 0-14 | 0-48 | 0-14 | 0-28 |
| TG(62:03)_[M+NH<sub>4</sub>]<sup>+</sup> | 1014-9423 | 0-36 | 0-14 | 0-35 | 0-25 |  | 0-20 |  |
| TG(62:04)_[M+NH<sub>4</sub>]<sup>+</sup> | 1012-9267 | 0-24 | 0-14 | 0-26 | 0-30 | 0-42 | 0-30 | 0-37 |
| TG(62:05)_[M+NH<sub>4</sub>]<sup>+</sup> | 1010-9110 | 0-11 |  | 0-13 | 0-20 | 0-39 | 0-18 | 0-21 |
| TG(62:06)_[M+Na]<sup>+</sup> | 1013-8508 | 0-25 | 0-14 | 0-23 | 0-24 | 0-41 | 0-21 | 0-29 |
| TG(62:06)_[M+NH<sub>4</sub>]<sup>+</sup> | 1008-8954 |  | 0-11 | 0-06 | 0-03 | 0-27 | 0-16 | 0-12 |
| TG(62:07)_[M+Na]<sup>+</sup> | 1011-8351 | 0-25 | 0-17 | 0-26 | 0-33 | 0-43 | 0-32 | 0-31 |
| TG(62:07)_[M+NH<sub>4</sub>]<sup>+</sup> | 1006-8802 | 0-19 | 0-22 | 0-41 | 0-59 | 0-45 | 0-39 | 0-19 |
| TG(62:08)_[M+Na]<sup>+</sup> | 1009-8195 | 0-26 |  | 0-33 | 0-40 | 0-53 | 0-43 | 0-29 |
| TG(62:08)_[M+NH<sub>4</sub>]<sup>+</sup> | 1004-8641 | 0-27 | 0-31 | 0-33 | 0-24 | 0-24 | 0-21 | 0-24 |
| TG(62:09)_[M+Na]<sup>+</sup> | 1007-8038 | 0-18 | 0-31 |  |  | 0-17 | 0-27 |  |
| TG(62:11)_[M+Na]<sup>+</sup> | 1003-7725 | 0-21 | 0-30 | 0-12 | 0-07 | 0-20 |  | 0-39 |
| TG(62:15)_[M+Na]<sup>+</sup> | 995-7099 | 0-09 | 0-09 | 0-01 | 0-05 | 0-12 | 0-04 | 0-12 |
| TG(62:15)_[M+NH<sub>4</sub>]<sup>+</sup> | 990-7545 | 0-15 | 0-09 | 0-15 | 0-11 |  | 0-10 | 0-21 |
| TG(62:16)_[M+NH<sub>4</sub>]<sup>+</sup> | 988-7389 | 0-24 | 0-25 | 0-24 | 0-23 | 0-31 |  | 0-26 |
| TG(63:00)_[M+Na]<sup>+</sup> | 1039-9603 | 1-22 | 1-00 | 1-24 |  |  | 0-84 |  |
| TG(63:00)_[M+NH<sub>4</sub>]<sup>+</sup> | 1035-0049 | 1-04 | 0-52 | 1-20 | 1-02 | 1-40 |  |  |
| TG(63:01)_[M+Na]<sup>+</sup> | 1037-9447 | 1-17 | 0-80 | 0-99 |  |  |  |  |
| TG(63:01)_[M+NH<sub>4</sub>]<sup>+</sup> | 1032-9893 | 0-85 | 0-32 | 1-05 | 0-93 | 1-12 |  |  |
| TG(63:02)_[M+Na]<sup>+</sup> | 1035-9290 | 1-18 | 0-81 | 1-12 | 1-14 | 1-59 | 0-51 |  |
| TG(63:02)_[M+NH<sub>4</sub>]<sup>+</sup> | 1030-9736 | 0-65 | 0-27 | 0-95 | 0-62 | 0-81 |  | 0-34 |
| TG(63:03)_[M+Na]<sup>+</sup> | 1033-9134 | 0-96 | 0-63 | 0-94 | 0-94 | 1-35 | 0-39 | 0-50 |
| TG(63:03)_[M+NH<sub>4</sub>]<sup>+</sup> | 1028-9580 | 0-74 | 0-19 | 0-61 |  | 0-79 |  |  |
| TG(63:04)_[M+NH<sub>4</sub>]<sup>+</sup> | 1026-9423 | 0-69 | 0-31 | 0-74 | 0-34 |  | 0-26 |  |
| TG(63:05)_[M+Na]<sup>+</sup> | 1029-8821 | 0-72 | 0-40 | 0-71 | 0-60 | 0-94 | 0-30 | 0-40 |
| TG(63:05)_[M+NH<sub>4</sub>]<sup>+</sup> | 1024-9267 | 0-44 | 0-30 | 0-36 | 0-30 |  |  |  |
| TG(63:06)_[M+Na]<sup>+</sup> | 1027-8664 | 0-63 | 0-36 | 0-65 | 0-53 | 0-77 | 0-30 | 0-45 |
| TG(63:06)_[M+NH<sub>4</sub>]<sup>+</sup> | 1022-9110 | 0-38 |  | 0-30 | 0-03 | 0-21 |  | 0-17 |
| TG(63:07)_[M+Na]<sup>+</sup> | 1025-8508 | 0-50 | 0-40 | 0-48 | 0-26 | 0-78 | 0-25 | 0-28 |
| TG(63:07)_[M+NH<sub>4</sub>]<sup>+</sup> | 1020-8954 | 0-40 | 0-16 | 0-38 | 0-18 | 0-72 | 0-18 | 0-32 |
| TG(63:09)_[M+NH<sub>4</sub>]<sup>+</sup> | 1016-8641 | 0-51 | 0-32 | 0-42 | 0-38 | 0-62 | 0-22 | 0-33 |
| TG(63:10)_[M+NH<sub>4</sub>]<sup>+</sup> | 1014-8484 | 0-28 | 0-17 | 0-26 | 0-26 | 0-59 | 0-20 | 0-28 |
| TG(63:11)_[M+NH<sub>4</sub>]<sup>+</sup> | 1012-8328 | 0-22 | 0-17 | 0-21 | 0-33 | 0-44 | 0-28 |  |
| TG(63:12)_[M+Na]<sup>+</sup> | 1015-7725 | 0-42 | 0-21 |  | 0-40 |  |  | 0-41 |
| TG(63:12)_[M+NH<sub>4</sub>]<sup>+</sup> | 1010-8171 | 0-16 |  | 0-20 | 0-27 |  |  | 0-19 |
| TG(63:13)_[M+Na]<sup>+</sup> | 1013-7569 |  | 0-16 | 0-19 | 0-28 | 0-31 | 0-23 | 0-31 |
| TG(64:01)_[M+Na]<sup>+</sup> | 1051-9603 | 0-14 | 0-08 | 0-08 |  |  | 0-20 |  |
| TG(64:01)_[M+NH<sub>4</sub>]<sup>+</sup> | 1047-0049 | 0-16 | 0-10 | 0-14 | 0-16 | 0-20 | 0-15 |  |
| TG(64:02)_[M+NH<sub>4</sub>]<sup>+</sup> | 1044-9893 | 0-51 | 0-53 | 0-47 | 0-71 | 0-68 | 0-34 |  |
| TG(64:03)_[M+Na]<sup>+</sup> | 1047-9290 | 0-10 | 0-10 | 0-08 | 0-13 | 0-14 | 0-18 |  |
| TG(64:03)_[M+NH<sub>4</sub>]<sup>+</sup> | 1042-9736 | 1-61 | 1-05 | 1-31 | 2-62 | 1-95 |  |  |
| TG(64:04)_[M+Na]<sup>+</sup> | 1045-9134 | 0-36 | 0-28 | 0-26 | 0-38 | 0-36 | 0-27 |  |
| TG(64:04)_[M+NH<sub>4</sub>]<sup>+</sup> | 1040-9580 | 1-29 | 1-13 | 1-18 |  | 1-59 | 0-80 |  |
| TG(64:05)_[M+NH<sub>4</sub>]<sup>+</sup> | 1038-9423 | 1-30 | 0-89 | 1-06 |  | 1-70 | 0-72 |  |
| TG(64:06)_[M+Na]<sup>+</sup> | 1041-8821 | 1-19 | 1-34 | 1-26 | 2-05 | 1-76 | 0-78 |  |
| TG(64:06)_[M+NH<sub>4</sub>]<sup>+</sup> | 1036-9267 | 1-37 | 0-95 | 0-99 | 1-27 | 1-66 | 0-41 |  |
| TG(64:07)_[M+Na]<sup>+</sup> | 1039-8664 | 1-35 | 1-07 | 1-21 | 1-69 | 1-77 | 0-81 |  |
| TG(64:07)_[M+NH<sub>4</sub>]<sup>+</sup> | 1034-9110 | 1-02 | 0-66 | 0-97 | 1-00 | 1-43 | 0-42 |  |
| TG(64:08)_[M+Na]<sup>+</sup> | 1037-8508 | 1-40 | 0-99 | 1-24 | 1-33 |  | 0-41 |  |
| TG(64:09)_[M+NH<sub>4</sub>]<sup>+</sup> | 1030-8797 | 0-77 | 0-43 | 0-75 | 0-61 | 0-99 | 0-29 | 0-46 |
| TG(64:10)_[M+Na]<sup>+</sup> | 1033-8195 | 1-02 | 0-46 | 0-70 | 1-02 |  | 0-46 |  |
| TG(64:10)_[M+NH<sub>4</sub>]<sup>+</sup> | 1028-8641 | 0-61 | 0-32 | 0-59 | 0-45 | 0-77 | 0-23 | 0-33 |
| TG(64:11)_[M+NH<sub>4</sub>]<sup>+</sup> | 1026-8484 | 0-57 | 0-30 | 0-43 | 0-32 | 0-77 | 0-26 | 0-27 |
| TG(64:12)_[M+Na]<sup>+</sup> | 1029-7882 | 0-86 | 0-40 | 0-57 | 0-55 | 1-19 | 0-31 | 0-40 |
| TG(64:12)_[M+NH<sub>4</sub>]<sup>+</sup> | 1024-8328 | 0-54 | 0-40 |  | 0-33 | 0-97 |  |  |
| TG(64:13)_[M+Na]<sup>+</sup> | 1027-7725 | 0-63 | 0-35 | 0-49 |  | 1-07 | 0-30 | 0-36 |
| TG(64:13)_[M+NH<sub>4</sub>]<sup>+</sup> | 1022-8171 | 0-35 |  |  |  | 0-25 |  | 0-20 |
| TG(64:14)_[M+Na]<sup>+</sup> | 1025-7569 | 0-47 | 0-25 | 0-45 | 0-22 | 1-12 | 0-23 | 0-34 |
| TG(64:15)_[M+NH<sub>4</sub>]<sup>+</sup> | 1018-7858 | 0-34 |  | 0-22 | 0-18 |  |  | 0-19 |
| TG(64:16)_[M+Na]<sup>+</sup> | 1021-7256 | 0-41 | 0-17 | 0-40 | 0-24 | 0-58 | 0-21 | 0-36 |
| TG(65:00)_[M+Na]<sup>+</sup> | 1067-9916 | 0-35 | 0-72 | 0-75 | 0-29 | 0-48 |  |  |
| TG(65:00)_[M+NH<sub>4</sub>]<sup>+</sup> | 1063-0362 | 1-07 | 1-69 | 1-46 | 0-28 | 0-62 |  |  |
| TG(65:01)_[M+NH<sub>4</sub>]<sup>+</sup> | 1061-0206 | 0-55 | 0-45 | 0-51 |  | 0-59 | 0-22 |  |
| TG(65:02)_[M+Na]<sup>+</sup> | 1063-9603 | 0-32 | 0-90 | 0-70 | 0-21 |  |  |  |
| TG(65:02)_[M+NH<sub>4</sub>]<sup>+</sup> | 1059-0049 | 0-43 | 0-17 | 0-28 | 0-35 | 0-47 |  |  |
| TG(65:03)_[M+NH<sub>4</sub>]<sup>+</sup> | 1056-9893 | 0-18 |  | 0-16 | 0-03 |  |  |  |

|  |  |  |  |  |  |  |  |
| --- | --- | --- | --- | --- | --- | --- | --- |
| TG(65:04)_[M+Na]<sup>+</sup> | 1059-9290 | 0-59 | 0-32 | 0-43 | 0-54 | 0-59 | 0-30 |
| TG(65:04)_[M+NH<sub>4</sub>]<sup>+</sup> | 1054-9736 | 0-20 | 0-02 | 0-02 | 0-04 |  |  |
| TG(65:05)_[M+Na]<sup>+</sup> | 1057-9134 | 0-21 |  | 0-06 | 0-12 | 0-25 | 0-03 |
| TG(65:05)_[M+NH<sub>4</sub>]<sup>+</sup> | 1052-9580 |  | 0-05 | 0-14 |  | 0-15 |  |
| TG(65:06)_[M+Na]<sup>+</sup> | 1055-8977 | 0-33 | 0-07 | 0-24 | 0-18 | 0-23 | 0-10 |
| TG(65:08)_[M+Na]<sup>+</sup> | 1051-8664 | 0-10 |  | 0-31 | 0-05 | 0-14 | 0-14 |
| TG(65:08)_[M+NH<sub>4</sub>]<sup>+</sup> | 1046-9110 | 0-17 | 0-15 | 0-15 | 0-23 | 0-21 | 0-24 |
| TG(65:09)_[M+NH<sub>4</sub>]<sup>+</sup> | 1044-8954 | 0-58 | 0-60 | 0-52 | 0-78 | 0-74 | 0-36 |
| TG(65:10)_[M+NH<sub>4</sub>]<sup>+</sup> | 1042-8797 | 1-79 | 1-72 | 1-54 | 2-23 | 1-95 | 0-66 |
| TG(65:11)_[M+NH<sub>4</sub>]<sup>+</sup> | 1040-8641 | 1-28 | 1-31 | 1-22 | 1-61 | 1-63 | 0-79 |
| TG(65:12)_[M+Na]<sup>+</sup> | 1043-8038 | 1-31 | 1-14 | 1-05 | 1-41 |  | 0-52 |
| TG(65:12)_[M+NH<sub>4</sub>]<sup>+</sup> | 1038-8484 | 1-36 |  | 1-13 | 1-83 |  | 0-49 |
| TG(66:01)_[M+Na]<sup>+</sup> | 1079-9916 |  | 0-62 | 0-30 |  | 0-18 |  |
| TG(66:04)_[M+NH<sub>4</sub>]<sup>+</sup> | 1068-9893 | 0-31 | 0-55 | 0-56 | 0-42 | 0-34 |  |
| TG(66:06)_[M+Na]<sup>+</sup> | 1069-9134 | 0-04 | 0-12 | 0-10 | 0-98 | 0-31 |  |
| TG(66:06)_[M+NH<sub>4</sub>]<sup>+</sup> | 1064-9580 | 0-36 | 0-51 | 0-59 |  | 0-53 | 0-08 |
| TG(66:07)_[M+Na]<sup>+</sup> | 1067-8977 |  | 0-43 | 0-40 | 0-24 | 0-77 |  |
| TG(66:07)_[M+NH<sub>4</sub>]<sup>+</sup> | 1062-9423 | 0-37 | 1-32 | 0-84 | 0-16 | 0-38 |  |
| TG(66:08)_[M+Na]<sup>+</sup> | 1065-8821 | 0-44 | 0-36 | 0-35 | 0-48 | 0-60 |  |
| TG(66:08)_[M+NH<sub>4</sub>]<sup>+</sup> | 1060-9267 | 0-55 | 0-45 | 0-52 | 0-54 | 0-55 | 0-23 |
| TG(66:11)_[M+Na]<sup>+</sup> | 1059-8351 | 0-59 | 0-31 | 0-40 |  |  |  |
| TG(66:11)_[M+NH<sub>4</sub>]<sup>+</sup> | 1054-8797 | 0-14 | 0-07 | 0-06 | 0-10 | 0-10 |  |
| TG(66:12)_[M+Na]<sup>+</sup> | 1057-8195 | 0-19 |  | 0-12 | 0-29 | 0-28 |  |
| TG(66:12)_[M+NH<sub>4</sub>]<sup>+</sup> | 1052-8641 |  | 0-17 | 0-20 | 0-08 | 0-17 | 0-14 |
| TG(66:15)_[M+Na]<sup>+</sup> | 1051-7725 |  | 0-13 |  | 0-05 |  | 0-22 |
| TG(66:16)_[M+Na]<sup>+</sup> | 1049-7569 | 0-04 | 0-04 |  |  | 0-06 | 0-22 |
| TG(66:17)_[M+Na]<sup>+</sup> | 1047-7412 | 0-09 | 0-09 | 0-07 | 0-13 | 0-13 | 0-18 |
| TG(66:17)_[M+NH<sub>4</sub>]<sup>+</sup> | 1042-7858 | 2-14 | 1-99 | 1-79 | 2-46 | 2-14 | 0-76 |
| TG(66:18)_[M+Na]<sup>+</sup> | 1045-7256 | 0-43 | 0-36 | 0-30 | 0-46 | 0-40 | 0-29 |
| TG(66:18)_[M+NH<sub>4</sub>]<sup>+</sup> | 1040-7702 | 1-36 | 0-83 | 1-06 | 1-72 |  | 0-78 |
| TG(67:01)_[M+NH<sub>4</sub>]<sup>+</sup> | 1089-0523 | 0-40 | 0-20 | 0-18 | 0-40 | 0-34 |  |
| TG(67:03)_[M+NH<sub>4</sub>]<sup>+</sup> | 1085-0210 | 0-35 | 0-15 | 0-13 | 0-34 | 0-27 |  |
| TG(67:11)_[M+NH<sub>4</sub>]<sup>+</sup> | 1068-8958 | 0-24 | 0-40 | 0-74 | 0-40 | 0-63 |  |
| TG(68:12)_[M+NH<sub>4</sub>]<sup>+</sup> | 1080-8958 | 0-26 | 0-58 | 0-18 | 0-20 | 0-31 |  |

**Table S6.** Relative signal intensities for high resolution m/z values consistent with sterol isoforms (n = 2). Signals were assigned according to high resolution masses with a deviation of <12 ppm from the calculated monoisotopic mass. Quantitative standards were not used. G, guanidinium. Collection points: +2 h, early part of G_1_; +4 h, mid-G_1_ (first CP); +6 h, end of G_1_ and pS; +8 h, S and second CP; +10 h, G_2_; +13 h, M; +15 h, G_3_, for others see Figure 1.

| Sterol annotation | m/z | Collection point |  |  |  |  |  |  |
| --- | --- | --- | --- | --- | --- | --- | --- | --- |
|  |  | +2 h | +4h | +6 h | +8 h | +10 h | +13 h | +15 h |
| 16:0 Campesteryl ester [M+H]<sup>+</sup> | 639-608 | 0-02 | 0-14 | 0-06 | 0-71 | 0-42 | 0-45 | 1-31 |
| 16:0 Campesteryl ester [M+K]<sup>+</sup> | 677-5634 | 1-59 | 2-20 | 2-31 | 1-40 | 1-17 | 1-22 | 1-25 |
| 16:0 Campesteryl ester [M+NH<sub>4</sub>]<sup>+</sup> | 656-634 | 0-32 |  | 0-69 | 0-80 | 1-01 |  | 1-38 |
| 16:0 Sitosteryl ester [M+H]<sup>+</sup> | 653-6236 | 0-18 | 0-13 |  | 0-07 |  |  |  |
| 16:0 Sitosteryl ester [M+Na]<sup>+</sup> | 675-6056 | 1-08 | 0-85 | 0-32 | 0-24 | 0-36 | 0-42 | 1-23 |
| 16:0 Stigmasteryl ester/16:1 Sitosteryl ester/16:0 Brassicasteryl ester [M+G]<sup>+</sup> | 745-6252 |  | 0-31 |  | 0-98 | 1-67 | 2-02 | 3-70 |
| 16:0 Stigmasteryl ester/16:1 Sitosteryl ester/16:0 Brassicasteryl ester [M+Na]<sup>+</sup> | 673-59 | 0-22 | 0-49 | 0-87 | 0-65 | 0-22 | 0-35 | 0-67 |
| 16:0 Stigmasteryl ester/16:1 Sitosteryl ester/16:0 Brassicasteryl ester [M+NH<sub>4</sub>]<sup>+</sup> | 668-634 | 0-25 | 0-30 |  | 0-11 | 0-11 | 0-17 | 0-28 |
| 16:1 Campesteryl ester [M+G]<sup>+</sup> | 731-6095 | 4-89 | 2-37 |  | 2-57 | 9-34 | 10-19 | 27-21 |
| 16:1 Campesteryl ester [M+H]<sup>+</sup> | 637-5923 | 0-11 | 0-27 |  | 0-39 | 0-12 | 0-21 | 0-38 |
| 16:1 Campesteryl ester [M+K]<sup>+</sup> | 675-5477 | 0-84 | 0-61 | 0-08 | 0-05 | 0-12 | 0-21 | 1-06 |
| 16:1 Stigmasteryl ester/16:2 Sitosteryl ester [M+G]<sup>+</sup> | 743-6095 | 1-35 | 1-98 | 1-70 | 4-55 | 4-58 | 3-28 | 4-33 |
| 16:1 Stigmasteryl ester/16:2 Sitosteryl ester [M+Na]<sup>+</sup> | 671-5743 | 0-07 | 0-25 | 0-16 | 0-13 | 0-20 | 0-12 |  |
| 16:2 Campesteryl ester [M+G]<sup>+</sup> | 729-5939 | 16-78 | 11-96 | 9-42 | 10-98 | 16-91 | 18-71 | 25-39 |
| 16:2 Campesteryl ester [M+H]<sup>+</sup> | 635-5767 | 0-13 | 0-53 | 0-34 | 0-52 | 0-19 | 0-08 | 0-72 |
| 16:2 Stigmasteryl ester/16:3 Sitosteryl ester [M+G]<sup>+</sup> | 741-5939 | 3-33 | 3-96 | 5-17 | 8-88 | 6-99 | 4-40 | 3-70 |
| 16:2 Stigmasteryl ester/16:3 Sitosteryl ester [M+H]<sup>+</sup> | 647-5767 | 0-58 | 0-17 | 0-16 | 0-24 | 0-18 | 0-15 | 0-18 |
| 16:3 Campesteryl ester/16:0 ergosteryl ester [M+G]<sup>+</sup> | 727-5782 | 41-56 | 34-30 | 23-66 | 16-99 | 28-99 | 34-08 | 35-77 |
| 16:3 Stigmasteryl ester [M+G]<sup>+</sup> | 739-5782 | 4-39 | 5-52 | 5-41 | 12-92 | 13-00 | 9-93 | 11-89 |
| 16:3 Stigmasteryl ester [M+K]<sup>+</sup> | 683-5164 | 0-93 | 1-18 | 2-05 | 0-51 | 0-37 | 1-03 | 0-26 |
| 18:0 Campesteryl ester [M+H]<sup>+</sup> | 667-6393 | 0-42 | 0-36 | 0-45 | 0-37 | 0-22 | 0-35 | 0-30 |
| 18:0 Campesteryl ester [M+K]<sup>+</sup> | 705-5947 | 37-00 | 43-13 |  |  |  | 32-73 |  |
| 18:0 Campesteryl ester [M+Na]<sup>+</sup> | 689-6213 | 0-16 | 0-32 |  | 0-10 |  | 0-03 | 0-23 |
| 18:0 Campesteryl ester [M+NH<sub>4</sub>]<sup>+</sup> | 684-6653 | 2-00 |  | 3-97 | 1-84 |  | 3-33 | 0-59 |
| 18:0 Sitosteryl ester [M+H]<sup>+</sup> | 681-6549 | 0-42 | 0-05 | 0-66 | 0-48 |  | 0-30 | 0-21 |
| 18:0 Sitosteryl ester [M+K]<sup>+</sup> | 719-6103 | 3-83 | 2-77 | 3-11 |  |  |  | 4-98 |
| 18:0 Sitosteryl ester [M+Na]<sup>+</sup> | 703-6369 | 5-16 | 6-60 | 5-51 | 7-23 | 7-12 | 8-32 | 5-99 |
| 18:1 Campesteryl ester/18:0 Brassicasteryl ester [M+G]<sup>+</sup> | 759-6408 | 17-27 | 20-01 | 15-91 | 44-84 | 49-51 | 44-52 | 51-49 |
| 18:1 Campesteryl ester/18:0 Brassicasteryl ester [M+K]<sup>+</sup> | 703-579 | 4-13 | 5-88 | 3-34 | 7-46 |  |  |  |
| 18:1 Campesteryl ester/18:0 Brassicasteryl ester [M+Na]<sup>+</sup> | 687-6056 | 0-16 | 0-45 | 0-57 | 0-27 | 0-22 | 0-31 | 0-31 |
| 18:1 Campesteryl ester/18:0 Brassicasteryl ester [M+NH<sub>4</sub>]<sup>+</sup> | 682-6496 | 0-61 |  | 1-12 | 0-39 |  |  |  |
| 18:1 Sitosteryl ester/18:0 Stigmasteryl ester [M+K]<sup>+</sup> | 717-5947 | 2-25 | 3-91 | 1-59 | 3-29 | 2-93 | 1-13 | 1-46 |
| 18:1 Stigmasteryl ester/18:2 Sitosteryl ester [M+G]<sup>+</sup> | 771-6408 | 12-72 | 13-42 | 14-51 | 22-12 | 16-62 | 14-44 | 6-92 |
| 18:1 Stigmasteryl ester/18:2 Sitosteryl ester [M+H]<sup>+</sup> | 677-6236 | 1-63 | 1-99 | 1-69 |  | 1-21 | 1-51 | 1-15 |
| 18:1 Stigmasteryl ester/18:2 Sitosteryl ester [M+Na]<sup>+</sup> | 699-6056 | 0-65 | 1-37 | 1-38 | 1-21 | 0-85 | 0-82 |  |
| 18:2 Stigmasteryl ester/18:3 Sitosteryl ester [M+G]<sup>+</sup> | 769-6252 | 30-59 | 28-31 | 33-72 | 35-78 | 25-78 | 23-07 | 8-36 |
| 18:2 Stigmasteryl ester/18:3 Sitosteryl ester [M+H]<sup>+</sup> | 675-608 | 1-26 | 1-04 | 0-57 | 0-50 | 0-57 | 0-63 | 1-36 |
| 18:3 Campesteryl ester/18:1 ergosteryl ester/18:2 Brassicasteryl ester [M+G]<sup>+</sup> | 755-6095 | 68-06 | 48-43 | 42-25 | 38-77 | 39-35 | 40-54 | 53-81 |
| 18:3 Campesteryl ester/18:1 ergosteryl ester/18:2 Brassicasteryl ester [M+Na]<sup>+</sup> | 683-5743 | 1-24 | 1-46 | 2-42 | 0-86 | 0-57 | 1-29 | 0-40 |
| 18:3 Campesteryl ester/18:1 ergosteryl ester/18:2 Brassicasteryl ester [M+NH<sub>4</sub>]<sup>+</sup> | 678-6183 | 1-45 | 1-22 | 1-45 | 1-82 |  | 1-59 |  |
| 18:3 Stigmasteryl ester [M+G]<sup>+</sup> | 767-6095 | 47-99 | 49-95 | 69-38 | 69-05 | 72-65 | 47-57 | 20-67 |
| 18:3 Stigmasteryl ester [M+H]<sup>+</sup> | 673-5923 | 0-13 | 0-38 | 0-73 | 0-50 | 0-11 | 0-25 | 0-60 |
| 18:3 Stigmasteryl ester [M+K]<sup>+</sup> | 711-5477 | 2-36 | 2-39 | 2-94 | 3-97 | 4-13 | 4-18 | 6-34 |
| 20:0 Campesteryl ester [M+Na]<sup>+</sup> | 717-6526 | 2-11 | 3-72 | 1-26 | 1-29 | 2-71 | 2-72 | 1-33 |
| 20:0 Sitosteryl ester [M+H]<sup>+</sup> | 709-6862 | 2-27 | 1-50 | 1-59 |  | 2-74 |  | 4-48 |
| 20:0 Sitosteryl ester [M+Na]<sup>+</sup> | 731-6682 | 8-63 | 7-18 | 4-41 | 6-50 | 15-15 | 14-43 | 26-96 |
| 20:0 Stigmasteryl ester/20:1 Sitosteryl ester [M+H]<sup>+</sup> | 707-6706 |  | 15-85 | 11-38 | 14-68 | 13-79 |  |  |
| 20:0 Stigmasteryl ester/20:1 Sitosteryl ester [M+K]<sup>+</sup> | 745-626 | 0-53 | 1-04 | 0-27 | 2-35 | 2-78 | 2-96 | 4-39 |
| 20:0 Stigmasteryl ester/20:1 Sitosteryl ester [M+NH<sub>4</sub>]<sup>+</sup> | 724-6966 | 18-16 |  | 8-67 |  |  | 9-02 |  |
| 20:1 Campesteryl ester [M+G]<sup>+</sup> | 787-6721 |  |  |  | 0-17 | 5-24 | 6-50 | 14-26 |
| 20:1 Campesteryl ester [M+K]<sup>+</sup> | 731-6103 | 8-92 | 6-65 | 3-95 | 8-11 | 13-81 | 13-97 | 27-13 |
| 20:1 Stigmasteryl ester/20:2 Sitosteryl ester [M+H]<sup>+</sup> | 705-6549 |  | 31-68 | 36-35 | 34-94 | 24-83 | 30-13 | 35-67 |
| 20:1 Stigmasteryl ester/20:2 Sitosteryl ester [M+K]<sup>+</sup> | 743-6103 | 1-41 | 2-02 | 1-78 | 4-58 | 4-64 | 3-33 | 4-89 |
| 20:1 Stigmasteryl ester/20:2 Sitosteryl ester [M+Na]<sup>+</sup> | 727-6369 | 39-92 | 33-40 | 23-08 | 25-97 | 28-27 | 27-90 | 34-33 |
| 20:1 Stigmasteryl ester/20:2 Sitosteryl ester [M+NH<sub>4</sub>]<sup>+</sup> | 722-6809 | 4-82 | 4-78 | 3-05 | 3-59 | 3-39 | 3-16 | 4-26 |
| 20:2 Campesteryl ester [M+H]<sup>+</sup> | 691-6393 | 1-34 | 2-01 | 1-92 | 1-40 |  | 0-77 |  |
| 20:2 Campesteryl ester [M+K]<sup>+</sup> | 729-5947 | 16-68 | 11-85 | 9-27 | 10-83 | 16-79 | 18-61 | 25-32 |
| 20:2 Campesteryl ester $[M+Na]^+$ | 713-6213 | 0-50 | 0-71 | | | 1-29 | 0-77 | |
| 20:2 Stigmasteryl ester/20:3 Sitosteryl ester $[M+G]^+$ | 797-6565 | 6-03 | 5-97 | 8-56 | 7-97 | 4-08 | 2-41 | 0-97 |
| 20:2 Stigmasteryl ester/20:3 Sitosteryl ester $[M+H]^+$ | 703-6393 | 1-20 | 2-07 | 1-11 | 1-37 | 2-39 | 4-30 | 3-06 |
| 20:2 Stigmasteryl ester/20:3 Sitosteryl ester $[M+K]^+$ | 741-5947 | 3-81 | 4-47 | 5-81 | 9-54 | 7-53 | 4-85 | 4-03 |
| 20:2 Stigmasteryl ester/20:3 Sitosteryl ester $[M+Na]^+$ | 725-6213 | 17-48 | 11-90 | 9-94 | 8-98 | 11-47 | | |
| 20:3 Campesteryl ester $[M+G]^+$ | 783-6408 | 19-92 | 32-45 | 36-39 | 69-63 | 71-49 | 58-70 | 50-51 |
| 20:3 Campesteryl ester $[M+K]^+$ | 727-579 | 32-65 | 24-84 | 11-72 | 4-55 | 19-98 | 25-68 | 29-65 |
| 20:3 Campesteryl ester $[M+NH_4]^+$ | 706-6496 | 19-01 | 19-81 | 19-40 | 31-84 | 20-37 | 20-31 | 29-90 |
| 20:3 Stigmasteryl ester $[M+H]^+$ | 701-6236 | 1-56 | 1-20 | 1-38 | 1-10 | 2-56 | 2-57 | 1-20 |
| 20:3 Stigmasteryl ester $[M+K]^+$ | 739-579 | 6-15 | 7-40 | 7-77 | 15-34 | 14-95 | | 13-74 |
| 20:3 Stigmasteryl ester $[M+Na]^+$ | 723-6056 | | 5-80 | 3-55 | 2-04 | 2-61 | 3-18 | 7-11 |
| 22:0 Sitosteryl ester $[M+NH_4]^+$ | 754-7435 | 62-85 | 54-09 | 50-74 | 59-35 | 43-05 | 44-17 | 43-23 |
| 22:0 Stigmasteryl ester $[M+G]^+$ | 829-7191 | 1-62 | 1-39 | 0-44 | 0-91 | 0-70 | 0-50 | 0-66 |
| 22:0 Stigmasteryl ester $[M+H]^+$ | 735-7019 | 65-11 | 84-93 | 63-88 | | 100-13 | 69-86 | |
| 22:0 Stigmasteryl ester $[M+NH_4]^+$ | 752-7279 | 22-42 | 19-46 | 9-61 | 11-38 | 9-05 | 13-10 | 21-73 |
| 22:1 Campesteryl ester $[M+K]^+$ | 759-6416 | 17-50 | 20-25 | 16-21 | 45-15 | 49-76 | 44-73 | 51-64 |
| 22:1 Campesteryl ester $[M+NH_4]^+$ | 738-7122 | 9-51 | 13-72 | 13-69 | 26-03 | 34-28 | | 31-21 |
| 22:1 Stigmasteryl ester/22:2 Sitosteryl ester $[M+K]^+$ | 771-6416 | 12-93 | 13-64 | 14-79 | 22-41 | 16-85 | 14-63 | 7-06 |
| 22:2 Campesteryl ester $[M+G]^+$ | 813-6878 | 0-20 | 2-29 | | 0-69 | 0-39 | 0-67 | 0-30 |
| 22:2 Campesteryl ester $[M+H]^+$ | 719-6706 | 3-23 | 3-68 | 2-29 | 2-58 | 3-25 | 2-21 | 4-00 |
| 22:2 Campesteryl ester $[M+K]^+$ | 757-626 | 49-87 | 47-98 | 46-51 | 58-76 | 56-76 | 49-07 | 54-41 |
| 22:2 Stigmasteryl ester/22:3 Sitosteryl ester $[M+H]^+$ | 731-6706 | 8-51 | 7-05 | 4-24 | 6-32 | 15-02 | 14-31 | 26-87 |
| 22:2 Stigmasteryl ester/22:3 Sitosteryl ester $[M+K]^+$ | 769-626 | 36-45 | 34-54 | 41-58 | 43-84 | 32-29 | 28-60 | 12-39 |
| 22:3 Campesteryl ester $[M+H]^+$ | 717-6549 | 1-49 | 3-06 | 0-43 | 0-42 | 2-03 | 2-13 | 0-89 |
| 22:3 Campesteryl ester $[M+K]^+$ | 755-6103 | 68-07 | 48-43 | 42-26 | 38-77 | 39-58 | 40-54 | 53-82 |
| 22:3 Campesteryl ester $[M+NH_4]^+$ | 734-6809 | 101-47 | 116-48 | 145-09 | 139-22 | 181-99 | 143-88 | 189-69 |
| 22:3 Stigmasteryl ester $[M+H]^+$ | 729-6549 | 16-09 | 11-35 | 9-13 | 10-41 | 16-49 | 18-48 | 24-16 |
| 22:3 Stigmasteryl ester $[M+K]^+$ | 767-6103 | 46-18 | 48-02 | 66-95 | 66-57 | 70-46 | 45-86 | 19-42 |
| 22:3 Stigmasteryl ester $[M+NH_4]^+$ | 746-6809 | 5-34 | 6-98 | 6-11 | 6-79 | | 6-63 | |
| Sitosteryl glucoside $[M+H]^+$ | 577-4468 | 10-19 | 9-80 | 7-42 | 16-87 | 20-20 | 17-00 | 20-66 |
| Sitosteryl glucoside $[M+K]^+$ | 615-4022 | 0-99 | 1-18 | 0-99 | 1-25 | 0-89 | 0-55 | 0-73 |
| Sitosteryl glucoside $[M+NH_4]^+$ | 594-4728 | 1-07 | 0-73 | 0-62 | 0-60 | 0-74 | 0-71 | 0-81 |
| Sphinganine-1-phosphocholine $[M+G]^+$ | 561-3786 | 1-23 | 1-28 | 2-26 | 2-43 | 1-72 | 1-31 | 0-81 |
| Sphingosine-1-phosphate $[M+G]^+$ | 474-2738 | 0-56 | 0-59 | 0-65 | | 0-78 | | 1-00 |
| Stigmasteryl glucoside $[M+H]^+$ | 575-4311 | 5-55 | 6-99 | 5-70 | 12-70 | 14-52 | 10-12 | 11-97 |

**Table S7.**
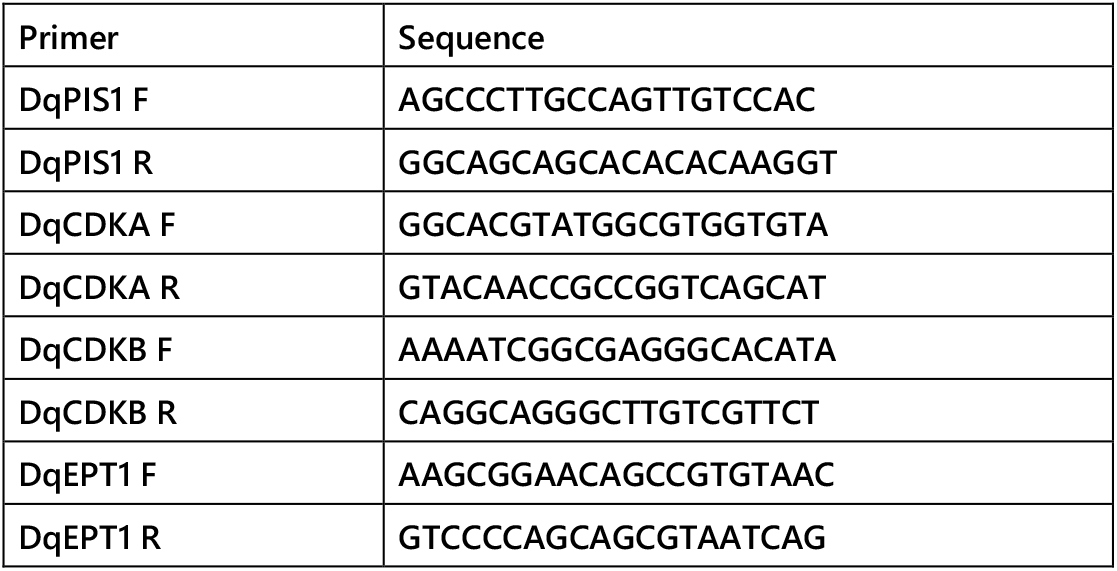
Sequences of primers used for qPCR amplification of homologues of cell cycle and lipid pathway genes. Each primer set was spanning at least one intron and amplified an amplicon of 150-200 nt.

## References

1. Hartwell, L. H. (2002) Nobel Lecture: Yeast and Cancer. Bioscience Reports 22, 373–394

2. Hunt, T. (2002) Nobel Lecture: Protein Synthesis, Proteolysis, and Cell Cycle Transitions. Bioscience Reports 22, 465–486

3. Nurse, P. (2002) Cyclin Dependent Kinases and Cell Cycle Control (Nobel Lecture). ChemBioChem 3, 596–603

4. Mironov, V., De Veylder, L., Van Montagu, M., and Inzé, D. (1999) Cyclin-Dependent Kinases and Cell Division in Plants—The Nexus. The Plant Cell 11, 509–521

5. Renaudin, J.-P., Doonan, J. H., Freeman, D., Hashimoto, J., Hirt, H., Inzé, D., Jacobs, T., Kouchi, H., Rouzé, P., Sauter, M., Savouré, A., Sorrell, D. A., Sundaresan, V., and Murray, J. A. H. (1996) Plant cyclins: a unified nomenclature for plant A-, B- and D-type cyclins based on sequence organization. Plant Mol Biol 32, 1003–1018

6. Bišová, K., Krylov, D. M., and Umen, J. G. (2005) Genome-Wide Annotation and Expression Profiling of Cell Cycle Regulatory Genes in *Chlamydomonas reinhardtii*. Plant Physiol 137, 475–491

7. Kono, K., Al-Zain, A., Schroeder, L., Nakanishi, M., and Ikui, A. E. (2016) Plasma membrane/cell wall perturbation activates a novel cell cycle checkpoint during G1 in Saccharomyces cerevisiae. Proceedings of the National Academy of Sciences 113, 6910–6915

8. Scotchman, E., Kume, K., Navarro, F. J., and Nurse, P. (2021) Identification of mutants with increased variation in cell size at onset of mitosis in fission yeast. Journal of Cell Science 134, jcs251769

9. Taheri-Araghi, S., Bradde, S., Sauls, John T., Hill, Norbert S., Levin, Petra A., Paulsson, J., Vergassola, M., and Jun, S. (2015) Cell-Size Control and Homeostasis in Bacteria. Current Biology 25, 385–391

10. Robert, L., Hoffmann, M., Krell, N., Aymerich, S., Robert, J., and Doumic, M. (2014) Division in Escherichia coli is triggered by a size-sensing rather than a timing mechanism. BMC Biology 12, 17

11. Umen, J. G. (2018) Sizing up the cell cycle: systems and quantitative approaches in Chlamydomonas. Current Opinion in Plant Biology 46, 96–103

12. Li, Y., Liu, D., López-Paz, C., Olson, B. J. S. C., and Umen, J. G. (2016) A new class of cyclin dependent kinase in Chlamydomonas is required for coupling cell size to cell division. eLife 5, e10767

13. Furse, S., Wienk, H., Boelens, R., de Kroon, A. I. P. M., and Killian, J. A. (2015) E. coli MG1655 modulates its phospholipid composition through the cell cycle. FEBS Letters 589, 2726–2730

14. Furse, S., Jakubec, M., Rise, F., Williams, H. E., Rees, C. E. D., and Halskau, O. (2017) Evidence that Listeria innocua modulates its membrane’s stored curvature elastic stress, but not fluidity, through the cell cycle. Scientific Reports 7, 8012

15. Koynova, R., and Caffrey, M. (1994) Phases and phase transitions of the hydrated phosphatidylethanolamines. Chemistry and Physics of Lipids 69, 1–34

16. Koynova, R., and Caffrey, M. (1998) Phases and phase transitions of the phosphatidylcholines. Biochimica et Biophysica Acta 1376, 91–145

17. Mulet, X., Templer, R. H., Woscholski, R., and Ces, O. (2008) Evidence That Phosphatidylinositol Promotes Curved Membrane Interfaces. Langmuir 24, 8443–8447

18. Furse, S., Brooks, N. J., Woscholski, R., Gaffney, P. R. J., and Templer, R. H. (2016) Pressure-dependent inverse bicontinuous cubic phase formation in a phosphatidylinositol 4-phosphate/phosphatidylcholine system. Chemical Data Collections 3-4, 15–20

19. Alley, S. H., Ces, O., Barahona, M., and Templer, R. H. (2008) X-ray diffraction measurement of the monolayer spontaneous curvature of dioleoylphosphatidylglycerol. Chemistry and Physics of Lipids 154, 64–67

20. Petrache, H. I., Tristram-Nagle, S., Gawrisch, K., Harries, D., Parsegian, V. A., and Nagle, J. F. (2004) Structure and Fluctuations of Charged Phosphatidylserine Bilayers in the Absence of Salt. Biophysical Journal 86, 1574–1586

21. Furse, S., and Shearman, G. C. (2018) Do lipids shape the eukaryotic cell cycle? Biochimica et Biophysica Acta 1863, 9–19

22. Zachleder, V., Bišová, K., and Vítová, M. (2016) The Cell Cycle of Microalgae. in The Physiology of Microalgae (Borowitzka, A. M., Beardall, J., and Raven, A. J. eds.), Springer International Publishing, Cham. pp 3–46

23. Bišová, K., and Zachleder, V. (2014) Cell-cycle regulation in green algae dividing by multiple fission. Journal of Experimental Botany 65, 2585–2602

24. Hlavova, M., Vitova, M., and Bisova, K. (2016) Synchronization of Green Algae by Light and Dark Regimes for Cell Cycle and Cell Division Studies. in Plant Cell Division: Methods and Protocols (Caillaud, M. C. ed.), Humana Press Inc, 999 Riverview Dr, Ste 208, Totowa, Nj 07512-1165 USA. pp 3–16

25. Zachleder, V., Bišová, K., Vítová, M., Kubín, S., and Hendrychová, J. (2002) Variety of cell cycle patterns in the alga Scenedesmus quadricauda (Chlorophyta) as revealed by application of illumination regimes and inhibitors. European Journal of Phycology 37, 361–371

26. Furse, S. (2017) Is phosphatidylglycerol essential for terrestrial life? Journal of Chemical Biology 10, 1–9

27. Furse, S., and de Kroon, A. I. P. M. (2015) Phosphatidylcholine’s functions beyond that of a membrane brick. Molecular Membrane Biology 32, 117–119

28. Furse, S., Fernandez-Twinn, D., Jenkins, B., Meek, C. L., Williams, H. E., Smith, G. C. S., Charnock-Jones, D. S., Ozanne, S. E., and Koulman, A. (2020) A high throughput platform for detailed lipidomic analysis of a range of mouse and human tissues. Anal Bioanal Chem 412, 2851–2862

29. Furse, S., Williams, H. E. L., Watkins, A. J., Virtue, S., Vidal-Puig, A., Amarsi, R., Charalambous, M., and Koulman, A. (2021) A pipeline for making 31P NMR accessible for small- and large-scale lipidomics studies In prep for Anal and Bioanal Chem

30. Furse, S., Liddell, S., Ortori, C. A., Williams, H., Neylon, D. C., Scott, D. J., Barrett, D. A., and Gray, D. A. (2013) The lipidome and proteome of oil bodies from Helianthus annuus (common sunflower). Journal of chemical biology 6, 63–76

31. Felde, R., and Spiteller, G. (1994) Search for plasmalogens in plants. Chemistry and Physics of Lipids 71, 109–113

32. Zhou, Y., Peisker, H., and Dörmann, P. (2016) Molecular species composition of plant cardiolipin determined by liquid chromatography mass spectrometry. Journal of Lipid Research 57, 1308–1321

33. Atilla-Gokcumen, G. E., Muro, E., Relat-Goberna, J., Sasse, S., Bedigian, A., Coughlin, Margaret L., Garcia-Manyes, S., and Eggert, Ulrike S. (2014) Dividing Cells Regulate Their Lipid Composition and Localization. Cell 156, 428–439

34. Blouin, A., Lavezzi, T., and Moore, T. S. (2003) Membrane lipid biosynthesis in Chlamydomonas reinhardtii. Partial characterization of CDP-diacylglycerol: myo-inositol 3-phosphatidyltransferase. Plant Physiology and Biochemistry 41, 11–16

35. Tyler, A. I. I., Barriga, H. M. G., Parsons, E. S., McCarthy, N. L. C., Ces, O., Law, R. V., Seddon, J. M., and Brooks, N. J. (2015) Electrostatic swelling of bicontinuous cubic lipid phases. Soft Matter 11, 3279–3286

36. Dawaliby, R., Trubbia, C., Delporte, C., Noyon, C., Ruysschaert, J.-M., Van Antwerpen, P., and Govaerts, C. (2016) Phosphatidylethanolamine Is a Key Regulator of Membrane Fluidity in Eukaryotic Cells. Journal of Biological Chemistry 291, 3658–3667

37. Emoto, K., and Umeda, M. (2000) An Essential Role for a Membrane Lipid in Cytokinesis: Regulation of Contractile Ring Disassembly by Redistribution of Phosphatidylethanolamine. The Journal of Cell Biology 149, 1215–1224

38. Seddon, J. M., and Templer, R. H. (1995) Polymorphism of Lipid-Water Systems. in The Handbook of Biological Physics (Lipowsky, R., and Sackman, E. eds.), Elsevier Science. pp

39. Peng, A., Pisal, D. S., Doty, A., and Balu-Iyer, S. V. (2012) Phosphatidylinositol induces fluid phase formation and packing defects in phosphatidylcholine model membranes. Chemistry and Physics of Lipids 165, 15–22

40. Emoto, K., Inadome, H., Kanaho, Y., Narumiya, S., and Umeda, M. (2005) Local Change in Phospholipid Composition at the Cleavage Furrow Is Essential for Completion of Cytokinesis. Journal of Biological Chemistry 280, 37901–37907

41. Emoto, K., Kobayashi, T., Yamaji, A., Aizawa, H., Yahara, I., Inoue, K., and Umeda, M. (1996) Redistribution of phosphatidylethanolamine at the cleavage furrow of dividing cells during cytokinesis. Proceedings of the National Academy of Sciences 93, 12867–12872

42. Kobayashi, K. (2016) Role of membrane glycerolipids in photosynthesis, thylakoid biogenesis and chloroplast development. Journal of Plant Research 129, 565–580

43. Schleiff, E., Tien, R., Salomon, M., and Soll, J. (2001) Lipid composition of outer leaflet of chloroplast outer envelope determines topology of OEP7. Molecular Biology of the Cell 12, 4090–4102

44. Hung, C.-H., Endo, K., Kobayashi, K., Nakamura, Y., and Wada, H. (2015) Characterization of Chlamydomonas reinhardtii phosphatidylglycerophosphate synthase in Synechocystis sp. PCC 6803. Frontiers in Microbiology 6

45. Kobayashi, K., Endo, K., and Wada, H. (2016) Multiple Impacts of Loss of Plastidic Phosphatidylglycerol Biosynthesis on Photosynthesis during Seedling Growth of Arabidopsis. Frontiers in Plant Science 7

46. Tanoue, R., Kobayashi, M., Katayama, K., Nagata, N., and Wada, H. (2014) Phosphatidylglycerol biosynthesis is required for the development of embryos and normal membrane structures of chloroplasts and mitochondria in Arabidopsis. FEBS Letters 588, 1680–1685

47. Cremonini, M. A., Laghi, L., and Placucci, G. (2004) Investigation of commercial lecithin by P-31 NMR in a ternary CUBO solvent. Journal of the Science of Food and Agriculture 84, 786–790

48. Burr, R., Stewart, E. V., Shao, W., Zhao, S., Hannibal-Bach, H. K., Ejsing, C. S., and Espenshade, P. J. (2016) Mga2 Transcription Factor Regulates an Oxygen-responsive Lipid Homeostasis Pathway in Fission Yeast*. Journal of Biological Chemistry 291, 12171–12183

49. Burr, R., Stewart, E. V., and Espenshade, P. J. (2017) Coordinate Regulation of Yeast Sterol Regulatory Element-binding Protein (SREBP) and Mga2 Transcription Factors*. Journal of Biological Chemistry 292, 5311–5324

50. Wanjie, S. W., Welti, R., Moreau, R. A., and Chapman, K. D. (2005) Identification and quantification of glycerolipids in cotton fibers: Reconciliation with metabolic pathway predictions from DNA databases. Lipids 40, 773–785

51. Maatta, S., Scheu, B., Roth, M., Tamura, P., Li, M., Williams, T., Wang, X., and Welti, R. (2012) Levels of Arabidopsis thaliana Leaf Phosphatidic Acids, Phosphatidylserines, and Most Trienoate-Containing Polar Lipid Molecular Species Increase during the Dark Period of the Diurnal Cycle. Frontiers in Plant Science 3

52. Jüppner, J., Mubeen, U., Leisse, A., Caldana, C., Brust, H., Steup, M., Herrmann, M., Steinhauser, D., and Giavalisco, P. (2017) Dynamics of lipids and metabolites during the cell cycle of Chlamydomonas reinhardtii. The Plant Journal 92, 331–343

53. Giroud, C., Gerber, A., and Eichenberger, W. (1988) Lipids of Chlamydomonas reinhardtii. Analysis of Molecular Species and Intracellular Site(s) of Biosynthesis. Plant and Cell Physiology 29, 587–595

54. Liu, G.-J., Xiao, G.-H., Liu, N.-J., Liu, D., Chen, P.-S., Qin, Y.-M., and Zhu, Y.-X. (2015) Targeted Lipidomics Studies Reveal that Linolenic Acid Promotes Cotton Fiber Elongation by Activating Phosphatidylinositol and Phosphatidylinositol Monophosphate Biosynthesis. Molecular Plant 8, 911–921

55. Kurat, C. F., Wolinski, H., Petschnigg, J., Kaluarachchi, S., Andrews, B., Natter, K., and Kohlwein, S. D. (2009) Cdk1/Cdc28-Dependent Activation of the Major Triacylglycerol Lipase Tgl4 in Yeast Links Lipolysis to Cell-Cycle Progression. Molecular Cell 33, 53–63

56. Radakovits, R., Jinkerson, R. E., Darzins, A., and Posewitz, M. C. (2010) Genetic Engineering of Algae for Enhanced Biofuel Production. Eukaryotic Cell 9, 486–501

57. Hu, Q., Sommerfeld, M., Jarvis, E., Ghirardi, M., Posewitz, M., Seibert, M., and Darzins, A. (2008) Microalgal triacylglycerols as feedstocks for biofuel production: perspectives and advances. The Plant Journal 54, 621–639

58. Vítová, M., Bišová, K., Hlavová, M., Zachleder, V., Rucki, M., and Čížková, M. (2011) Glutathione peroxidase activity in the selenium-treated alga Scenedesmus quadricauda. Aquatic Toxicology 102, 87–94

59. Furse, S., and Koulman, A. (2019) The Lipid and Glyceride Profiles of Infant Formula Differ by Manufacturer, Region and Date Sold. Nutrients 11, 1122

60. Furse, S., Watkins, A. J., and Koulman, A. (2020) Extraction of Lipids from Liquid Biological Samples for High-Throughput Lipidomics. Molecules 25, 3192

61. Furse, S., and Koulman, A. (2020) Lipid extraction from dried blood spots and dried milk spots for untargeted high throughput lipidomics. Molecular Omics

62. Kochen, M. A., Chambers, M. C., Holman, J. D., Nesvizhskii, A. I., Weintraub, S. T., Belisle, J. T., Islam, M. N., Griss, J., and Tabb, D. L. (2016) Greazy: Open-Source Software for Automated Phospholipid Tandem Mass Spectrometry Identification. Analytical Chemistry 88, 5733–5741

63. Harshfield, E. L., Koulman, A., Ziemek, D., Marney, L., Fauman, E. B., Paul, D. S., Stacey, D., Rasheed, A., Lee, J.-J., Shah, N., Jabeen, S., Imran, A., Abbas, S., Hina, Z., Qamar, N., Mallick, N. H., Yaqoob, Z., Saghir, T., Rizvi, S. N. H., Memon, A., Rasheed, S. Z., Memon, F.-u.-R., Qureshi, I. H., Ishaq, M., Frossard, P., Danesh, J., Saleheen, D., Butterworth, A. S., Wood, A. M., and Griffin, J. L. (2019) An Unbiased Lipid Phenotyping Approach To Study the Genetic Determinants of Lipids and Their Association with Coronary Heart Disease Risk Factors. Journal of Proteome Research 18, 2397–2410

64. Hlavová, M., í?ková, M., Vítová, M., Bi?ová, K. i., and Zachleder, V. (2011) DNA Damage during G2 Phase Does Not Affect Cell Cycle Progression of the Green Alga *Scenedesmus quadricauda*. PLoS ONE 6, e19626

65. Laemmli, U. K. (1970) Cleavage of Structural Proteins during the Assembly of the Head of Bacteriophage T4. Nature 227, 680

66. Towbin, H., Staehelin, T., and Gordon, J. (1979) Electrophoretic transfer of proteins from polyacrylamide gels to nitrocellulose sheets: procedure and some applications. Proceedings of the National Academy of Sciences 76, 4350–4354

67. Vítová, M., Hendrychová, J., Čížková, M., Cepák, V., Umen, J. G., Zachleder, V., and Bišová, K. (2008) Accumulation, Activity and Localization of Cell Cycle Regulatory Proteins and the Chloroplast Division Protein FtsZ in the Alga Scenedesmus quadricauda under Inhibition of Nuclear DNA Replication. Plant and Cell Physiology 49, 1805–1817

68. Langan, T. A., Gautier, J., Lohka, M., Hollingsworth, R., Moreno, S., Nurse, P., Maller, J., and Sclafani, R. A. (1989) Mammalian growth-associated H1 histone kinase: a homolog of cdc2+/CDC28 protein kinases controlling mitotic entry in yeast and frog cells. Mol. Cell. Biol. 9, 3860–3868

69. Bisova, K., Krylov, D. M., and Umen, J. G. (2005) Genome-wide annotation and expression profiling of cell cycle regulatory genes in *Chlamydomonas reinhardtii*. Plant Physiology 137, 1–17

70. Altschul, S. F., Gish, W., Miller, W., Myers, E. W., and Lipman, D. J. (1990) Basic local alignment search tool. Journal of Molecular Biology 215, 403–410

71. Fan, J., Ning, K., Zeng, X., Luo, Y., Wang, D., Hu, J., Li, J., Xu, H., Huang, J., Wan, M., Wang, W., Zhang, D., Shen, G., Run, C., Liao, J., Fang, L., Huang, S., Jing, X., Su, X., Wang, A., Bai, L., Hu, Z., Xu, J., and Li, Y. (2015) Genomic Foundation of Starch-to-Lipid Switch in Oleaginous *Chlorella* spp. Plant Physiol 169, 2444–2461

72. Lu, S., Wang, J., Ma, Q., Yang, J., Li, X., and Yuan, Y.-J. (2013) Phospholipid Metabolism in an Industry Microalga Chlorella sorokiniana: The Impact of Inoculum Sizes. PLOS ONE 8, e70827

73. Khozin-Goldberg, I. (2016) Lipid Metabolism in Microalgae. in The Physiology of Microalgae (Borowitzka, M. A., Beardall, J., and Raven, J. A. eds.), Springer International Publishing, Cham. pp 413–484

74. Ledesma-Amaro, R., Dulermo, R., Niehus, X., and Nicaud, J.-M. (2016) Combining metabolic engineering and process optimization to improve production and secretion of fatty acids. Metabolic Engineering 38, 38–46

